# Functional characterization of the thrombospondin-related paralogous proteins rhoptry discharge factor 1 and 2 unveils phenotypic plasticity in *Toxoplasma gondii* rhoptry exocytosis

**DOI:** 10.1101/2022.03.02.482699

**Authors:** Alessia Possenti, Manlio Di Cristina, Chiara Nicastro, Matteo Lunghi, Valeria Messina, Federica Piro, Lorenzo Tramontana, Simona Cherchi, Mario Falchi, Lucia Bertuccini, Furio Spano

## Abstract

To gain access to the intracellular cytoplasmic niche essential for their growth and replication, apicomplexan parasites such as *Toxoplasma gondii* rely on the timely secretion of two types of apical organelles named micronemes and rhoptries. Rhoptry proteins are key to host cell invasion and remodelling, however, the molecular mechanisms underlying the thight control of rhoptry discharge are poorly understood. Here, we report the identification and functional characterization of two novel *T. gondii* thrombospondin-related proteins implicated in rhoptry exocytosis. The two proteins, already annotated as MIC15 and MIC14, were renamed rhoptry discharge factor 1 (RDF1) and rhoptry discharge factor 2 (RDF2) and found to be exclusive of the Coccidia class of apicomplexan parasites. Furthermore, they were shown to have a paralogous relationship and share a C-terminal transmembrane domain followed by a short cytoplasmic tail. Immunofluorescence analysis of *T. gondii* tachyzoites revealed that RDF1 presents a diffuse cytoplasmic localization not reminiscent of any know subcellular compartment, whereas RDF2 was not detected. Using a conditional knockout approach, we demonstrated that RDF1 loss caused a marked growth defect. The lack of the protein did not affect parasite gliding motility, host cell attachment, replication and egress, whereas invasion was dramatically reduced. Notably, while RDF1 depletion did not result in altered microneme exocytosis, rhoptry discharge was found to be heavily impaired. Interestingly, rhoptry secretion was partially reversed by spontaneous upregulation of the *RDF2* gene in knockdown parasites grown under constant *RDF1* repression. Collectively, our results identify RDF1 and RDF2 as additional key players in the pathway controlling rhoptry discharge. Furthermore, this study unveils a new example of compensatory mechanism contributing to phenotypic plasticity in *T. gondii*.

## INTRODUCTION

The protozoan *Toxoplasma gondii* is an obligate intracellular parasite of the phylum Apicomplexa, which includes several other pathogens belonging to genera of great medical or veterinary relevance such as *Plasmodium, Cryptosporidium, Eimeria* and *Theileria*. Toxoplasmosis affects humans and farm animals worldwide, causing life-threatening infections in immunocompromised subjects, and abortion or serious birth defects if congenitally transmitted to the fetus. Seroprevalence data indicate that one third of the human population has been exposed to *T. gondii* and is likely chronically infected.

The symptomatic manifestations of acute toxoplasmosis depend on tachyzoites, which couple efficient dissemination to any body district of the infected host to a fast replicating phenotype, leading to tissue damage. Before the onset of the host immune response, *T. gondii* tachyzoites undergo multiple lytic cycles marked by four sequential steps, i.e., host cell attachment, active invasion, intravacuolar replication and lytic egress. During the motile phases of the lytic cycle, a major role is played by the apical complex, a sophisticated system of cytoskeletal and vesicular elements distinctive of the invasive stages of Apicomplexa. The cytoskeletal component includes the conoid, a peculiar structure made of spirally arranged tubulin fibers that is reversibly extruded during parasite gliding motility, invasion and egress (Dos Santos Pacheco et al., 2020). Membrane bound elements include an array of non secretory apical vesicles [2,3] and two types of secretory organelles, called micronemes and rhoptries, that discharge their content in a sequential and tightly regulated fashion, orchestrating the transition from the extracellular to the intravacuolar environment. Upon microneme secretion (Dubois et al., 2019), various transmembrane adhesins are apically exposed on the parasite plasma membrane (PM) and act as a link between extracellular ligands and the subpellicular acto-myosin motor of the parasite, playing a key role in substrate-dependent gliding motility, host cell attachment and invasion. The club-shaped rhoptries consist of an elongated neck passing through the conoid and a posterior bulb (Ben Chaabene et al., 2021). These two organellar districts contain specific sets of proteins called RONs and ROPs, respectively, which are released from the rhoptries upon host cell attachment and microneme secretion. When a complex of specific RON proteins is inserted into the host cell membrane and recognized by the micronemal protein apical membrane antigen-1 (AMA1) displayed on the tachyzoite surface, a ring-like moving junction is produced (Alexander et al., 2005; Besteiro et al., 2009), that allows parasite firm attachment to and traversal of the host cell PM. Following RONs secretion, ROP proteins are injected into the host cell cytoplasm within clusters of small vesicles called evacuoles (Håkansson et al., 2001) and contribute to parasitophorous vacuole formation, modulation of host cell transcr iption and virulence (Ben Chaabene et al., 2021).

In the complex interplay between *T. gondii* and the host cell, a prominent role is played by modular proteins containing adhesive amino acid motifs conserved in metazoan organisms, such as the PAN/Apple, epidermal growth factor-like, von Willebrand factor type A (vWA) and thrombospondin type 1 (TSP1) domains. This latter motif of approximately 60 amino acids is able to bind carbohydrate and protein substrates and was identified in the extracellular matrix glycoprotein thrombospondin-1 and in dozens of other human proteins involved in cell-to-cell an cell-to-matrix interactions (Adams and Tucker, 2000). In apicomplexan parasites the TSP1 domain is present in several secretory proteins fulfilling important functions. In *Plasmodium* spp., stage-specific members of the TSP1 domain superfamily are implicated in processes as diverse as gliding motility, cell binding, invasion, egress and cell traversal and include the proteins CSP (Nardin et al., 1982), TRAP (Robson et al., 1988), CTRP (Trottein et al., 1995), MTRAP (Baum et al., 2006), PTRAMP (Thompson et al., 2004), TLP (Moreira et al., 2008), TREP (Combe et al., 2009) and TRP1 (Klug and Frischknecht, 2017). Notably, except the surface GPI-anchored CSP, all these proteins have a type I transmembrane (TM) topology and share with the prototypic TRAP an extracellular adhesive domain, a C-terminal TM region and a cytoplasmic tail that interacts with the force-generating acto-myosin system of the parasite. In *T. gondii*, MIC2 (Huynh et al., 2003), MIC12 (Opitz et al., 2002) and MIC16 (Sheiner et al., 2010) are the only TSP1 domain-containing proteins characterized so far. They share with *Plasmodium* members of the TRAP family the overall structural organization and the localization to the micronemes. The most extensively characterized is the TRAP functional ortholog MIC2, which plays a crucial role in tachyzoite host cell attachment (Huynh and Carruthers, 2006) and is involved in gliding motility and host cell egress (Gras et al., 2017). MIC2 contains a single vWA domain and six tandemly arrayed TSP1 motifs, one of which directly mediates the formation of a complex with the soluble MIC2-associated protein (M2AP) essential for microneme targeting (Huynh et al., 2015).

In the present work we explored the genome of *T. gondii* to search for additional and functionally relevant proteins containing the TSP1 domain. We identified nine new members of this protein subclass, the majority of which are expected to play important or essential roles based on a genome wide CRISPR-Cas9 dataset (Sidik et al., 2016). We report herein on the functional characterization of two paralogous TSP1 domain-containing proteins specific of Coccidia. Despite their TRAP-like structural organization, the two proteins do not traffic to the micronemes but present a dispersed cytoplasmic localization resembling that of proteins recently implicated in rhoptry secretion [2,25]. Using reverse genetics, we demonstrate that the two proteins represent novel rhoptry discharge factors and that their redundancy confers to the parasite a further degree of phenotypic plasticity capable to preserve an essential function related to host cell invasion.

## MATERIALS AND METHODS

### Parasites

*Toxoplasma gondii* tachyzoites of strains RHΔ*ku80* and TATiΔ*ku80* (Sheiner et al., 2011) were grown in human foreskin fibroblasts (HFF) monolayers in Dulbecco’s modified Eagle’s medium (DMEM) with GlutaMAX (Gibco) supplemented with 10% Nu-Serum (Gibco), 10 mM HEPES (N-2-hydroxy-ethylpiperazine-N’-2-ethanesulfonic acid) and penicillin/stretptomycin at 37°C in a humid 5% CO_2_ atmosphere. Tachyzoites from actively lysing HFF cultures were recovered by passage through a 27-gauge syringe needle and purified by a 3-□m membrane filter.

### Generation of parasite strains

Parasite transfections were carried out as previously described (Afonso et al., 2017). Primers used in this study are listed in S1 Table.

To generate the vector *pMIC15-iKD-TRE/S4*, genomic fragments of 1269 and 1176 bp located upstream and downstream of the MIC15 promoter of *T. gondii* RH strain were amplified with primers P44/P45 and P46/P47, respectively. The TRE/S4 cassette for the replacement of the *MIC15* promoter was amplified from plasmid *pDt7S4* (van Dooren et al., 2008) using the primer pair P48/P49. The upstream sequence, the TRE/S4 cassette and the downstream sequence were cloned sequentially in the *Eco*RV-*Apa*I, *Bam*H1-*Spe*I and *Spe*I-*Sac*II restriction sites of a modified version of *pTUB5-DHFR* (Afonso et al., 2017), in which an additional *Eco*RV restriction site was added upstream of the resistance marker.

The complementing vector *pMIC15-comp-CAT* was generated by amplifying the *MIC15* cDNA coding region from strain RHΔ*ku80* using the primers P13 and P14. The 8960 bp amplicon was cut with the restriction enzymes *Xba*I and *Pac*I and cloned in the corresponding restriction sites upstream of the 3’UTR of the *GRA2* gene, already present in the acceptor plasmid. A region of 2065 bp encompassing the *MIC15* promoter was then amplified from RHΔ*ku80* genomic DNA with primers P15/P16 and cloned between the *Not*I and *Xba*I restriction sites upstream of the *MIC15* coding region. Finally, the *TUB-CAT* resistance cassette was amplified from the *pTUB5-CAT* plasmid using the primers P17 and P18 and inserted in the *Asc*I and *Not*I restriction sites upstream of the *MIC15* minigene, obtaining plasmid *pMIC15-comp-CAT*. The vector was sequenced to rule out the occurrence of nucleotide substitutions. The complementing vector was then linearized at the *Xcm*I restriction site and transfected in the MIC15-iKD strain for integration by single crossing over upstream of the tetracycline-controlled *MIC15* locus. The correct integration of the complementing plasmid was assessed by PCR using the primer pairs P25/P26 and P27/P28 (**Supplementary Figure 5**).

To produce the knockout construct pMIC14KO, a genomic region of approximately 1.1 kb lying upstream of the *MIC14* promoter and one spanning exons 4 and 5 of *MIC14*, were amplified from RHΔ*ku80* genomic DNA with the primer pairs P9/10 and P11/P12, respectively. The two amplicons were cloned in the *pTub5/CAT* plasmid (Soldati and Boothroyd, 1993) 5’ and 3’ to the *CAT* gene, using the restriction sites *Apa*I/*Hin*dIII and *Spe*I/*Not*I, respectively. The *Not*I linearized plasmid was introduced into the MIC15-iKD/MIC14^R^ strain to replace the promoter and the 5’-terminal exons with the *CAT* cassette by double homologous recombination. Transfected parasites were selected with 20 μM chloramphenicol and cloned, obtaining strain MIC15-iKD/MIC14^R^KO/MIC14KO. The occurrence of the correct recombination event was tested by PCR (**Figure 7D**).

To generate strain MIC15-smMYC the *MIC15* gene was endogenously tagged by inserting the sequence encoding the spaghetti monster-c-myc (smMYC) protein in the second exon of *MIC15*. The sequence encoding the smMYC tag was amplified from plasmid *pLIC-SMGFP_MYC* (Hortua Triana et al., 2018) with primers P55 and P56, containing at the 5’ end 40 bp sequences lying upstream and downstream of the selected *MIC15* insertion point, respectively. Insertion of the smMYC epitope between amino acids 29 and 30 was obtained by co-transfecting RHΔ*ku80* tachyzoites with the amplified smMYC repair template and a CRISPR/Cas9 vector encoding the Cas9 endonuclease, and containing the bleomycin resistance gene and the *MIC15*-specific gRNA sequence 5’-GUAACUUGUCAACUGCAGUU-3’. Twenty-four hours post transfection parasites were mechanically extruded from HFF and treated for 4 h with 50 μg/ml of bleomycin at 37°C to enrich for parasites harboring the CRISPR/Cas9 vector. Bleomycin-treated parasites were allowed to expand in HFF monolayer and then cloned by limiting dilution into 96-well plates. Identification of clones carrying the correct integration of the smMYC tag was carried out as described (Piro et al., 2020) using primer pairs P57/59 or P58/P47.

Insertion of the Ty epitope upstream of the MIC15 TM region (between amino acids 2688 and 2689) was obtained by co-transfecting RHΔ*ku80* tachyzoites with a Ty repair template along with a CRISPR/Cas9 vector encoding the Cas9 endonuclease, and containing the *MIC15*-specific gRNA sequence 5’-AUCGAUCUUUCCGAAGCUUC-3’ and the bleomycin resistance cassette. The repair template was generated by annealing the complementary 110 bp long oligonucleotides P29 and P30 (Di Cristina and Carruthers, 2018), and consisted of the Ty coding sequence flanked by 40 bp long genomic regions lying upstream and downstream of the selected *MIC15* insertion point. Following the bleomycin selection procedure described above for the MIC15-smMYC strain, individual parasite clones were screened for the correct integration of the TY tag using primer pairs P1/P2 or P3/P4.

Strain MIC15-2xHA was generated by endogenously tagging the 3’ end of *MIC15* in RHΔ*ku80* tachyzoites by single homologous recombination, using plasmid *pMIC15-2xHA-DHFR*. A DNA sequence of □1.3 kb spanning exons 41 and 42 of *MIC15* was amplified from the RHΔ*ku80* genome with primers P60 and P61, containing the restriction sites *Eco*RV and *Pac*I, respectively. In addition, primer P61 encoded two tandemly repeated HA epitopes followed by a stop codon. A second genomic fragment encompassing the 3’UTR of the *DHFR* gene was amplified with primers P62 and P63, which carried the restriction sites *Pac*I and *Apa*I, respectively. Exploiting the common *Pac*I restriction site, the two amplicons were tandemly cloned in the *Eco*RV and *Apa*I restriction sites of a modified version of *pTUB5-DHFR* (Afonso et al., 2017). Following transfection with *Bst*EII-linearized plasmid *pMIC15-2xHA-DHFR* and pyrimethamine selection, parasite clones were screened for the C-terminal insertion of 2xHA epitopes using primers P75 and P63.

Strain MIC15-iKD/MIC14^R^-3xTy was generated by endogenously tagging the 3’ end of *MIC14* with three consecutive Ty epitopes in MIC15-iKD/MIC14^R^ parasites. A genomic sequence of □1.0 kb spanning exons 42 and 43 of *MIC14* was amplified with primers P70 and P71 harboring the restriction sites *Not*I and *Sph*I, respectively. A second PCR product, encompassing the sequence coding for 3xTy epitopes and the 3’UTR of SAG1, was obtained from plasmid *DGK2-3xTy* (gently donated by Dominique Soldati-Favre, University of Geneva) using primers P68 and P69, which carried the restriction sites *Sph*I and *Bam*HI, respectively. Exploiting the common *Sph*I restriction site, the two amplicons were tandemly cloned in the *Not*I and *Bam*H1 sites of *pTUB5-CAT*. The resulting plasmid *pMIC14-3xTy* was linearized at the *Nhe*I restriction site and introduced in strain MIC15-iKD/MIC14^R^ by single homologous recombination. Individual parasite clones were screened for the correct recombination event by PCR using primers P72 and P69.

Strain TUB-MIC14^R^-3xTy was obtained in MIC15-iKD/MIC14^R^-3xTy parasites by a single homologous recombination event that replaced the *MIC14* endogenous promoter with that of the β-tubulin gene. Using primers P73 and P74, a DNA sequence of □1200 bp spanning *MIC14* from the translation initiation codon to the second exon was amplified from the RHΔ*ku80* genome and cloned downstream of the TUB promoter of plasmid pGRA1-NAT1_TUB (Van Tam et al., 2006), exploiting the restriction sites *Eco*RI and *Xba*I. Following transfection with the resulting plasmid *pTUB/MIC14-3xTY-NAT* linearized at the *Hpa*I restriction site and two rounds of selection of extracellular tachyzoites with 500 μg/ml of nourseothricin, parasites were cloned by limiting dilution and screened with the diagnostic primers P43 and P73.

### Analysis of full-length cDNAs

The full length cDNAs of *MIC14* and *MIC15* were reconstructed by combining the sequences of polyadenylated cDNA clones isolated from a tachyzoite library, with those of overlapping RT-PCR and 5’ RACE products. A λZAPII cDNA library of strain RH (NIH AIDS Research and Reference Reagent Program) was screened by limiting dilution (Israel, 1993) using the oligonucleotide pairs P5/P6 for *MIC14* and P7/P8 for *MIC15*. The 5’ end regions of *MIC14* and *MIC15* were amplified using the SMART RACE cDNA Amplification Kit (Clontech BD) using total or poly(A)^+^tachyzoite RNA (strain RH) as starting material. For amplification of the *MIC14* 5’ end, 1.5 μg of poly(A)^+^ tachyzoite RNA was reverse transcribed with random examers prior to PCR amplification with the *MIC14* primer P43 and the nested universal primer included in the kit. For amplification of the 5’ end of *MIC15*, 1.5 μg of total RNA was reverse transcribed with primer P39 prior to PCR amplication of the RT product with the *MIC15* nested primer P40 and the nested universal primer provided with the kit. The *MIC14* cDNA region encompassing the full length gene from exon 2 to exon 29 was amplified from reverse transcribed poly(A)^+^ RNA using the primer pair P41/P42. Individual 5’ RACE and RT-PCR products were cloned into the pCR2.1 TA cloning vector (Invitrogen) and sequenced.

### Antibodies

Antibodies directed against *T. gondii* proteins or epitope tags used in this study and the relative dilutions used in Western blot and immunofluorescence analyses are described in S2 Table. Rabbit polyclonal antibodies MIC15Nt and MIC15Ct were raised against recombinant fragments of MIC15 spanning amino acids 928-1460 and 2494-2706, respectively. Mouse polyclonal serum T148 was raised against the MIC14 region encompassing amino acids 1761-1996. Recombinant protein fragments were produced in *Escherichia coli* (strain M15) as fusion products with an N-terminal tag of six histidines using plasmid pQE30 (Qiagen) and purified from total bacterial lysates by nickel affinity chromatography. The anti-MIC15 monoclonal antibody 22E6 was produced as previously described (Possenti et al., 2010) using as immunogen the same protein fragment employed to obtain rabbit serum MIC15Nt.

### Quantitative real-time PCR

Total tachyzoite RNA was extracted using the RNeasy Plus Mini kit (Qiagen) and treated with Turbo DNA free kit (Ambion) to remove residual genomic DNA. Following RNA quantification with the Qubit RNA HS assay kit (Invitrogen) and inspection of RNA integrity by agarose gel electrophoresis, 0.5-1 μg of total RNA was reverse transcribed using Protoscript II Reverse Transcriptase (New England Biolabs). Quantitative real-time PCR was carried out using the Light-Cycler 480 SYBR Green I Master (Roche). Each 20 μl reaction contained 10 μl of 2x SYBER Green I Master mix, 20 pmol of each primer and 4 μl of cDNA (diluted 1:20). One pre-incubation step (5 min, 95°C) was followed by 45 cycles of denaturation (10 sec, 95°C), annealing (10 sec, 58°C) and elongation (10 sec, 72°C) performed on a LyghtCycler 96 Instrument (Roche). *MIC14* and *MIC15* transcripts were amplified with the primer pairs P64/P65 and P66/P67. Relative levels of *MIC14* and *MIC15* mRNAs with respect to the β-tubulin mRNA internal control were calculated using the 2-^ΔΔCt^ method. All samples were analyzed in triplicate using both biological and technical replicates.

### Western blot analysis

Excreted-secreted antigens (ESA) were prepared as described below in the paragraph on microneme secretion. Total protein lysates were obtained by incubating purified *T. gondii* tachyzoites in RIPA buffer (150 mM NaCl, 1% NP-40, 0.5% sodium deoxycholate, 0.1% SDS, 50 mM Tris–HCl pH 8.0) for 20 min on ice and removing insoluble material by centrifugation at 3000 x g for 20 min at 4°C.. Parasite proteins were reduced with 200 □M dithiotreitol (DTT) and separated on 4-12% NuPAGE Bis-Tris gels (Thermo Fisher Scientific). Following wet protein transfer to nitrocellulose, the filters were blocked for 1 hr with 5% skimmed milk (Sigma-Aldrich) in 2x Tris Buffered Saline/Tween 20 (20 mM Tris–HCl pH 8.0, 300 mM NaCl, 0.1% Tween 20) and subsequently incubated for 1 hr with primary antibodies at dilutions reported in S2 Table. Reactive bands were visualised using goat anti-mouse IgG or goat anti-rabbit IgG secondary antibodies conjugated to horseradish peroxidase (Invitrogen) and the LiteAblot Plus Enhanced Chemiluminescent Substrate (EuroClone).

### Immunofluorescence analysis

Indirect immunofluorescence assays were performed on extracellular *T. gondii* tachyzoites or parasite-infected HFF monolayers grown for 24-30 h at 37°C on 12 mm glass coverslips or in 8-well chamber slides. The samples were fixed for 10 min with 4% formaldehyde, quenched with 0.1 M glycine/PBS, permeabilized with 0.5% Triton X-100 if required and blocked for 1 h with 2% fetal bovine serum in PBS. Incubation with primary antibodies diluted in 2% fetal bovine serum in PBS was carried out for 1 hr at dilutions reported in S2 Table. Bound primary antibodies were detected using 1:500-1:2000 dilutions of goat anti-mouse or goat anti-rabbit IgG secondary antibodies conjugated with Alexa Fluor 488 or Alexa Fluor 594 (Invitrogen). Tachyzoite and HFF nuclei were labelled with DAPI (4’,6’-diamidino-2-phenylindole). Samples were mounted in SlowFade Antifade reagent (Invitrogen) and stored at 4°C in the dark. Confocal images were taken by a Zeiss LSM 980 confocal microscope (Zeiss, Germany), using a planapo objective 60x oil A.N. 1,42. Images recorded have an optical thickness of 0.4 □m. Other images were taken by an Axioplan 2 epifluorescence microscope (Zeiss, Germany) using a 100 x oil immersion objective and an Axiocam digital camera using the Zeiss Axiovision 4.7 software.

### Microneme secretion assay

To obtain the excreted-secreted antigen (ESA) fraction, freshly egressed tachyzoites grown ± Atc for 60 h were resuspended in DMEM/10 mM HEPES at a concentration of 2×10^9^ parasites/ml and incubated for 20 min at 37°C with or without 1% ethanol to stimulate microneme secretion (Carruthers et al., 1999). Following parasite centrifugation at 1000 g for 10 min at 4°C, the pellet fraction was collected and extracted with RIPA buffer, the supernatant was centrifuged at 2000 g for 5 min at 4°C to remove residual tachyzoites. The supernatant, representing the ESA fraction, was concentrated using Centricon 20 Plus concentrators (Millipore) and stored at −80°C for further use. Pellet and ESA protein amounts equivalent to 1.2×10^7^ or 4×10^7^ parasites, respectively, were analyzed by SDS gel electrophoresis on NuPAGE™ 4-12% Bis-Tris gels (Thermo Fisher Scientific) and blotted.

### Plaque assay

Freshly egressed tachyzoites cultivated ± Atc for 60 h were used to infect HFF monolayers in 6-well plates. Infected cultures were grown ± Atc for nine days before fixation with cold methanol and staining with crystal violet.

### Conoid extrusion assay

Conoid extrusion ability was assessed in parasites grown ± Atc for 60 h and incubated for 30 sec with 3 μM of calcium ionophore A23187 in DMSO or absolute DMSO as control. Following fixation with 4% paraformaldehyde (PFA), parasites were deployed on glass slides and scored for conoid protrusion by phase contrast microscopy. Percentage of tachyzoites with extruded conoids was determined by counting 200 parasites per condition in three independent experiments conducted in triplicate.

### Replication assay

Freshly egressed parasites grown ± Atc for 60 h were used to infect HFF monolayers for 20 min in chamber slides. After removal of uninvaded tachyzoites, the cultures were incubated ± Atc for 24 h prior to fixation with 4% PFA. Intracellular parasites were stained with anti-GAP45 antibodies and DAPI and the number of parasites per vacuole was determined, counting 100 vacuoles for each condition in 3 independent experiments performed in triplicate.

### Invasion-attachment assay

Freshly egressed parasites grown ± Atc for 60 h were resuspended in DMEM, 2% NuSerum, 10 mM HEPES and seeded on confluent HFF monolayers in 8-well chamber slides (Eppendorf) at a density of 5×10^5^ tachyzoites/well. Following centrifugation at 100 g for 2 min at 4°C, the cultures were incubated at 37°C for 20 min to allow host cell attachment and invasion, washed with DMEM to remove free extracellular parasites and fixed with 4% PFA. Samples were analyzed by differential IFA using a red-green assay (Huynh et al., 2003). Briefly, to label exclusively attached (extracellular) tachyzoites, non permeabilized samples were incubated with an anti-SAG1 mAb followed by Alexa 488-conjugated anti-mouse IgG antibodies (Invitrogen). After permeabilization with 0.5% Triton X-100 (Sigma-Aldrich), all parasites were stained with rabbit anti-GRA1 antibodies followed by 594-conjugated anti-rabbit IgG antibodies. Attached (green+red) and invaded (red) tachyzoites were enumerated by examining 30 microscopic fields/well at 630x magnification. The data shown are representative of four independent experiments performed in triplicate.

### Evacuole assay

Rhoptry proteins discharge was tested using the evacuole detection assay, as previously described (Håkansson et al., 2001). Briefly, parasites grown for 48 h ± Atc were collected by syringe lysis of HFF monolayers, washed in ice-cold DMEM, resuspended in DMEM supplemented with 1μM Cytochalasin D (Sigma-Aldrich) and incubated on ice for 10 min. Pre-chilled HFF monolayers in DMEM/1μM Cytochalasin D were infected with 8×10^6^ parasites per glass coverslips, centrifuged at 1100 g for 1 min and allowed to settle on ice for 20 min. The medium was replaced with pre-warmed growth medium containing 1μM Cytochalasin D and cultures were incubated for 20 min at 37°C in a water bath. Following fixation with 4% PFA, dual IFA was performed using rabbit anti-GAP45 antibodies to visualize the tachyzoites and an anti-ROP1 mAb to detect evacuoles. Results are representative of three independent experiments performed in triplicate.

### Induced egress assay

Freshly harvested parasites grown ± Atc for 60 h were used to infect HFF cultures on 12 mm glass coverslips. After 30 h of growth ± Atc, infected cultures were washed with PBS, incubated for 2 min at 37°C in 2 μM calcium ionophore A23187 diluted in Hank’s Balanced Salt Solution and fixed with 4% PFA. Egressed versus non egressed vacuoles were scored by IFA staining the parasites with an anti-SAG1 mAb and parasitophorous vacuole membranes with rabbit anti-GRA7 antibodies. The percentage of egressed vacuoles was determined in three independent experiments conducted in triplicate by counting 100 vacuoles for each experimental condition.

### Gliding motility assay

Intracellular parasites grown ± Atc for 60 h were mechanically extruded from HFF monolayers, resuspended in Hank’s Balanced Salt Solution and seeded in 8-well poly-Lysine-coated (0.025%; Sigma-Aldrich) chamber slides (Falcon) at a density of 200,000 parasites/well. Following centrifugation at 100 g for 2 min, parasites were incubate for 30 min at 37°C ± Atc, fixed with 4% PFA and surface stained with an anti-SAG1 mAb to reveal fluorescent SAG1 trails. Results are representative of three independent experiments carried out in triplicate.

### Transmission Electron Microscopy

Freshly egressed tachyzoites grown ± Atc for 60 h were fixed with 4% paraformaldehyde/0.1% glutaraldehyde in 0.1 M sodium cacodylate buffer overnight at 4°C. Next, the suspensions were gently washed in buffer, dehydrated in ethanol serial dilutions and embedded in LR White, medium-grade acrylic resin (London Resin Company, UK). The samples were polymerised in a 55°C oven for 48 h and ultrathin sections, obtained by an UC6 Ultramicrotome (Leica), were collected on gold grids, stained by uranyl acetate 2% in H_2_O for 10 min and examined at 100kV by EM 208S Transmission Electron Microscope (FEI - Thermo Fisher) equipped with the acquisition system Megaview SIS camera (Olympus).

### Bionformatic tools

Genome-wide homology searches were performed using the PSI-BLAST and BLASTP algorithms (https://blast.ncbi.nlm.nih.gov), multiple amino acid sequence comparisons with CLUSTAL Omega (https://www.ebi.ac.uk/Tools/msa/clustalo/). Leader peptide and TM regions were predicted with the programs SignalP 5.0 (https://services.healthtech.dtu.dk/service.php?SignalP-5.0) and TMHMM 2.0 (https://services.healthtech.dtu.dk/service.php?TMHMM-2.0), respectively. Protein domain architecture was analyzed with the Simple Modular Architecture Research Tool (SMART; http://smart.embl-heidelberg.de/) and by interrogating the Pfam (https://pfam.xfam.org/) and NCBI CDD (https://www.ncbi.nlm.nih.gov/Structure/cdd/cdd.shtml) databases. Phylogenetic analyses were performed with the MEGA 11 software, using MIC14 and MIC15 amino acid sequences aligned with the MUSCLE software.

### Statistical analysis

Statistical analyses were carried out with Prism 9 (GraphPad Software Inc., La Jolla, CA, USA) using Student’s t tests as indicated in the Figure legends. All assays were performed in triplicate and results are shown as mean values ±SEM of at least three independent biological replicates. Differences were considered significant if P values were <0.05.

## RESULTS

### Definition of the repertoire of thrombospondin-related proteins in *Toxoplasma gondii*

To expand our knowledge of the repertoire of TSP1 domain-containing proteins encoded by the *T. gondii* genome, we searched the ToxoDB database with the PSI-BLAST algorithm using as query the amino acid consensus sequence WXXWXXCXXXC spanning highly conserved amino acid positions within the TSP1 domain. This allowed us to identify nine additional polypeptides containing one or more copies of this amino acid motif. Table 1 summarizes the main characteristics of the predicted proteins, including their major structural features, the expression in different invasive stages and the fitness-conferring score (FCS). This value, derived from a genome-wide loss of function genetic screen (Sidik et al., 2016), ranks individual *T. gondii* proteins based on their indispensability during the tachyzoite lytic cycle in cultured human fibroblasts. Interestingly, seven of the newly identified TSP-related proteins had FCSs ranging from −0.48 to −3.9, with higher negative values indicating a higher degree of protein essentiality. With the exception of TGME49_279420 and TGME49_319992, all the identified proteins possess a single C-terminal transmembrane region (TMR), however, only TGME49_209060 and TGME49_247195 have a putative N-terminal leader peptide for targeting to the secretory pathway. The lack of this amino acid element in the other proteins is possibly due to inaccurate 5’-end predictions. Out of five TSP-related genes lacking proteomic-based expression evidence of the encoded proteins in tachyzoites, bradyzoites or sporozoites, four (*TGME49_223480, TGME49_237585, TGME49_247970* and *TGME49_319992*) show transcriptional profiles compatible with expression in *T. gondii* enteric stages. Instead, *TGME49_218310* exhibits no significant stage-specific upregulation. Mass-spectrometry data support expression of the four remaining proteins in tachyzoites and in three cases (TGME49_209060, TGME49_247195 and TGME49_279420) also in bradyzoites and sporozoites.

**Table 1.** Uncharacterized thrombospondin-related proteins in *Toxoplasma gondii*.

| ToxoDB gene ID | Chr. n. | ToxoDB annotation | FCS<sup>a</sup> | #TSP1 domains | Other domains | LP | TMR | length | Proteomics<sup>b</sup> | Transcriptomics<sup>c</sup> |
| --- | --- | --- | --- | --- | --- | --- | --- | --- | --- | --- |
| TGME49_223480 | X | sushi domain-containing protein | -3.9 | 16 | Sushi, NL | no | yes | 4752 | nd | EES |
| TGME49_247195 | XII | Microneme protein MIC15 | -3.31 | 3 | H-type lectin | yes | yes | 2909 | T,S,B | T,S,B |
| TGME49_277910 | XII | TSP1 domain-containing protein | -2.95 | 1 | -- | no | yes | 1232 | T | all stages |
| TGME49_279420 | IX | Hypothetical protein | -1.96 | 1 | -- | no | no | 1329 | T,S,B | T,S,B |
| TGME49_209060 | Ib | TSP1 domain-containing protein | -1.7 | 7 | -- | yes | yes | 876 | T,S,B | all stages |
| TGME49_247970 | XII | Hypothetical protein | -0.96 | 2 | -- | yes | yes | 1237 | nd | EES |
| TGME49_319992 | IV | Hypothetical protein | -0.48 | 1 | -- | no | no | 223 | nd | EES |
| TGME49_218310 | XII | Microneme protein MIC14 | 0.28 | 3 | -- | no | yes | 968 | nd | all stages |
| TGME49_237585 | X | Hypothetical protein | 1.41 | 2 | -- | no | yes | 1253 | T* | EES |

<sup>a</sup>FCS: fitness conferring score [24] indicating protein essentiality (highly negative values) or dispensability (positive values)
<sup>b</sup>Stages in which the protein was identified by mass-spectrometry according to proteomic studies reported in ToxoDB
<sup>c</sup>Stages with the relative highest abundance of gene transcript as deduced from ToxoDB data
\*Weak single-peptide identification
Abbreviations: B, bradyzoite; EES, enteroepithelial stages; LP, leader peptide; nd, no data; NL, Notch domain; S, sporozoite; T, tachyzoite; TMR, transmembrane region; vWA, von Willebrand factor type A domain

The protein TGME49_247195, annotated in ToxoDB as MIC15, is characterized by a markedly negative FCS (−3.31), indicative of an important functional role. The predicted 2924 amino acid long MIC15 consists of a N-terminal leader peptide, a large ectodomain spanning three TSP1 motifs and a C-terminal TMR followed by a putative cytoplasmic tail. This overall architecture closely resembles that of several type I transmembrane adhesins secreted by the micronemes, and key to *T. gondii* gliding motility (MIC2) (Gras et al., 2017), rhoptry secretion (MIC8) (Kessler et al., 2008) or moving junction formation (AMA1) (Alexander et al., 2005; Besteiro et al., 2009). Furthermore, BLAST analysis showed that the 968 amino acid long TGME49_218310 identified by our search and annotated in ToxoDB as MIC14, shares 27% identity and 41% similarity with the C-terminal third of MIC15 and a comparable domain organization. Based on the hypothesis that MIC15 might play an important role in parasite-host cell interaction and that the leaderless MIC14 might represent an incompletely predicted and closely related homolog, we decided to characterize the two proteins in more detail.

### Structure of the *MIC14* and *MIC15* genomic loci

To verify the ToxoDB gene models of *MIC14* and *MIC15*, which lie about 1.6 Mb apart on chromosome XII, we screened a *T. gondii* (strain RH) cDNA library using a PCR-based limiting dilution method (Israel, 1993) and isolated two polyadenylated cDNAs consisting of 4259 bp for *MIC14* and 5656 bp for *MIC15* (**Figure 1A,B**). The *MIC14* cDNA fully accounted for the 17 exons gene model displayed in ToxoDB, whereas the *MIC15* cDNA spanned only the 23 terminal exons of the predicted genomic locus. Using gene-specific reverse primers designed in the 5’ end of the newly isolated cDNAs, we subjected total tachyzoite RNA to 5’ RACE elongations. By this approach we extended the *MIC15* transcript by approximately 3.8 Kb (**Figure 1A**), obtaining a combined full-length cDNA sequence of 9473 bp that confirmed the 42 exons structure of the genomic locus and the MIC15 amino acid sequence reported in ToxoDB. On the contrary, 5’ RACE experiments yielded no *MIC14*-specific elongation products. However, a BLASTP search of ToxoDB with the MIC15 amino acid sequence as query provided indirect evidence that the *MIC14* transcript might extend considerably upstream of the cDNA 5’ end. In fact, MIC15 showed significant homology (25% identity, 40% similarity) with the hypothetical protein TGME49_218340 encoded by the gene lying immediately upstream of the equally oriented *MIC14* (**Figure 1B**). Interestingly, the 1187 amino acid long TGME49_218340 aligned for its entire length with the two N-terminal thirds of MIC15, strongly suggesting that *TGME49_218340* and *TGME49_218310 (MIC14*) belong to the same transcriptional unit. As represented in Figure 1B, this hypothesis was confirmed by RT-PCR amplification of a 4.5 Kb product that spanned the entire *TGME49_218340* transcript and overlapped with the cloned *MIC14* cDNA. Using poly-A^+^tachyzoite RNA, the resulting transcript was further extended by 5’ RACE, yielding a combined mRNA sequence of 7593 bp (GenBank accession number AY089992) that encoded a putative N-terminal signal-anchor sequence, and brought MIC14 length from 968 to 2344 amino acids. As shown in Figure 1C, the corresponding genomic locus consists of 43 exons, 25 of which are perfectly conserved with respect to *MIC15*.

**FIGURE 1.**
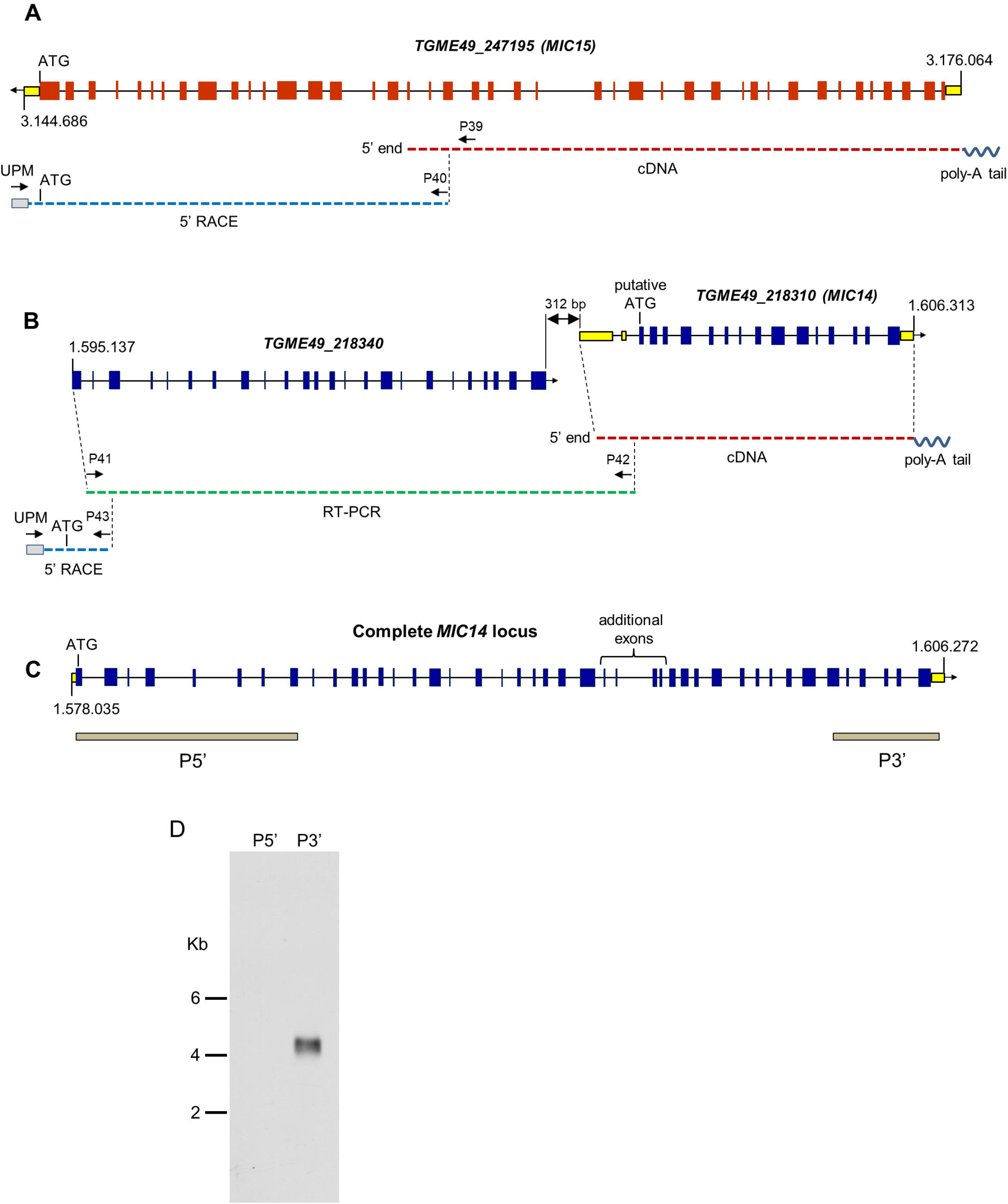
Structure of the *MIC14* and *MIC15* genomic loci and analysis of *MIC14* transcription. Schematic representation of the ToxoDB gene models of *MIC15* **(A)** and *MIC14* with the adjacent *TGME49_218340* **(B)**. Predicted exons are shown as red (*MIC15*) or blue (*MIC14* and *TGME49_218340*) boxes, untranslated regions as yellow boxes, intronic sequences as black lines. The numbers at the extremes of the gene models correspond to the nucleotidic positions on chromosome XII. Cloned cDNAs, RT-PCR amplicons or 5’RACE products are represented as red, green or blue dashed lines, respectively. The position of the primers used for transcript amplification is indicated by black arrows. **(C)** Structure of the complete *MIC14* locus as deduced from the combined sequence resulting from the overlap between a library derived cDNA, RT-PCR and 5’ RACE products. The grey bars indicate the exons spanned by the two probes used for Northern blot hybridizations. **(D)** Northern blot analysis of tachyzoite total RNA with the *MIC14* probes P5’ and P3’. Exposure time, 96 hours.

To further investigate transcription at the *MIC14* locus, we carried out a Northern blot analysis of tachyzoite RNA using the probes P5’ and P3’ (**Figure 1D**), mapping at the two extremes of the *MIC14* locus. While the 3’ end probe recognized a single RNA species of approximately 4.5 Kb compatible with the size of the cloned *MIC14* cDNA, the 5’ end probe showed no hybridization, suggesting that in *T. gondii* tachyzoites the *MIC14* locus is predominantly transcribed as a 5’-truncated mRNA form of approximately 4.5 Kb while the full-length transcript is scarcely represented.

### Phylogenetic relationship and structural organization of MIC14 and MIC15

BLASTP searches for MIC14 and MIC15 homologs in vEuPathDB (**https://veupathdb.org/**) and in the NCBI non-redundant proteins database recovered highly homologous sequences exclusively from the phylum Apicomplexa and in particular from the Coccidia class, that includes parasitic genera of high veterinary relevance such as *Eimeria, Neospora* and *Sarcocystis*. As illustrated in Figure 2A, the comparison between the amino acid sequences of MIC14 and MIC15 homologs retrieved from eight genera of Coccidia yielded a robust phylogenetic tree demonstrating that the two proteins have a paralogous relationship.

**FIGURE 2.**
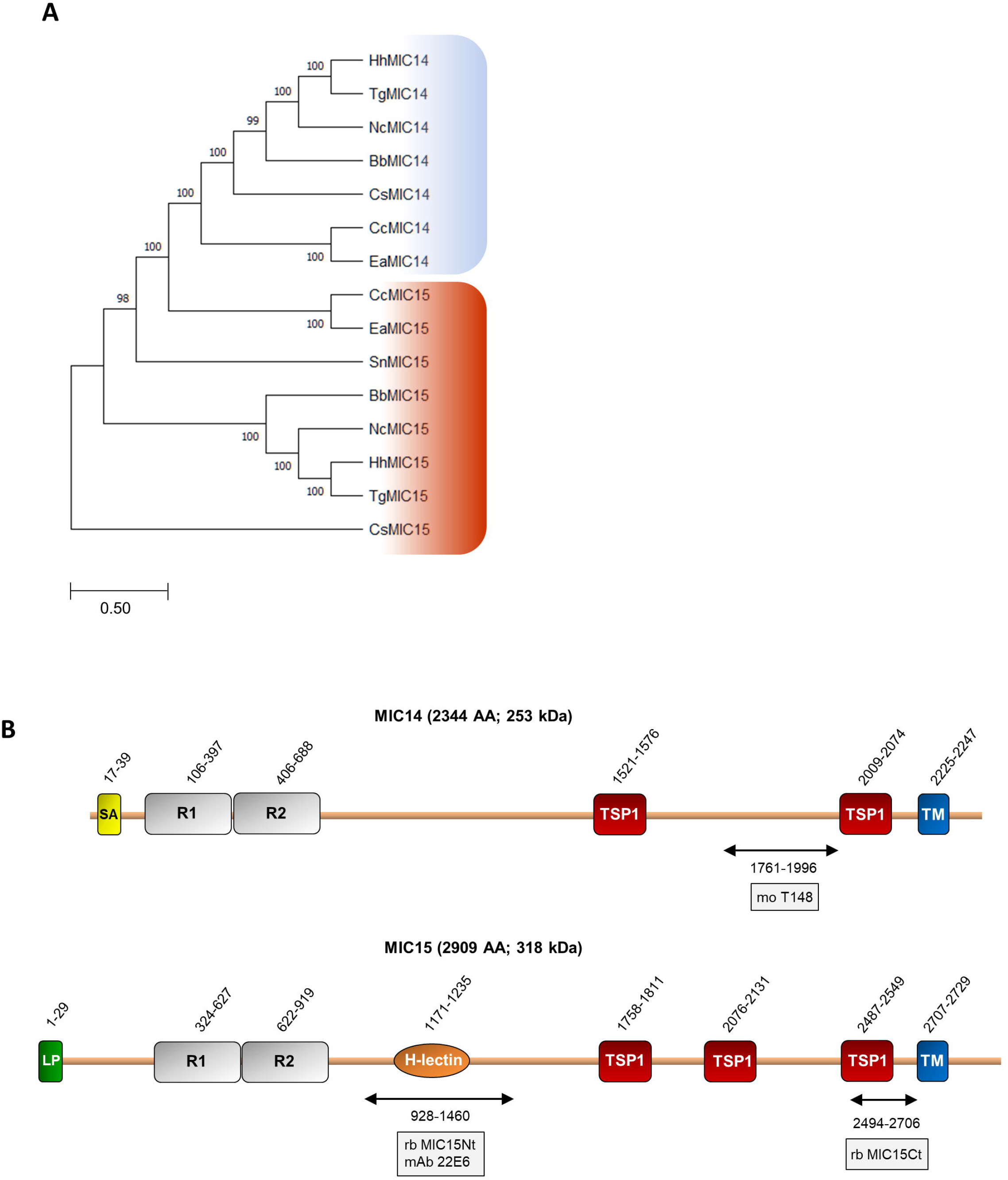
Phylogenetic analysis and domain organization of MIC14 and MIC15. **(A)** Unrooted maximum likelihood phylogenetic tree of MIC14 and MIC15 proteins in Apicomplexa. Bootstrap values for 1000 trials are shown. The amino acid alignment used to generate the tree is shown in Supplementary Figure 8. Amino acid sequences used to generate the phylogenetic tree are identified by the following accession numbers: *Toxoplasma gondii*, MIC14 (AY089992), MIC15 (DQ459408); *Besnoita besnoiti*, MIC14 (XP_029221473), MIC15 (XP_029222264); *Cyclospora cayetanensis*, MIC14 (XP_026191791), MIC15 (XP_026191793); *Cystoisospora suis*, MIC14 (PHJ15367), MIC15, (PHJ22147); *Eimeria acervulina*, MIC14 (XP_013251268), MIC15 (XP_013251270); *Hammondia hammondi*, MIC14, (HHA_218310), MIC15, (HHA_247195); *Neospora caninum*, MIC14 (CEL70510), MIC15 (CEL70693); *Sarcocystis neurona*, MIC15 (SN3_00200915). **(B)** Schematic representation of *T. gondii* MIC14 and MIC15 full-length proteins. The numbers identify the amino acid positions of major domains or elements or the extremes of the protein regions (double arrows) expressed in bacteria for antibody production. The names of the antibodies are boxed. The numbers of amino acids and the predicted molecular mass of the full-length proteins are shown in brackets. H-lectin, H-lectin domain; TSP1, thrombospondin type 1 domain; LP, leader peptide; R1 and R2, N-terminal repeats; SA, signal-anchor sequence; TM, transmembrane region.

Pairwise alignment between the full-length amino acid sequences of MIC14 and MIC15 showed an identity of 19% (44% similarity) and a comparable domain organization (**Figure 2B**). The C-terminal halves of MIC14 and MIC15 are characterized by two and three TSP1 domains, respectively, followed by a C-terminally located TMR and cytoplasmic tails of 97 and 180 amino acids, respectively. Moreover, the N-terminal region of both proteins contains two tandemly arrayed cysteine-rich repeats of approximately 300 amino acids. Nonetheless, the two paralogs differ for the presence in MIC15 of a H-type lectin carbohydrate-binding domain conserved in the majority of the Coccidia genera analyzed. In addition, while MIC15 is predicted to have a cleavable N-terminal leader peptide, in MIC14 this element is replaced by a putative N-terminal signal-anchor sequence. However, these structural features were not strictly conserved throughout evolution, as both MIC14 and MIC15 orthologs exhibit either of the two types of N-terminal signals. Noteworthy, about 82% (108) of the cysteine residues distributed throughout *T. gondii* MIC14 and MIC15 are conserved (**Supplementary Figure 1**), strongly suggesting a common three-dimensional folding.

### Expression and immunolocalization of MIC15

The expression of MIC15 in *T. gondii* tachyzoites was investigated by Western blot using two distinct rabbit polyclonal antibodies denominated MIC15Nt, recognizing a protein region spanning the H-type lectin domain, and MIC15Ct, directed against the C-terminal portion of the ectodomain (**Figure 2B**). In total tachyzoite lysates, both antisera specifically recognized a single protein band >260 kDa, compatible with the MIC15 predicted molecular mass of approximately 300 kDa (**Figure 3A, left panel**). The same antibodies showed no reactivity with the excreted-secreted antigen (ESA) fraction from ethanol-treated parasites (**Figure 3A, right panel**), questioning the possibility that MIC15 is a new member of the group of microneme transmembrane adhesins which are proteolytically shed from the parasite surface upon secretion (Sheiner et al., 2010). In the majority of cases, this process is driven by a parasite rhomboid protease recognizing an intramembrane amino acid sequence which is absent from the TMR of both MIC15 and MIC14 (**Supplementary Figure 2**).

**FIGURE 3.**
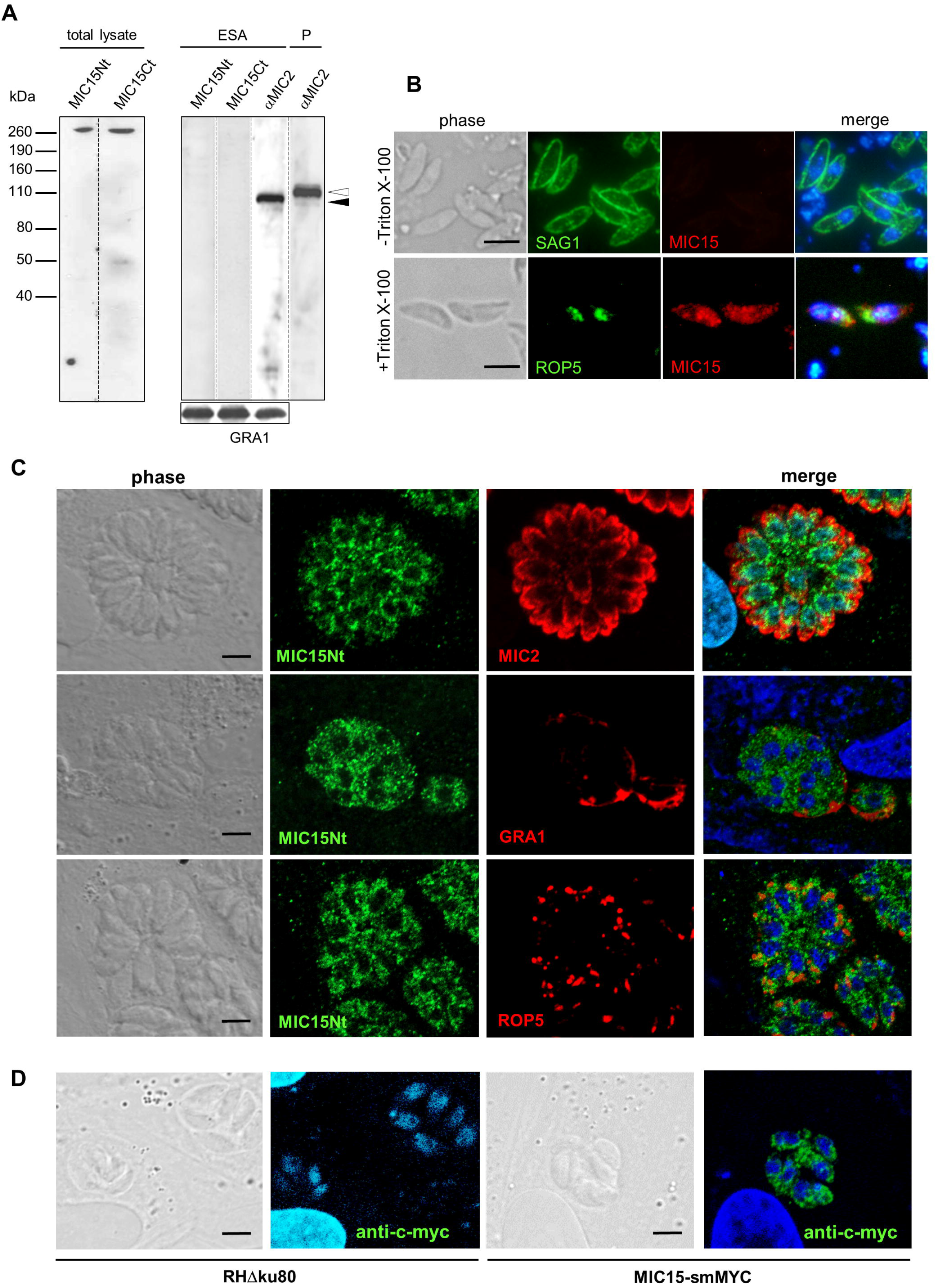
MIC15 is expressed in *T. gondii* tachyzoites and shows a dispersed intracellular localization. **(A)** Western blot analysis of proteins contained in the total tachyzoite lysate (left panel) or in the excreted-secreted antigens (ESA) fraction and in the corresponding parasite pellet (P) (right panel). MIC15 was detected with the rabbit antibodies MIC15Nt or MIC15Ct (Figure 2B). The anti-MIC2 mAb T34A11 was used as positive control of the ESA fraction and its associated pellet. The intracellular and the cleaved extracellular forms of MIC2 are indicated by white and black arrowheads, respectively. *T. gondii* GRA1 was used as loading control for the ESA fraction. **(B)** Immunolocalization of MIC15 in extracellular tachyzoites under non permeabilizing (-Triton X-100) or permeabilizing (+Triton X-100) conditions. Intact or membrane permeabilized RH parasites were stained with anti-SAG1 or anti-ROP5 mAbs, respectively, in conjunction with the rabbit serum MIC15Nt. Scale bar, 5 μm. **(C)** Immunolocalization of MIC15 in intracellular *T. gondii* tachyzoites with the rabbit serum MIC15Nt. Parasites were double stained with antibodies to protein markers specific for the micronemes (MIC2), the dense granules (GRA1) or the rhoptries (ROP5). **(D)** Immunolocalization of MIC15 tagged with the smMYC protein, that consists of 10 copies of the c-myc epitope inserted into a darkened version of the green fluorescence protein. The punctate cytoplasmic distribution of tagged MIC15 disclosed by the anti-c-myc mAb is comparable to that of the native protein shown in (C). Nuclei are stained with DAPI. Scale bar, 5 μm.

The possible exposure of MIC15 on the parasite PM was investigated by indirect immunofluorescence analysis (IFA) in non permeabilized tachyzoites of strain RH dual stained with the MIC15Nt rabbit serum and an anti-SAG1 mAb as positive control. As shown in Figure 3B, the anti-MIC15 antibodies showed no surface reactivity, whereas in detergent treated parasites they produced a punctate diffuse cytoplasmic signal. This unexpected localization was then studied in intravacuolar tachyzoites using the rabbit serum MIC15Nt in conjunction with antibodies to a series of reference intracellular structures (**Figure 3C**). The staining pattern confirmed a diffuse dotted distribution of the protein throughout the parasite cytoplasm and showed no MIC15 colocalization with apical organelles, micronemes and rhoptries, or with dense granule proteins. This unexpected result was independently supported by IFA of three distinct parasite lines in which MIC15 was endogenously tagged by adding either the spaghetti monster-c-myc (smMYC) protein at the N-terminus, a Ty epitope upstream of the TMR or two HA tags at the C-terminus. All endogenous taggings were confirmed by Western blot analysis, showing that tagged strains expressed similar MIC15 protein levels (**Supplementary Figure 3A-C**). All three mutants exhibited the same punctate staining pattern irrespective of the tagging strategy adopted (**Figure 3D and Supplementary Figure 3D**). Taken together, the immunolocalization studies showed that MIC15 has a diffuse cytoplasmic distribution reminiscent of that recently reported for a series of proteins essential for *T. gondii* rhoptry secretion, including the calcium sensor TgFER2 (Coleman et al., 2018) and members of the so-called “non-discharge” (Nd) protein family conserved in the Alveolata superphylum (Aquilini et al., 2021).

### Generation of a *MIC15* conditional knockout mutant

To gain functional insights into MIC15 function, we tried to disrupt the encoding gene by double homologous recombination in the RHΔ*ku80 T. gondii* strain. Following repeated failures of this approach supporting the predicted essentiality of MIC15, we generated an inducible knockdown parasite line, named MIC15-iKD, in the TATiΔ*ku80* strain [28,29]. To this aim, we replaced the cell cycle regulated promoter of *MIC15* with a constitutive, tetracycline-repressible regulatory element (TRE), consisting of seven tandemly arrayed TetO sequences placed immediately upstream of the SAG4 minimal promoter S4 (**Figure 4A**). Correct construct integration and endogenous promoter replacement was confirmed by PCR (**Figure 4B**). Western blot analysis showed a marked increase of MIC15 expression in strain MIC15-iKD compared to the TATi parental line (**Figure 4C**). As determined by quantitative real-time PCR, this upregulation was paralleled by 2.8-fold higher levels of MIC15 mRNA (**Figure 4D**). In MIC15-iKD parasites grown in presence of 0.5 μg/ml Atc for various time periods, Western blot analysis revealed a marked reduction of MIC15 expression after 24 h, with the protein dropping to undetectable levels from the 29 h time point onwards (**Figure 4C**). Accordingly, IFA demonstrated that Atc treatment of the knockdown strain caused the loss of MIC15 expression (**Figure 4E**). Of notice, untreated MIC15-iKD parasites exhibited an evident perinuclear staining compared to the parental strain, indicative of protein accumulation in the endoplasmic reticulum (**Figure 4E and Supplementary Figure 4**). This is likely due to increased *MIC15* expression under the control of the constitutive TRE/S4 promoter. To evaluate the overall impact of MIC15 depletion on the tachyzoite lytic cycle we carried out plaque assays in which HFF monolayers were infected with 250 tachyzoites of either the TATi parental strain or the MIC15-iKD mutant. Unlike the parental strain, MIC15-iKD tachyzoites displayed a normal plaque forming ability only in absence of Atc treatment but were unable to form plaques in presence of the drug, even using a 10-fold higher parasite inoculum (**Figure 4F**). This marked defect was fully restored by complementation of the mutant with an extra-copy of the wild type *MIC15* gene (**Figure 4G,H; Supplemetary Figure 5**).

**FIGURE 4.**
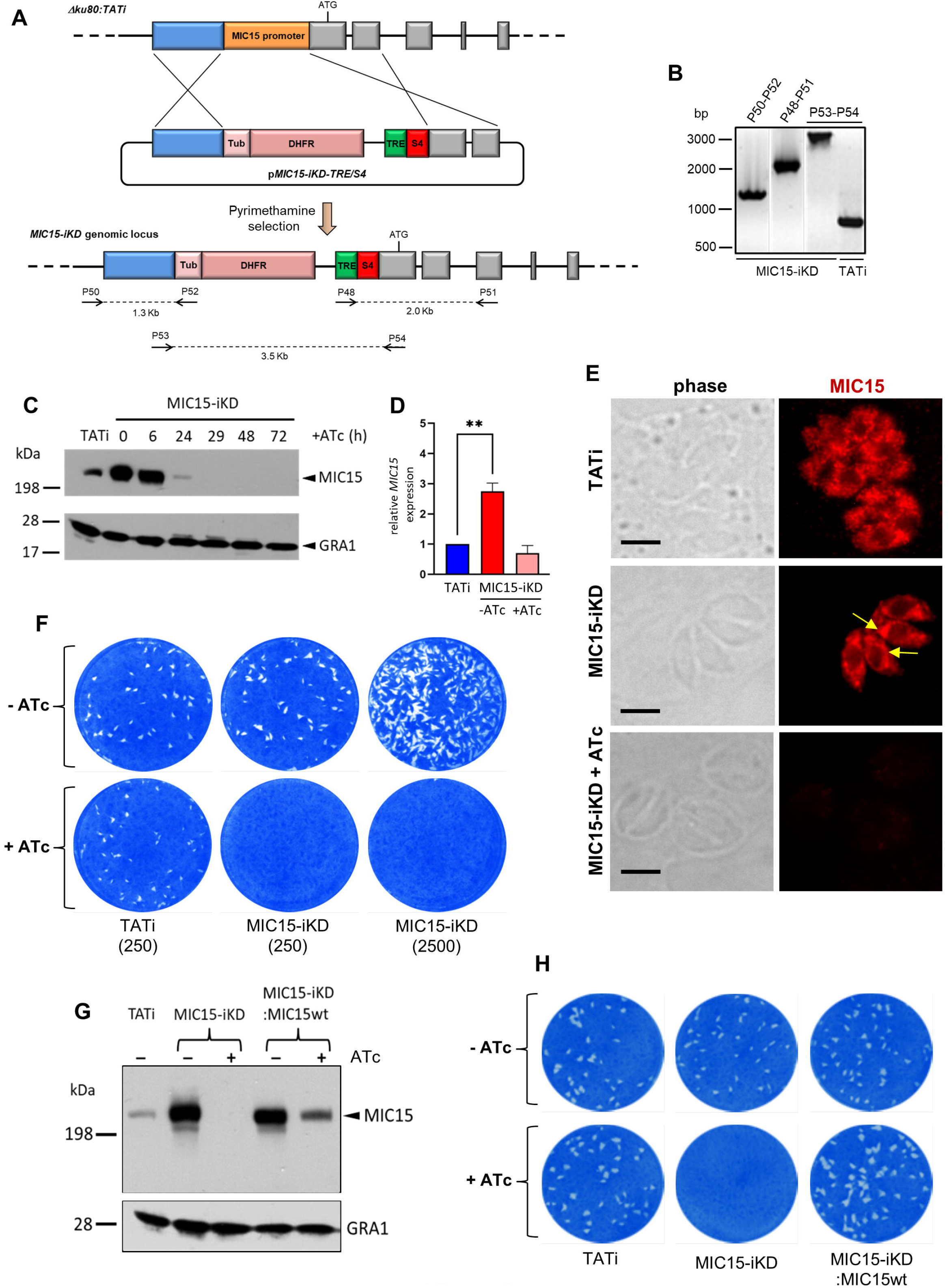
Establishment and validation of a MIC15-iKD conditional-knockdown strain. **(A)** Double homologous recombination strategy used to replace the *MIC15* edogenous promoter with the Tetracycline Responsive Element (TRE) and the SAG4 minimal promoter. The 5’-terminal exons of *MIC15* are represented as grey boxes. The arrows represent the primers used for diagnostic genomic PCR amplifications. **(B)** PCR analysis of the *MIC15-iKD* strain confirming the correct genomic integration of the DHFR/TRE-SAG4 cassette. **(C)** Time course Western blot analysis of MIC15-iKD tachyzoites treated with 0.5 μg/ml Atc for 6 to 72 h showing downregulation of MIC15 below detectable level at 29 h. GRA1 was used as loading control. **(D)** Quantitative real-time RT-PCR analysis of *MIC15* transcription normalized to mRNA levels in strain TATi. Values represent the means ± SEM of three independent experiments. Statistical significance was determined by the Student’s two tailed t-test. **p<0.01. **(E)** Immunofluorescence analysis of intracellular parasites demonstrating MIC15 depletion in Atc-treated MIC15-iKD tachyzoites. The arrows indicate the perinuclear accumulation of MIC15 due to protein overexpression in MIC15-iKD strain. Nuclei are stained with DAPI. Scale bars, 5 μm. **(F)** Plaque assay of the TATi parental strain and MIC15-iKD mutant grown for 9 days in presence or absence of Atc. The numbers in brackets refer to the initial inoculum of tachyzoites. **(G)** Western blot analysis showing the rescue of MIC15 expression in MIC15-iKD parasites complemented with a second copy of the wt gene. **(H)** Plaque assay showing full restore of the lytic phenotype in complemented MIC15-iKD:wtMIC15 tachyzoites. In all experiments MIC15 was detected with the rabbit serum MIC15Nt.

### Secretory organelle localization and microneme discharge are not affected by MIC15 deficiency

The strong phenotype displayed by the knockdown strain upon Atc treatment suggested that MIC15 is involved in one or more key steps underlying the tachyzoite lytic cycle. This prompted us to dissect this complex process by a series of observational and functional assays. The replication rate of MIC15-iKD parasites was compared to that of the parental strain by counting the number of tachyzoites per vacuole at 24 h after pulse infection of HFF monolayers (**Supplementary Figure 6A**). This assay showed similar distribution of vacuole size irrespective of parasite pretreatment with Atc for 60 h, indicating that MIC15 loss has no influence on tachyzoite division. Also the efficiency of conoid extrusion, a calcium-dependent process which occurs during gliding, invasion and egress, was evaluated with a specific test. Extracellular parasites were incubated in a medium containing or not the calcium ionophore A23187, fixed and microscopically scored for the presence or absence of conoid protrusion. No significant difference was observed between MIC15-depleted and control tachyzoites of the TATi and MIC15-iKD strains (**Supplementary Figure 6B**).

Using confocal IFA we explored the possibility that MIC15 downregulation might grossly alter the biogenesis or positioning of the tachyzoite secretory organelles, which orchestrate the interaction with the host cell at multiple levels. To this end, MIC15-iKD tachyzoites treated or not with Atc for 60 h were dual stained with serum MIC15Nt together with mAbs recognizing either MIC2 or MIC8, known to be targeted to distinct subpopulations of micronemes (Kremer et al., 2013), the rhoptry marker ROP5 or the dense granule protein GRA1. As shown in Figure 5A-D, all selected proteins appeared comparably abundant and correctly localized in both strains and regardless of Atc treatment. Furthermore, the efficiency of microneme discharge was investigated by Western blot by detecting MIC2, MIC4 and MIC8 in the ESA fraction of parasites unstimulated or treated with 1% ethanol to induce microneme discharge. As shown in Figure 5E, this non quantitative assay showed that the supernatant of TATi and MIC15-iKD strains contained comparable levels of the three microneme markers irrespective of MIC15 expression. A normal microneme functionality was indirectly suggested also by two distinct in vitro assays that measured the ability of tachyzoites to glide on a solid substrate or egress from the host cell upon stimulation with 2 μM of the calcium ionophore A23187. Both processes, which are strictly dependent on the secretion of specific micronemal effectors such as MIC2 (Huynh and Carruthers, 2006) and PLP1 (Kafsak et al., 2009), respectively, were shown to occur with equal efficiency in the parental strain and in MIC15-iKD parasites regardless of Atc treatment (**Figure 5F,G**). Collectively, these results demonstrate that MIC15 downregulation does not affect microneme secretion.

**FIGURE 5.**
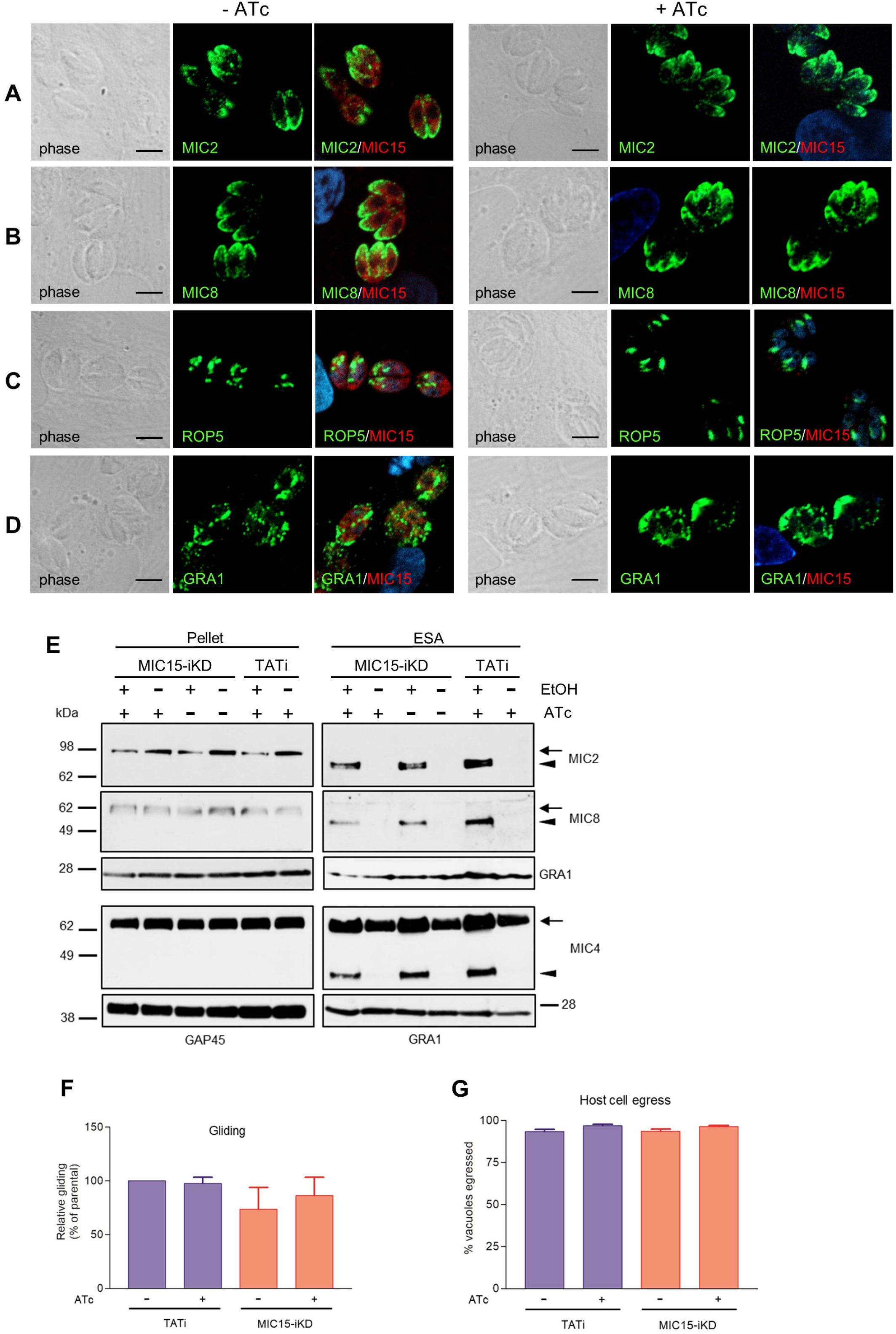
Impact of MIC15 depletion on organelle localization and microneme-dependent functions. **(A-D)** Immunolocalization of secretory proteins in MIC15-iKD tachyzoites grown in presence or absence of Atc for 60 h. The parasites were stained with rabbit serum MIC15Nt plus a mAb specific for either MIC2 (A), MIC8 (B), ROP5 (C) or GRA1 (D). Nuclei are stained with DAPI. Space bar, 5 □M. **(E)** Western blot analysis of constitutive and ethanol-induced microneme secretion in MIC15-iKD parasites. The presence of MIC2, MIC4 and MIC8 was revealed in the cytosolic (pellet) and excreted/secreted antigen (ESA) fractions of TATi and MIC15-iKD tachyzoites grown ±Atc for 60 h. The uncleaved (intracellular) and processed (extracellular) forms of the microneme proteins are indicated by arrows or arrowheads, respectively. GRA1 was used as loading control for the ESA fractions, while cytosolic proteins were normalized with either GRA1 or GAP45. **(F)** Gliding assay comparing the fraction of motile TATi and MIC15-iKD tachyzoites as revealed by the presence of SAG1^+^ posterior trails deposited on poly-Lysine-coated glass chamber slides. Prior to incubation on the substrate for 30 min, the two strains were grown for 60 h in presence or absence of Atc. **(G)** Egress assay estimating the capability of untreated or Atc-treated tachyzoites of strains TATi and MIC15-iKD to exit the parasitophorous vacuole upon stimulation with the calcium ionophore A23187. The data in panels F and G are expressed as mean values ± SEM from three independent experiments conducted in triplicate. In both assays, the Student’s two tailed *t*-test did not reveal statistically significant differences between strains irrespective of MIC15 depletion.

### MIC15 depletion dramatically inhibits host cell invasion

The inability of MIC15-depleted tachyzoites to produce plaques in human fibroblasts was further investigated in vitro by measuring the efficiency of active host cell penetration by TATi and MIC15-iKD parasites. To this end, the two strains were grown for 60 h in presence or absence of Atc and used to infect HFF monolayers for 20 min at 37°C. After removal of free exracellular tachyzoites by extensive washing, the cultures were fixed with 4% paraformaldehyde and invaded (intracellular) versus attached (extracellular) parasites were enumerated by differential IFA using a red-green assay (Huynh et al., 2003). As shown in Figure 6A, Atc treatment of MIC15-iKD tachyzoites significantly reduced invasion by approximately 80% compared to the untreated control and the parental strain. Noteworthy, this marked inhibition was not accompanied by a reduction in the number of attached MIC15-iKD parasites, which rather showed a significant increase upon MIC15 depletion. This result excluded that the strong MIC15-iKD phenotype was due to the impairment of host cell attachment, strongly indicating a relation to molecular events occurring downstream along the invasion process.

**FIGURE 6.**
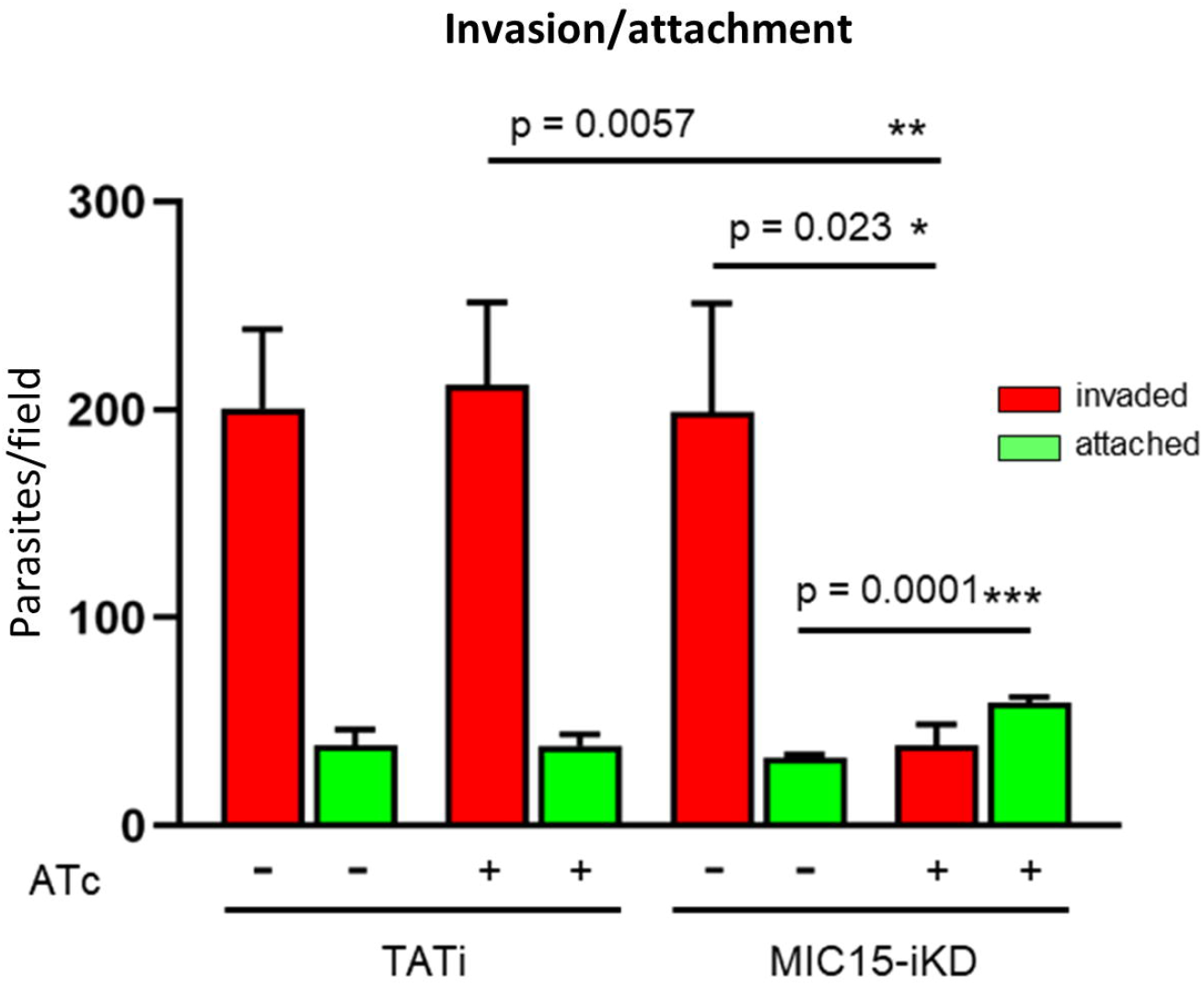
MIC15 depletion impacts on host cell invasion. **(A)** Invasion assay showing a marked reduction of the number of invaded tachyzoites (red bars) following Atc-induced depletion of MIC15. The green bars represent the number of attached parasites enumerated for each strain and condition. Values represent means ± SEM from 4 independent experiments performed in triplicate. Statistical significance was determined by the Student’s two tailed *t*-test. *p<0.05, **p<0.01, ***p<0.001.

### MIC14 expression restores the invasion defect in MIC15-depleted tachyzoites

In order to explore the existence of possible compensatory mechanisms capable to counterbalance the loss of fitness due to MIC15 downregulation, the evolution of the non invasive phenotype was studied over time in MIC15-iKD tachyzoites grown in presence of Atc for up to 120 days. Surprisingly, we found that as soon as 36 days after the start of the Atc treatment, parasites depleted in MIC15 were able to produce sporadic and very small areas of lysis (**Figure 7A**), compared to the null plaque count of MIC15-iKD tachyzoites treated with Atc for the standard 2.5 days before the plaque assay. Restoration of the lytic phenotype became more evident after 120 days of continuous growth with Atc, as shown by the increased number of lytic events and the formation of distinctly larger plaques, though reduced in size compared to parasites grown in absence of Atc during the assay (**Figure 7A**). To investigate the possible involvement of the MIC15 paralog, MIC14, in the restored lytic ability, tachyzoites treated with Atc for 120 days were subjected to limiting dilution cloning in presence of the drug. Seven independent clones were isolated and screened by Western blot with the mouse serum T148 raised against a recombinant fragment of the MIC14 ectodomain (**Figure 2A**). As shown in Figure 7B for a representative clone denominated MIC15-iKD/MIC14^R^, the polyclonal antibody specifically detected a high molecular weight product which was absent in the parental strains TATi and MIC15-iKD and exhibited a molecular mass compatible with the 253 kDa predicted for MIC14. To validate the hypothesis that the newly identified protein corresponded to MIC14, we successfully knocked out the corresponding gene in MIC15-iKD/MIC14^R^ tachyzoites by replacing the first two exons and upstream region of *MIC14* with a TUB/CAT resistance cassette (**Figure 7C**). Western blot analysis with serum T148 showed that MIC15-iKD/MIC14^R^/MIC14KO parasites lacked the high MW band characteristic of the parental strain, which was unequivocally identified as MIC14. This result was corroborated by the return of MIC15-iKD/MIC14^R^/MIC14KO parasites to the null plaque phenotype of the parental line (**Figure 7E**). The analysis of MIC15-iKD/MIC14^R^ strain by RT-qPCR using a primer pair mapping to exon 16 of the complete gene showed that the upregulation of MIC14 and the consequent rescue of the lytic phenotype was associated with a 12.7-fold increase of the full-length *MIC14* mRNA (**Figure 7F**). The sequencing of 2.3 kb upstream of the MIC14 translation start codon of both MIC15-iKD and MIC15-iKD/MIC14^R^ strains excluded the presence of mutations that could be linked to the observed transcriptional upregulation.

**FIGURE 7.**
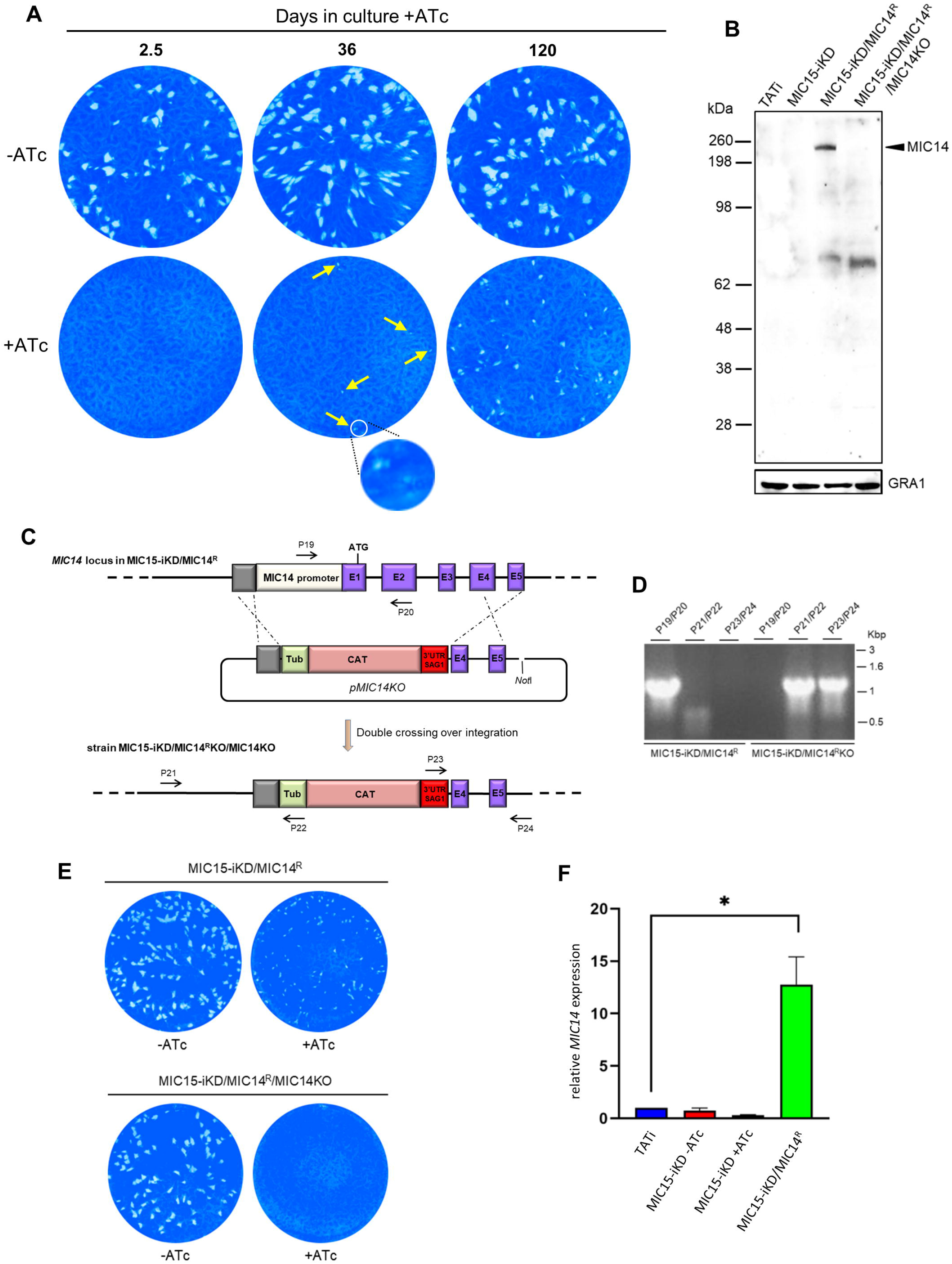
Selection and characterization of a MIC15-iKD variant strain expressing MIC14. **(A)** MIC15-iKD tachyzoites cultured in presence of Atc for extended periods of time show a partial rescue of the lytic phenotype. Minuscule plaques detected after 36 days of Atc treatment are indicated by arrows and a group of them, encircled in white, is shown at higher magnification. Parasites grown in presence of Atc for 120 days formed larger plaques, still significantly smaller than those produced omitting Atc from the assay. **(B)** Western blot analysis of strain MIC15-iKD/MIC14^R^, which was cloned from MIC15-iKD parasites propagated in presence of Atc for 120 days. The anti-MIC14 mouse serum T148 recognized a protein band >198 kDa (arrowhead) in MIC15-iKD/MIC14^R^ tachyzoites which was absent in the TATi and MIC15-iKD parental strains. The reactivity of serum T148 with the >198 kDa product was abolished by the knockout of the *MIC14* gene. **(C)** Schematic diagram of the strategy used to knockout the *MIC14* gene in the MIC15-iKD/MIC14^R^ strain. The promoter region and the first two exons of *MIC14* were replaced with the CAT selectable marker by double homologous recombination at the sites indicated with dashed lines. Exons in the *MIC14* gene are shown as purple rectangles with numbers and introns as continuous lines. Primers used to assess loss of the *MIC14* promoter and 5’ terminal exons and integration of the knockout construct are shown as numbered arrows. **(D)** Analysis of PCR products amplified from the indicated strains with primers shown in panel C. Migration of size standards in kilobase pairs (Kbp) are shown to the right. **(E)** Plaque assay demonstrating restoration of the non lytic phenotype following *MIC14* disruption in MIC15-iKD/MIC14^R^ tachyzoites. **(F)** Quantitative real-time RT-PCR analysis of *MIC14* transcription normalized to mRNA levels in the TATi parental strain. Values represent the means ± SEM of three independent experiments. Statistical analysis was performed by the Student’s two tailed *t*-test.

Attempts to immunolocalize MIC14 in MIC15-iKD/MIC14^R^ tachyzoites using serum T148 were unsuccessful. This prompted us to engineer this strain by endogenously tagging the *MIC14* gene by insertion of three C-terminal Ty epitopes, obtaining the MIC14^R^-3xTy line (**Figure 8A**). To enhance the expression level of the tagged protein, we also generated the strain TUB-MIC14^R^-3xTy by replacing the *MIC14* endogenous promoter with the constitutive promoter of *T. gondii* β-tubulin (**Figure 8A).** Western blot analysis by anti-Ty antibodies showed expression in both mutant strains of the □250 kDa full-length MIC14 and confirmed that TUB-MIC14^R^-3xTy parasites produced higher amounts of the tagged protein (**Figure 8B**). Moreover, anti-Ty antibodies revealed the presence in both mutants of two additional bands of approximately 180 and 100 kDa, suggesting that MIC14 undergoes proteolytic processing at two distinct sites in the extracellular domain of the molecule. Consistent with the Western blot result, MIC14 was localized much more efficiently by the anti-Ty mAb in TUB-MIC14^R^-3xTy parasites, in which the tagged protein displayed the same diffuse cytoplasmic distribution of MIC15 (**Figure 8C**). Taken together, these data demonstrate that MIC15 depletion can favour the emergence of parasite variants overexpressing MIC14 and that this protein, undetectable in wild type tachyzoites, has a MIC15-like punctate localization and can partially compensate the pronounced invasion defect due to MIC15 loss.

**FIGURE 8.**
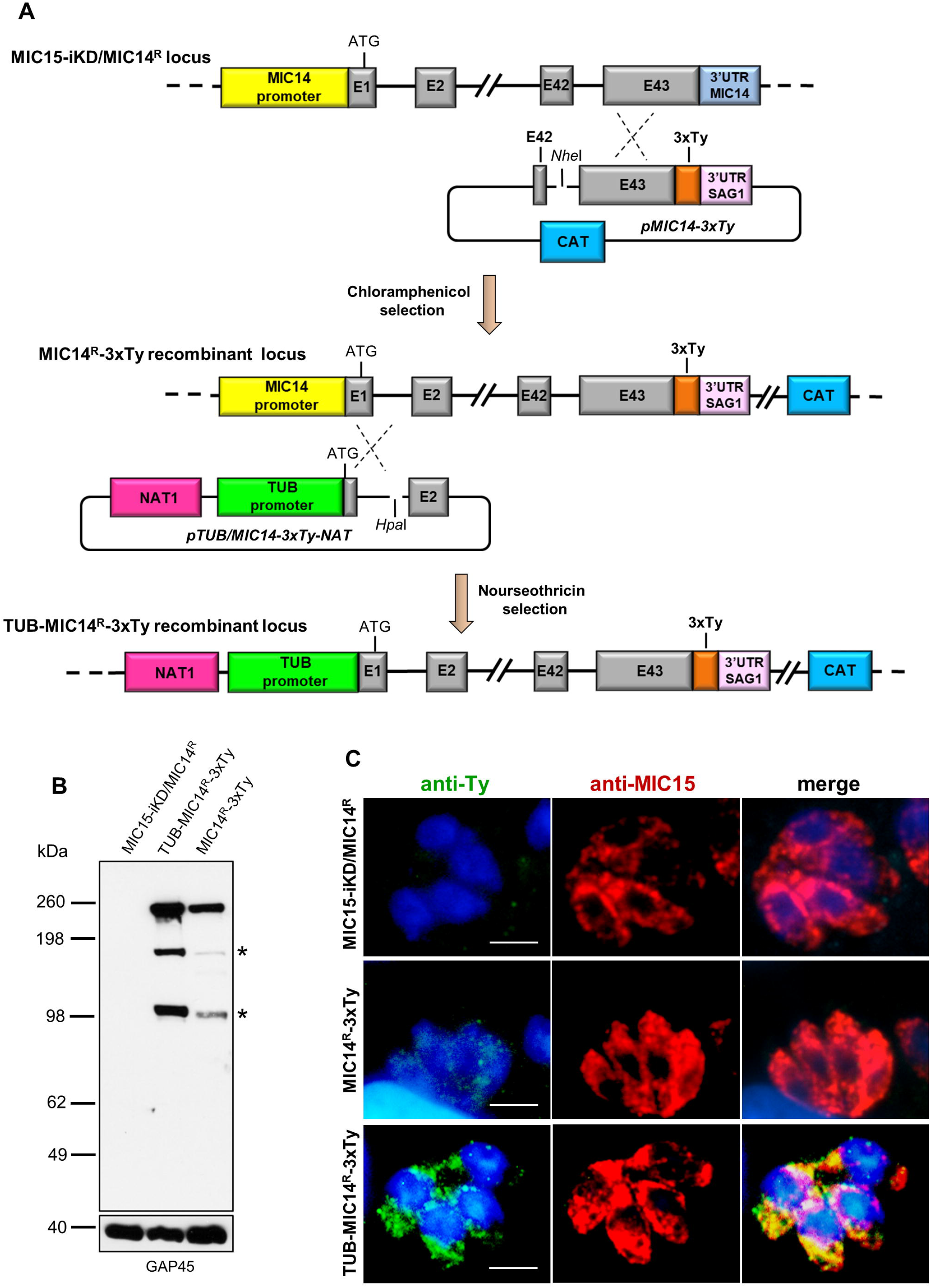
Generation and characterization of parasite lines expressing C-terminally tagged MIC14. **(A)** The upper part of the diagram shows the single homologous recombination strategy for integration of the 3x-Ty epitope sequence (in orange) at the 3’end of the *MIC14* coding sequence in MIC15-iKD/MIC14^R^ parasites. In the obtained chloramphenicol-resistant strain MIC15-iKD/MIC14^R^-3xTy, the *MIC14* locus was further modified by single homologous recombination (lower part of the diagram) by replacing the endogenous promoter of *MIC14* with the *T. gondii* β-tubulin constitutive promoter, obtaining the nourseothricin-resistant strain TUB-MIC14^R^-3xTy. **(B)** Western blot analysis of total protein extracts from MIC15-iKD/MIC14^R^ parasites, expressing the native form of MIC14, and strains MIC15-iKD/MIC14^R^-3xTy and TUB-MIC14^*R*^-3xTy encoding the Ty-tagged version of the protein. In both tagged strains, the anti-Ty mAb recognized the □250 kDa full-length MIC14 and two additional bands of approximately 150 and 100 kDa (indicated by asterisks), likely resulting from specific proteolytic events. As a consequence of promoter swapping, TUB-MIC14^*R*^-3xTy parasites show higher levels of the three MIC14 products. The inner membrane complex protein GAP45 was used as loading control. **(C)** Immunofluorescence detection of MIC14-3xTy in strains MIC15-iKD/MIC14^R^-3xTy and TUB-MIC14^R^-3xTy. The two tagged strains and the parental line MIC15-iKD/MIC14^R^ were dual stained with an anti-Ty mAb and the anti-MIC15 serum MIC15Nt. Nuclei are stained with DAPI. Scale bars, 5 μm.

### MIC15 and MIC14 are involved in rhoptry secretion

To gain insight in the mechanism involved in the impairment of invasion consequent to MIC15 depletion, we explored the possible impact of this condition on rhoptry discharge, which is crucial to moving junction formation and host cell penetration. Rhoptry secretion was investigated by an evacuole assay, consisting in an invasion assay performed in presence of Cytochalasin D (CytD). Under this condition, actin polymerization and therefore parasite invasion are inhibited, without interfering with the ability of extracellular tachyzoites to inject in the host cell cytoplasm small lipid vesicles enriched in rhoptry bulb proteins (evacuoles). The detection of evacuoles by IFA using an anti-ROP1 antibody was used as an indicator of correct rhoptry secretion by parasites attached to the host cell surface. Notably, depletion in MIC15 dramatically reduced evacuole formation by Atc treated MIC15-iKD parasites if compared to the untreated control and the TATi parental strain (**Figure 9**), and to levels comparable to tachyzoites depleted in RON5, key protein in the moving junction complex formation (Beck et al., 2014). On the contrary, Atc treated strain MIC15-iKD/MIC14^R^ did not present the evacuole defect, further supporting the compensatory effect of MIC14 upregulation. Importantly, transmission electron microscopy observations of MIC15-iKD tachyzoites showed that MIC15 downregulation did not cause ultrastructural anomalies in the apical positioning of the rhoptries (**Supplementary Figure 7**). Collectively, our results demonstrate that the paralogous proteins MIC15 and MIC14 are novel components of a *T. gondii* molecular pathway controlling rhoptry exocytosis. Consistent with their newly revealed function, we propose to adopt a more appropriate nomenclature and to rename MIC15 as rhoptry discharge factor 1 (RDF1) and MIC14 as rhoptry discharge factor 2 (RDF2).

**FIGURE 9.**
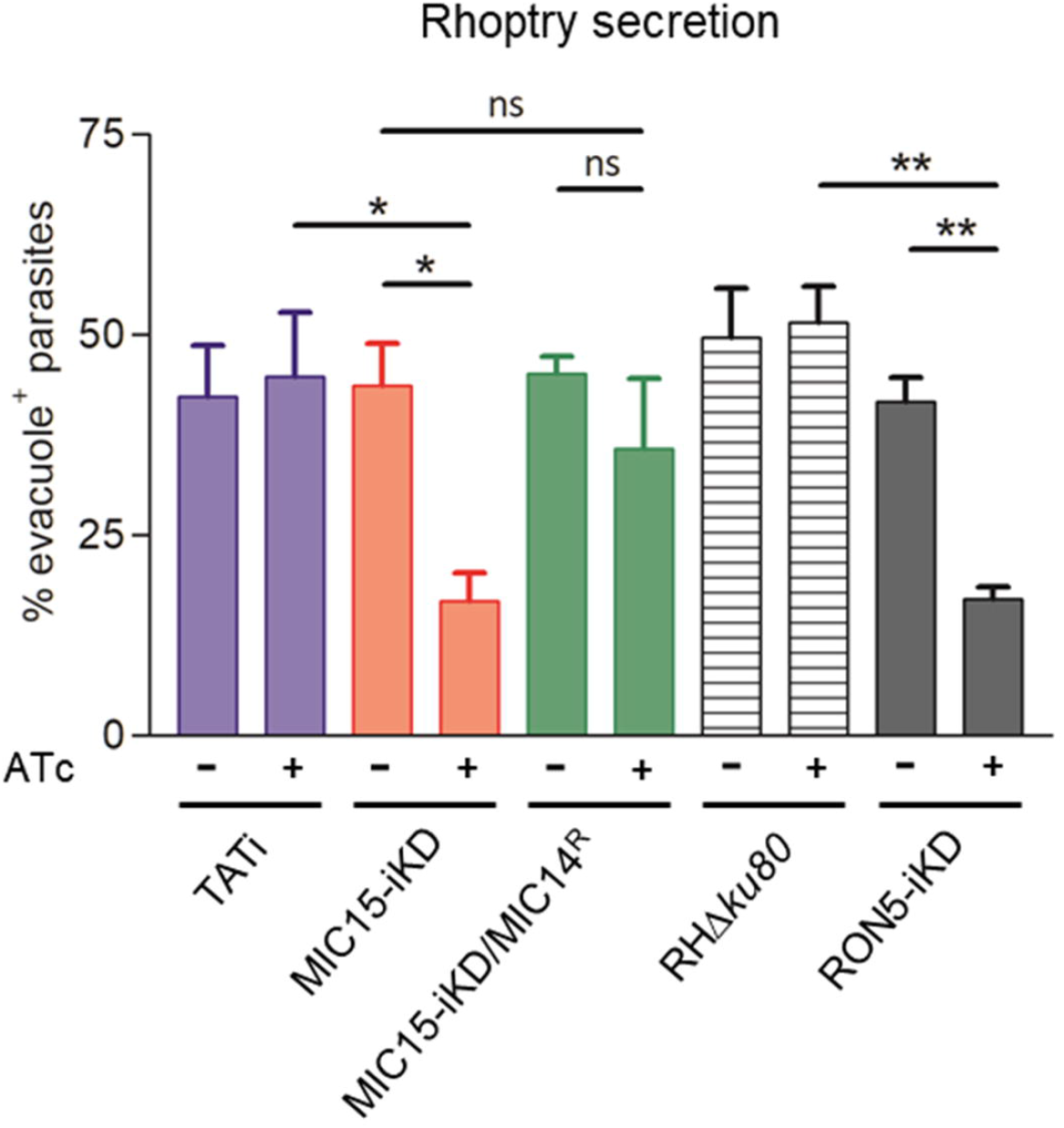
MIC15 and MIC14 are novel components of the rhoptry discharge pathway. Rhoptry secretion assay based on the detection of ROP1^+^ evacuoles released in the host cell by Cytochalasin D-arrested extracellular tachyzoites. Values represent the percentage of evacuole-associated tachyzoites and are expressed as means ± SEM from three independent experiments performed in triplicate. Statistical significance was determined by the Student’s two tailed *t*-test. *p<0.05, **p<0.01. The number of parasites able to produce evacuoles was significantly lower in MIC15-depleted tachyzoites (red bars) compared to the untreated MIC15-iKD control and the parental strain. No statistically significant reduction in evacuole formation was observed in MIC15-iKD/MIC14^R^ parasites depleted of MIC15 (green bars), demonstrating that MIC14 upregulation is able to functionally compensate the MIC15-dependent rhoptry secretion defect. The strain RON5-iKD defective in rhoptry secretion was used as positive control (grey bars). Based on this evidence, MIC15 and MIC14 are renamed rhoptry discharge factor 1 (RDF1) and rhoptry discharge factor 2 (RDF2), respectively.

## DISCUSSION

Apicomplexan parasites exhibit a lineage-specific expansion of genes encoding thrombospondin-related proteins (Naitza et al., 1998; Deng et al., 2002; Montenegro et al, 2020), whose functional role has been more widely investigated in *Plasmodium* spp. In the present work we explored the repertoire of *Toxoplasma gondii* adhesive proteins containing the TSP1 domain and showed that two novel members of this family, RDF1 (MIC15) and RDF2 (MIC14), are implicated in rhoptry discharge. While this process has remained poorly understood for decades, recent studies shed novel light on this crucial secretory event. According to the current ultrastructural model (Mageswaran et al., 2021), a couple of *T. gondii* rhoptries at a time are primed for secretion by docking their necks through the conoid and in proximity of an Apicomplexa-specific apical vesicle (AV) anchored to the cytosolic face of the PM. Four similar vesicles are aligned to the intraconoidal microtubules and are presumed to serve for successive rounds of secretion. The anchoring of the AV to the parasite PM was recently shown to be mediated by the rhoptry secretory apparatus, a highly elaborated structure described also in the distantly related apicomplexan *Cryptosporidium parvum*, that includes an apical rosette embedded in the parasite PM and electrondense proteinaceous material bridging the rosette to the AV. The rosette, which consists of eight peripheral particles and a central one resembling a pore, is the rhoptry secretion site and is evolutionarily conserved in the Alveolata superphylum, being implicated in the exocytosis of the extrusomes of free-living ciliates (Gubbels and Duraisingh, 2012).

Rhoptry discharge factors 1 and 2 add to the narrow group of *T. gondii* proteins so far linked to rhoptry exocytosis. Unlike microneme secretion, which can be induced by artificially raising parasite intracellular [Ca^2+^], rhoptry discharge strictly requires the contact with the host cell. Although the nature of the initial external trigger remains obscure, the micronemal proteins MIC8 (Kessler et al., 2008) and AMA1 (Mital et al., 2005) are know to be involved, as their downregulation significantly impairs rhoptry discharge. Within the parasite, RASP2, a member of the rhoptry apical surface proteins (RASPs), was shown to be essential for rhoptry secretion in both *T. gondii* and *Plasmodium falciparum* (Suarez et al., 2019). By locally enhancing phosphatidic acid and phosphoinositides concentration, RASP2 is presumed to be involved in the docking of the rhoptries and fusion of their membrane with that of the AV. A major advance in the definition of the molecular machinery governing rhoptry exocytosis was the recent identification of a new class of *T. gondii* proteins evolutionarily conserved throughout the Alveolata. Aquilini et al. [2021] identified *T. gondii* orthologs of Non-discharge (Nd) proteins known to be essential for trichocyst exocytosis and rosette formation in *Paramecium tetraurelia* (Froissard et al., 2004; Gogendeau et al., 2005). In *T. gondi*, Nd6 and Nd9 were shown to form a complex that includes five additional proteins, i.e, two other Nd molecules conserved in Ciliata named TgNdp1 and TgNdp2, the calcium-sensing protein TgFER2 (Coleman et al., 2018) belonging to the ferlin family and essential for rhoptry secretion, a putative GTPase and a Coccidia-specific hypothetical protein. Conditional depletion of each of the four TgNd proteins significantly reduced rhoptry secretion leading to severe invasion deficiency, that in Nd1, Nd2 and Nd9 iKD lines was paralleled by defective rosette assembly. In addition to sharing with Nd1, Nd9 and TgFER2 a similar dotted distribution throughout the tachyzoite cytoplasm, Nd2 and Nd6 were also shown to accumulate at the AV. In accordance with the functional characteristics of individual Nd complex partners, it has been suggested that calcium signalling and nucleotide binding/hydrolysis are instrumental to rosette assembly and/or rhoptry exocytosis (Aquilini et al., 2021).

In contrast to all TSP1 domain-containing proteins so far characterized in the Apicomplexa, RDF1 and RDF2 display an unprecedented intracellular localization and no evidence of trafficking to the micronemes. The punctate cytoplasmic distribution of RDF1 and RDF2 is strikingly similar to that reported for TgFER2 (Coleman et al., 2018) and individual TgNd proteins (Aquilini et al., 2021) and is suggestive of a highly dynamic situation. However, such a dispersed localization is not evocative of known vesicular trafficking routes or intracellular structures. The presence at the N-terminus of RDF1 and RDF2 of a typical leader peptide or a signal-anchor sequence, respectively, is predictive of protein targeting to the endoplasmic reticulum. Further research is needed to understand under which form and by which mechanism the two proteins are translocated to the parasite cytosol and if their trafficking is mediated by a not yet defined vesicular system. Our attempts to clarify this point by immunoelectron microscopy localization of RDF1 produced inconsistent results, as previously reported for TgFER2 (Coleman et al., 2018). The non micronemal localization of RDF1 and RDF2 shown in the present work is further supported by spatial hyperLOPIT (Localisation of Organelle Proteins by Isotopic Tagging) data (Barylyuk et al., 2020), which demonstrated that RDF1 is part of a separate minicluster including another member of the thrombospondin family (TGME49_277910) and two multipass TM proteins (TGGT1_261080, TGGT1_292020) showing homology to the *Plasmodium* Cysteine-Repeat Modular Proteins (Douradinha et al., 2011; Thompson et al., 2007). Interestingly, two recent studies deposited in a preprint repository (Singer et al., 2022; Sparvoli et al., 2022) showed that TGGT1_261080 and TGGT1_292020 depletion impairs rhoptry discharge and that these two molecules are part of a multiprotein complex also including RDF1. These results support a model in which RDF1 and RDF2 play a key role as Coccidia-specific components of a protein complex acting as a sensor of parasite-host cell contact leading to rhoptry exocytosis.

The characterization of RDF1 and RDF2 uncovered an example of *T. gondii* phenotypic plasticity related to rhoptry exocytosis. The functional redundancy of genes crucial for parasite survival is a strategy widely used by microorganism including the Apicomplexa, which have to face diverse environmental changes during their life cycles (Frénal and Soldati-Favre, 2015). In *P. falciparum*, extensive phenotypic plasticity was documented by the identification of hyerarchically organized invasion pathways relying on the EBA and PfRh redundant families of merozoite surface antigens (Baum et al., 2005). In *T. gondii*, several adaptive mechanisms have been described which are able to counteract mutations detrimental for parasite fitness, with most examples being related to host cell invasion. These include MyoA (Meissner et al., 2002), a major component of the subpellicular glidosome, that is responsible for the retrograde translocation of surface microneme adhesins and crucially involved in invasion and egress (Frénal et al., 2017). MyoA knockout and knockdown mutants were shown to possess a 16-25% residual invasion capability that was attributed to the compensatory action of MyoC, normally associated with a distinct glidosome localized to the basal polar ring. In MyoA-KO parasites, MyoC was shown to rapidly redistribute along the parasite inner membrane complex and to functionally rescue the loss of MyoA. A second compensatory mechanism was demonstrated in tachyzoites lacking GAP80, that is functionally replaced by the MyoA anchoring protein GAP45, in recruiting MyoC to the posterior glidosome (Frénal et al., 2014). A series of compensatory mechanisms enhance the capability of *T. gondii* to form a functional moving junction. This essential invasion-related structure depends on the interaction between the rhoptry neck protein RON2 and the micronemal AMA1 exposed on the host cell and parasite PM, respectively (Alexander et al., 2005; Besteiro et al., 2009). It was shown that AMA1 ablation by gene excision is compensated by the transcriptional upregulation of the RON2-binding paralog AMA2 (Bargieri et al., 2013; Poukchanski et al., 2013). Furthermore, the generation of a double knockout mutant AMA1-KO/AMA2-KO revealed that the divergent AMA1 homolog AMA4, which is abundantly expressed in the sporozoite stage, is upregulated in the tachyzoite along with its specific ligand RON2_L1_ (Lamarque et al., 2014), thus providing a second line compensatory mechanism preserving a molecular interaction of pivotal importance for parasite survival. To our knowledge, none of the molecular mechanisms underlying the mentioned compensatory systems has been elucidated.

Our results confirmed the expression of a *RDF2* mRNA form of □ 4.5 Kb consistent with the TGME49_218310 ToxoDB gene model. Furthermore, we showed that RH tachyzoites express a second and much longer *RDF2* transcript that could be detected only by amplification techniques. Importantly, this allowed to unveil the structure of the full-length *RDF2* gene, so far not identified by gene prediction algorhitms, and to demostrate that it harbors two promoter regions. In wild type tachyzoites, the ▢ 4.5 Kb mRNA transcribed from the downstream promoter is the dominating form. The predicted encoded protein of 968 amino acids was neither detected by anti-RDF2 polyclonal antibodies in Western blot, nor identified by mass-spectrometry in published *T. gondii* proteomic studies. This lack of proteomic data is likely to reflect very low expression levels of full-length RDF2 and the possible instability of the short RDF2 form, that in absence of an N-terminal leader sequence might be released in the cytoplasm and degraded.

The expression of a RDF2 form of □ 250 kDa predicted in the present study was demonstrated by Western blot detection, whose specificity was indicated by the disappearance of this band upon knocking out of the encoding gene in MIC15-iKD/MIC14^R^ parasites. The emergence of a variant strain in which full-length RDF2 upregulation partially compensated RDF1 depletion raised the question as to which mechanism is implicated in this rapid adaptation. We speculate that the mechanism regulating the activity the two *RDF2* promoters and accounting for the 12.7-fold increase of the long *RDF2* mRNA in MIC15-iKD/MIC14^R^ parasites might involve epigenetic control by specific antisense RNAs. Although their role in apicomplexan parasites remains largely unexplored, non coding RNA species are associated with more than 20% of *T. gondii* (Radke et al., 2005) and *Plasmodium* (Siegel et al., 2014) open reading frames. Our hypothesis is based on the observation that some *T. gondii* strains represented in ToxoDB show peaks of antisense transcription in the region encompassing the *RDF2* downstream promoter. While scarcely represented in strain RH, these *RDF2* antisense RNA(s) are more abundant in *T. gondii* strains showing a concomitant increase of sense RNA peaks indicating a higher degree of expression of the full lenght *RDF2* gene. A possible explanation is that the rate of transcription initiation at the *RDF2* downstream promoter is repressed by the binding of complementary antisense RNA(s), thus favouring the activity of the upstream promoter. Further experimental work will be required to explore the levels of the *RDF2* antisense transcripts in MIC15-iKD/MIC14^R^ parasites compared to the parental strain, and to characterize the RNA species possibly involved in the modulation of *RDF2* expression.

## CONCLUSION

In conclusion, our findings show the unprecedented involvement of two thrombospondin-related proteins in rhoptry exocytosis and expand current knowledge of the molecular players acting along the pathway that regulates this critical invasion-related process. In addition, the demonstration of structural/functional homology between RDF1 and RDF2 uncovers a further adaptation mechanism enhancing *T. gondii* phenotypic plasticity.

## DATA AVAILABILITY STATEMENT

The full-length sequence of MIC14 (RDF2) is available in GenBank with accession number AY089992. The full-length sequence of MIC15 (RDF1) is available in GenBank with accession number DQ459408.

## ACKNOWLEDGMENTS

We wish to thank Drs Vern B. Carruthers (University of Michigan, US), Dominique Soldati-Favre (University of Geneva, Switzerland) and Maryse Lebrun (University of Montpellier, France) for kindly providing critical reagents and for helpful discussion.

## AUTHOR CONTRIBUTIONS

FS designed the research. FS, AP, MDC, CN, ML, VM, FP, LT, SC, MF and LB carried out the research. FS and ML performed statistical analysis. FS and MDC supervised the project and wrote the manuscript. FS, MDC, AP and ML edited the manuscript. All authors approved the submitted version of the manuscript.

## ETHICS STATEMENT

Animals were housed and maintained at the Animal Care Unit of the Istituto Superiore di Sanità (ISS) in Italy according to D.L.gs. n. 26/2014 (art. 26 of D.L.gs.n.26/14). The in vivo protocol (n. 972/2017-PR; 11/12/2017) was approved by the Italian Ministry of Health.

