## Supplemental Fig 1 for "Functional characterization of the thrombospondin-related paralogous proteins rhoptry discharge factor 1 and 2 unveils phenotypic plasticity in *Toxoplasma gondii* rhoptry exocytosis"

MIC15 MVFRATREPFRLPLVAAFIALFLLKGVTCQLQLGTSTLQKNMWTTVKFDEPMIDPVVFVSPPEAPTNSFALVLIGGVTTSGFRARVYFPSCEYSPSGGTLYAVRWLAVEATSAGYYDSSN	120

```
MIC14                                                    MGARAVWSAGS--------DAGRT-----------------------WLLRAGPVCALLVLSLCSFLFLV    39
MIC15  RQTIDWTAHAVTLASSGTIPAGYVWSQSFSAWEDRPGTPGVLIGVQNHREMISAMYWSSEYLNPLVINSNSGGTLTFSFLRQISHRIGFTLKVGIFQYDARRGAVINGVNLVAQRYDLKV   240

MIC14  CK----SSAEARATTPVS------EPREATTVRENSETKPLSQGQNAAA----NPARRS-VSTSQALSFFDSMRPGPYKGRSSVYSNAGPDCYNLAATTLTVPKGLCTETCVLVWTDGKK   144
MIC15  NEVLPVADVVAGGTTPATFAVQWVTPTEAPL-ALVVKPKLLPDGSYSESLVAVEDCRNGATYEPHAFDVTHAFTGN--PASTAHRINGGPNCFRPSASVPTWLTEPCKLACESLWTANSD   357

MIC14  QYASCKDRVTCFLRDYTSRQVQAAQAGIPRKLSAAISRECSFVSKSPHLPLDQFQHMFILSASDVEHHGSWDAASACTAGQR-LDVAGALVGPKAVVAGTAAESVGCESINVTSHLKDIC   263
MIC15  DYWNCDDVNACFETDFRSAETLMALPQIDESIADVLVTSCNFATRTADL-IKQYEYAVNLKVDDIAIGTMWSTSSHCGGDKMEVEIVGAFYEKLGVS---ILTGSTCEVQDVSREVEALC   473

MIC14  AEAIKNPTERCIIEPSVYSNIISKF--SCNVKETPYLALYVTCIDVKVDDDVGCKVVTPASSQPDPEGGCTCPSGTVQCSLDESRLYGDVMSGIA--YGYTVHTKHNRGFKNGNVTA---   376
MIC15  KAE---GGIGCSISAAQIYELIMNYPGVCFALEDAELQIFFNCTFA-STSPFDCELYE-SKGFDQENGDCMCPNALAACTREEATFKSGWLQNLNKSYQYMLHGQIRYVAKSSVFQAFYE   588

MIC14  -------FQTPFCSGATVKVACKNAPDSASKKGEPATTAWLDRGPNCYKPTDSDNRQMCKFACLKLWNPADGALVN-----VNTPFDDFKDRIKSGRLTVMELTADGNAAIRSCDWAKRI   484
MIC15  APETSSAGDNTLCARMNTIVVCKN--------PTPVSAVVGNRGPNCYVIEH--GSQYCHLPCLTAWDAFLAAKDLPQQSDYQAEFERFWLSFDFPAIP-PHTSQSLLEQLQRCRFATRP   697

MIC14  RFAPAEQYEYETWTTSEA-----LSNFCPNSGTLQIVRAFLGCSYSLVDGNESQNCNYVDITKEVTKKCGSAWRSSKKGCTLGADELVAEADACETSCEGEWGIHLYYNCTPET--P-SA   596
MIC15  ATAPLQQYEDEIDVQLPDYGAVIIPLGCDKLHHREVVQALVGSASSIMLGVHEPGCRYEDATETVQRECKAAVSARKTTCEIDSSLFTSKDILCNTSTIK---LIIRGTCIPGPYLPTNY   814
MIC14  SCSLHQATTGLDETENEYACGCPYLAEMCDLEQAE-TSTVWKDTVRGYTVAILKGNLTYGLNGGYKRYHTTTV--SSDTCES-TGSRVLCKPPPASIELSPPNQTPDCTTSFSTKTGASS   712
MIC15  TCALYP-TTGVV-SRDGRSCTCPNSAYPCTYEEGQLTASRWKNEVNTGATVSLANNVVWTAPPGTSEYTYTHTDAYDYVCSTDEHHFVLCKDLT-PSALPLTEREPHCLNSFSVKPDAST   931

MIC14  SDDGLCVNECIIATQGECSKATNAWLCVVKKVSNCFIPNKAKSTCFIETEVGNYQAEFQSCSCPEASMPCTEEEVEATRYEWEPTFAPDNAYI--VVAPN-KVLG--PTKTIQPDYGMAA
MIC15  VDDRDCQDQCASLFATRCRQSFYKWLCVAQGLPECYIPNTDSFTCQIDTNPSAYNYSYGSCGCSGAYPPCSRNEANATRLDWITAFRALSHKKGVVVARGKQALWTENPDRWRYENGMLQ  1051

MIC14  EPGCGS-SRYKRVFCRGTSGNTSVTRYDATADCENARSLDASVISDFECRYGCSTRIKKCKAIMTQNPD--VHESEYACYTAQLNTIPKLRQCLVESKLVDPETGVGSMEAGFIVVTQEW   827
MIC15  GGGCLNNSWFVGVFCGAHLSVTPPARGDVLPLCSSAAKVENPDTPYPACRVLCAQVLNECKKVFSTTEGAAAYASVFDCFKARGK-GTTLDNCTYELGPENRATGVAELHTGWTVASASW  1170

MIC14  TSVAFKKEIENPV-VVTSLPATTNAVPVIQVKGVNSTGFKIRMKNDFCSVGF-ASPYTSVGWLASSQGTFLVSGAKSYVRVGTTLASL-------------NQPTTIRHLPRMNSAAPLV  1049
MIC15  SRVLFDVEFADPPVVFLGIPKDTQPFYTPSVRLVTKTSFEVKLYRNNCGLDDTRSSSAPVSWMAIPEGAYLTDFVENPVRVMKLPMTTRTPTVLTFSLSGLKEPEEMVALAQVQDVTPTS  1290
```

```
MIC14  FLQHEDSEGGSGNYIVASTVTASSASEATFKLTLVGS-GQPSEPIVIGYMIFDEIRQEYC-SLGCQLNGISMQTDYADSNERWTAESTDNVILAGQTVVYGSVVDPDGTSPVFVSKHDVA  1167
MIC15  VST-------DRVAVVIS-----NLKTNSLELSLVVDESVNFSTVAVGILISGKQNPDLAAGLSPLLNRYHLQT------FRIPAAAYSATSILGEGLYLS-----GGIMPHVFAAAIQT  1387

MIC14  WSSGNEIRSEGGV--------------ILDWRYAVPGGKDEHARSH-----------------PSGAQTSDSKTSKNDIGESLMQVSENLSVPGARSGWYPAIDIIYNSKCTVGSVSRTN  1256
MIC15  RSMTKELNESDRVVISRRATTTPFTWSALLWKKTCEAKHMFEAVENPSDLIIAGIYVEARKDEPSAILGNRHDLCLSFIGQSFESGVVATCMGACRSALPASVQLHCQDECAL------N  1501

MIC14  IFPAAGGRKNSVSRRLVALLYVSSGTSEISSSSFPPYGADCASAAGKKNYAAAECAAQCEAT------------VEGCTDS--DEDAWPCFFRQFPIDQKKKCYITAVAS------DLAA  1356
MIC15  IFAPSVH---------------------------TKCMTDCESLL---VAQYEECAGQCTATGTASCQRKCKQFLGNKCDSTASSDVAACLEFVTPPSVFEQCTLLVEYSVSEGGPSDGF  1591

MIC14  TSTTTAPPVDKDERCVLVDMSEAAYSAERGWDPETMTCKCPNDAPACSAEQATYNLINHSSVALGQGAICAAATGTVSGIARHFKRSTLDLCGD--LSKLSPLSWEQLPLPPYMADDVIL  1474
MIC15  EIEEETTPSQTYENCIIVDMRDERFTAFTGWDSRVASCKCPNDFRACTSETVNTQNHWRTELLASAGLCKEQPDGRFNAVTQQLWSTTGDLCALAADEQQPPISWASNMLPAYVDQDIVL  1711
```

```
MIC14  DDEVIRSGHYLCTKSYHHVFCPSSLITTSTTTPEPKWDWPARQDCVPGEWTSWSKCSMDCYVGSGALPVKSRSRSVLVDRRGTGNVCVLEDHTLCKVGSEVPYCEDLCWITDWGDWSGCE  1594
MIC15  DNSTIRSGDYQCREIWHYVVCPAAATSTTTQEPPVPPFTATLNTAIVGEWGEWSACTGTCFSQWW-TPKRTRTRLVLAELSHSQ-IPSVSE---TATCLDLPPCGTVCWEREWTEWSECK  1826
```

```
MIC14  QKVLQIGQAAVRARTKLRHIFMGTGGVCSSDH-IQYDYSGCK------------------------------------------------------------------------------  1635
MIC15  LFTIVMGQGLEYFRQQIKPVFDFVEEACGLDEHERYERCGEEQDNGETPATSTLQTDSLLETRSLTVSHGTAARALPRVSMDPALERPSFASLHGTRSHVPASTDAKIIERRKGIMSRRR  1946

MIC14  -----------------------------------------------------------------------------------SGMDSSGGREDDSAHSM------------------VE  1654
MIC15  SVPPSGYLEETAEQVAHGGESEQSGKASQNGSRRHRASRKQKRDLESIYSDASVRGSGESTLHGTGTNAYRDQIEWTSKSSSRSAIRDSGARGAEAGTSLLQKARRARRGAGRFRKSRSQ  2066

MIC14  QKSRVTDAGCKKSSGWSACSLPCQQPGANAVTQQFQLGLPETVTASTTSYCASRS---RACVEKKPPCVLDVAPDCSLVKPRYDSPEEALYCHELCNNALTRCREQAALQFQT----QWE  1767
```

MIC15 ARNQTPDKS**C**SVVSEWSG**C**DSP**C**LPHKG-STPRRYRLAVPGQNEA---NY**C**TVPLSGTEHLCTDLPQ**C**A-HPNFD**C**AKVTASRLTDEDIAA**C**KAV**C**VEVVKT**C**ETMMSAVFSLYTSLEQC 2181

```
MIC14  CIMAFINSSGVTGRCQIQKGAVSKTTLRPPVCFLSRTRQVDASKSLSFLTTDK---------------------------------------------TSGRGSCMCAVTNSVPCSPEEV  1842
MIC15  ALVQFQEYEQFPGQCHIPEEY--TSTRRPTKCFPSRTRRRVASGELPFIFQVSTASGTSPSATSDAASSVAESFVSTGTASAPSSRVMNALAFEASASQTSIDSCECYDPADEPCTAQEA  2299

MIC14  ALS-FDDWAKQLGSTVCPF--GQLGAFFNSPNSKGSADSFVYFGASDLGRVHCPISKSS-------DLGSRFPTFTEFESPESMNNFCENGLPFWQEKREPLVD-------CRQVIPKN-  1944
MIC15  RDSLFDSLYLFLTHPLCAESPAAEGALFTMSEPNQTADAASYFALRGLGRMHCPVPMYKSSTEVKGVLTPKRVTFSEFKTKEDLNEFCHKGLSNWRQNVPPEIIGSASFPNCMLVTPREG  2419
```

```
MIC14  ASTTDCPGRCHKAVVNCMV---SNDSLDNCINEALSAGDFAENCELQSSVKLGKGLMFCKRIRTDCEYSEWSEWSECSRTCRSGGDDEESVRVRGRKLLVAAEHG-GSCDADIEDSHGGS  2060
MIC15  AEPQDCALLCSETMSSCSAASLSFVELSQCVQDKLTESDFYSKCSAPEVLAPGEGIILCKKKVSTCDYTEWSEWSTCTATCFNWDEGVIPLRVRSRDFASSSADSRVLCRLE----SQND  2535
```

```
MIC14  LSDVQLCDWLPPCDALADTE----TYVIMPKPEPKLADWSATRTTTTTVIP---DLQNENKILCTIVDMAD-FSKSHRGYDTETESCKCPYNARVCSRTEAVNSRDNWDELMQTVCESNG  2172
MIC15  AIQTEKCDWMPVCPEAEGEEEDDATGGVEPRGEPIVPPWSPERPTDENNQAMGSEDIVSGTVECYVTNMGTIMTSYYRGYNQEYHGCNCPGGRRPCTRAEAVASLDLWTKDSGGLCDQDM  2655
```

```
MIC14  QGQILAQGMETFSCSDRVFRSFRGTLGQTPEAHCKSEDATYIFCEDGAIDNDLIFTEVMMSIITGVVMGIAVVYWAIQYSGDVQKVLGLAGRYVELTNLLHEVSQEPQAHEEEKAEEGEG  2292
MIC15  ATMISAAEGEAFFCATGSFGKIDTS---LSESSCASSEYQYVLCEGHPYEGIANLTTWVICLLLGVGGGICFVLSCVQYSSDIQKLLGLAGSYPVLVQNVTELQERESHKL-RRQGNISA  2771
```

```
MIC14  PDEDAEPEAAVEGNSEFDCNPRAKEENGEESEQDASDSESGSDDDDDIVSEE                                                                      2344
MIC15  TSERSDAHALSLSDSGWDVDGNQS--AGSAFPEEE-PWQFEDRDEEPLLSTRKYSRNLGSAGIPEPSEIGQTSPTQQRVPRASLAAQRSTCQLSSRLGQANSPEQSLERRSLKLRDSHAD  2888

MIC15  VELQRTLRKEMKGNQVWDKTV                                                                                                     2909
```

**Supplementary Figure 1.** Sequence homology between MIC14 and MIC15. The full length amino acid sequences of the two proteins were aligned with Clustal Omega. Conserved amino acids are boxed in yellow, invariable cysteine residues are highlighted in red. The main amino acids domains or elements are indicated by horizontal bars according the following color code: TSP1 domain (red), H-lectin domain (grey), transmembrane region (blue), leader peptide (green), signal-anchor sequence (orange).

**Supplementary Figure 1**
