## Supplemental Fig 2 for "Functional characterization of the thrombospondin-related paralogous proteins rhoptry discharge factor 1 and 2 unveils phenotypic plasticity in *Toxoplasma gondii* rhoptry exocytosis"

```
TgMIC14 ..IDNDLIFTEVMMSIITGVVMGIAVVYWAIQYSGDVQKVLGLAGRYVELTNLLHEVSQE...
NcMIC14 ..IDSGLIFTQLLMAFFTGVVLAFAVVYWAIQYSGDVQKVLGFTGRYVELSNLLNEVSEA...
HhMIC14 ..IDNDLIFTEVMMSMITGVVMGIAVVYWAIQYSGDVQKVLGLAGRYVELTNLLHEVSLE...
BbMIC14 ..IQNDTVMTHMTISLILGIVFGFIFIYWAVQYSLDIQKVLGITGRYVELSNMLEEVDKQ...
CcMIC14   ..RRQAAITKILILLVGIMLGVAVVLWFIQYSVDVQKVLGLRGRYVELVNMTKEESER...
CsMIC14 ..LQNDRTVLQFIVALVIGILLGLTFVYCALQYSVDIQQALGLAGRYVELSHMLQEVEEK...
TgMIC15  ..EGIANLTTWVICLLLGVGGGICFVLSCVQYSSDIQKLLGLAGSYPVLVQNVTELQER...
NcMIC15  ..EGITTLTTWLICVLLGMGGGVCFVLFCLQYSSDLQKLVGLAGSYPVLVQNVTELQAR...
HhMIC15  ..EGIANLTTWVICLLLGIGGGICFVLSCVQYSSDIQKLLGLAGSYPVLVQNVTELQER...
BbMIC15  ..WEEHTVTRWIVCLLLGLTLGIAVVLCCLQYSGDLQKLVGLAGSYPRLVQELVALETR...
CcMIC15  .IAAENATVFLITCLCLGAFTGIVVAYISLQYSIDLQKLVGLRGRYATISLELQQLREK...
EaMIC15  .EDAEHSLTFVIASLCLGVVAGLAVAYLSLQYSIDLQKLVGLRGRYVAATLELQQLQEM...
TgMIC2     ...SGIAGAIAGGVIGGLILLGAAGGA-SYHYYLSSSVGSPSAEIEYEADDGATKVV...
TgMIC6     ...SGHAGAIAGGVIGGLLLLSAAGAGVAYMRKSGSGGG---EEIEYERGIEAAEAS...
TgMIC8     ...RYSKGTIALVVVGCVALLGIIAGGISYARNRGGERDDEDLAPPPRSTRERRLSS...
TgMIC12    ...GVPVAAIAGGVVGGVLLIAGGAGAAVYASQGGVEAAEDEVMFESEEDGTQAGEN...
TgMIC16    ...NGVTYAVAGGIGLVVVIGLGFIGRKFYRALPAERTRDNYL
TgAMA1     ...NTALIAGLAVGGVLLLALLGGGCYFAKRLDRNKGVQAAHHEHEFQSDRGARKKR...
TgAMA2     .SVNIWLIAGPCIAAGVLLLGG-LIYWMAQ---RNKREPAVEKPQIVDETREHAVRT...
TgAMA3     ..GNTALIAGSVLGMLIILALVGTCVGFYYRKRP-------LPPTERPTVEASGGRE...
TgAMA4     ...SDYMAIYLAAGGVLLLLLAAGMAYGLRKRSTAAGGGGAMDFGVGEDGKPATAAE...
PfTRAP     ...YKIAGGIAGGLALLACAGLAYKFVVPGAATPYAGEPAPFDETLGEEDKDLDEPE...
```

**Supplementary Figure 2.** Comparison of the MIC14 and MIC15 transmembrane domains of *Toxoplasma gondii* and other Coccidia with those of characterized transmembrane microneme proteins. MIC14 and MIC15 amino acid sequences from other Coccidia are from *Neospora caninum*, *Hammondia hammondi*, *Besnoitia besnoiti*, *Cyclospora cayetanensis, Cystoisospora suis* and *Eimeria acervulina.* The amino acid alignment includes the C-terminal transmembranes of the micronemal proteins MIC2, MIC6, MIC8, MIC12, MIC16, AMA1, AMA2, AMA3 and AMA4 of *T. gondii* and TRAP of *Plasmodium falciparum*. The amino acid sequences of the transmembrane domains are shown in grey, characterized and predicted rhomboid cleavage sites in yellow. Invariable amino acid residues located in the MIC14 and MIC15 cytoplasmic tails immediately downstream of the transmembrane domain are highlighted in green.

**Supplementary Figure 2**
