## Supplementary figures and images for "Functional characterization of the thrombospondin-related paralogous proteins rhoptry discharge factor 1 and 2 unveils phenotypic plasticity in *Toxoplasma gondii* rhoptry exocytosis"

### Supplemental Fig 3

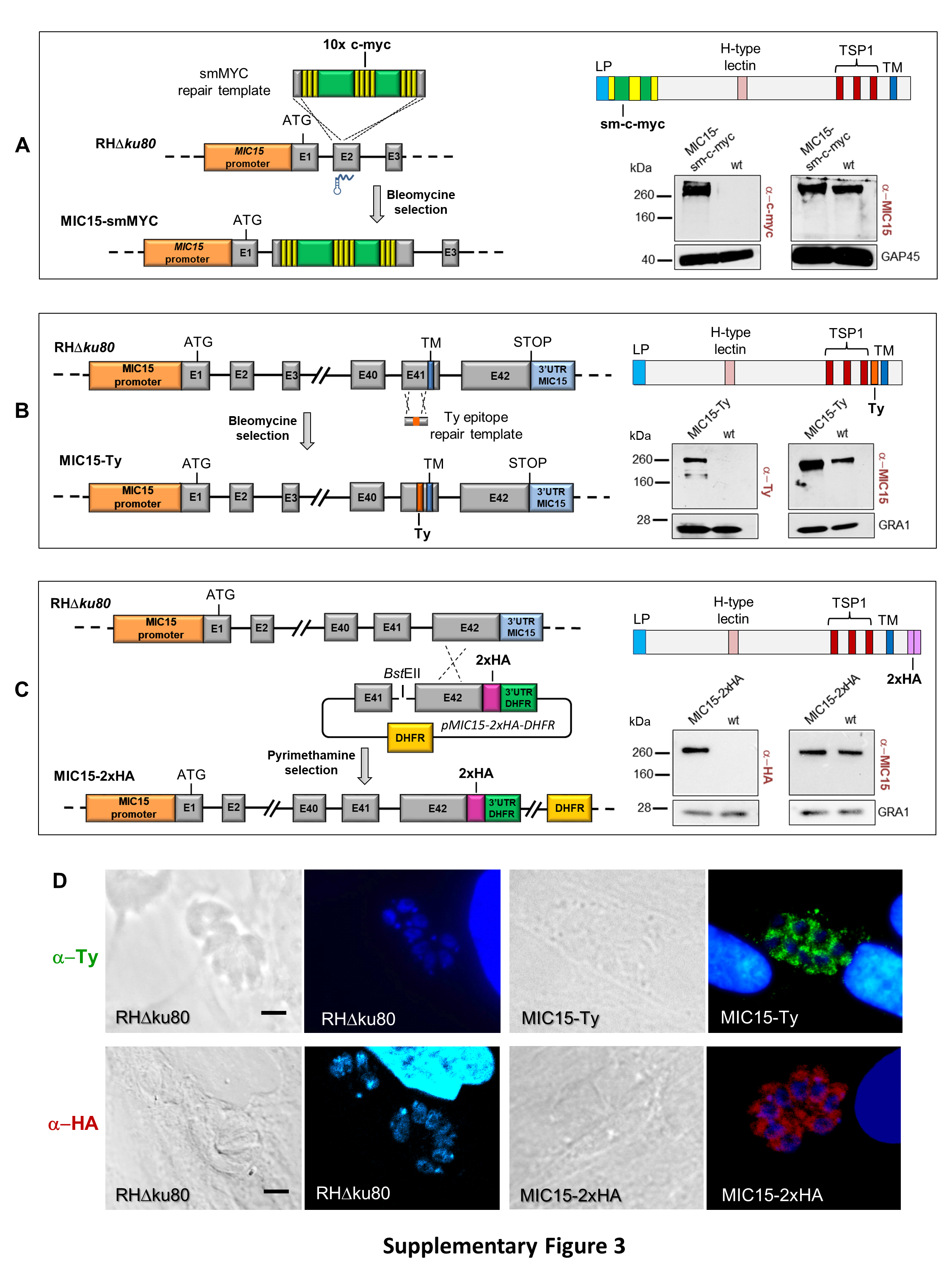

### Supplemental Fig 4

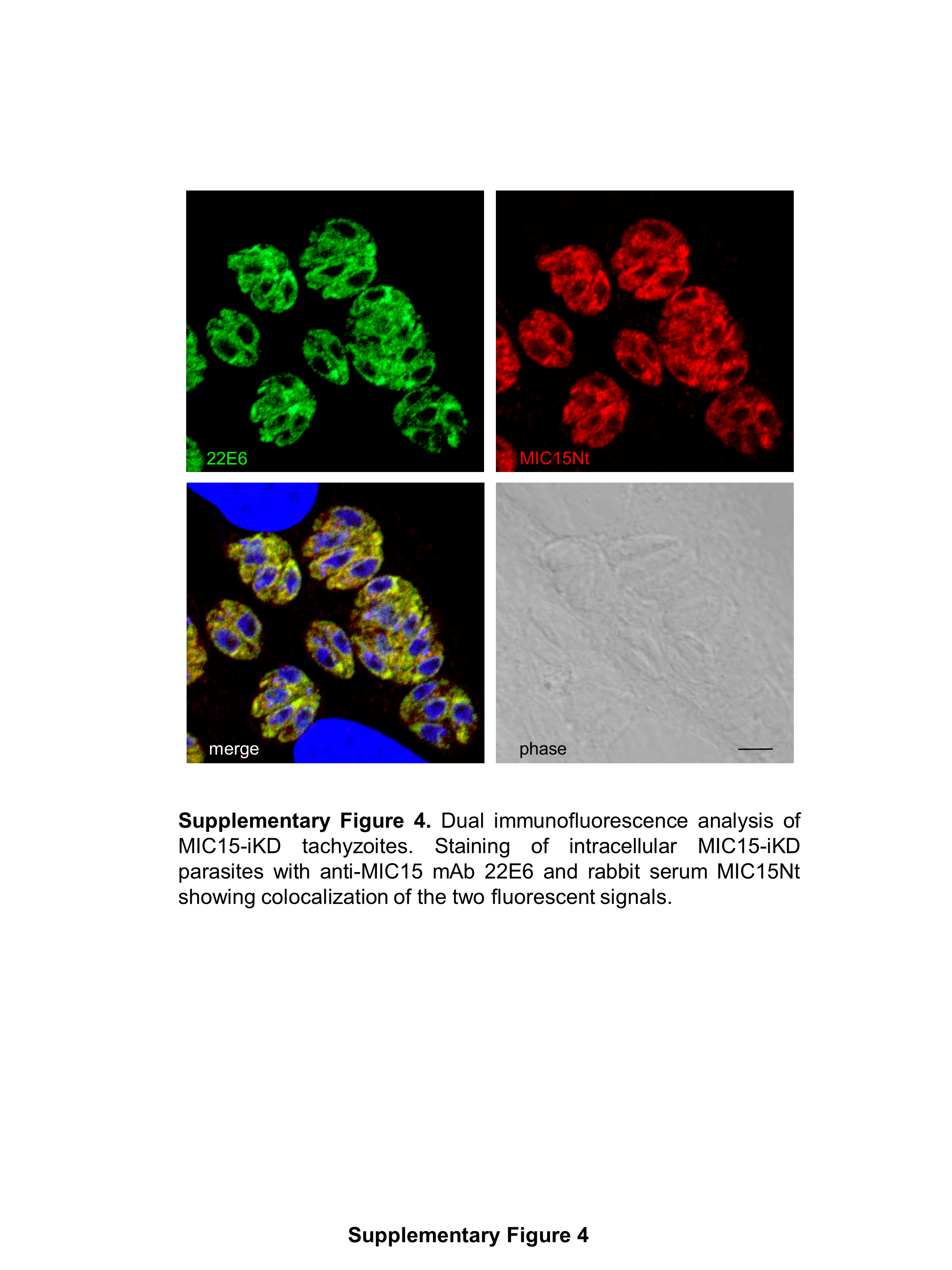

### Supplemental Fig 5

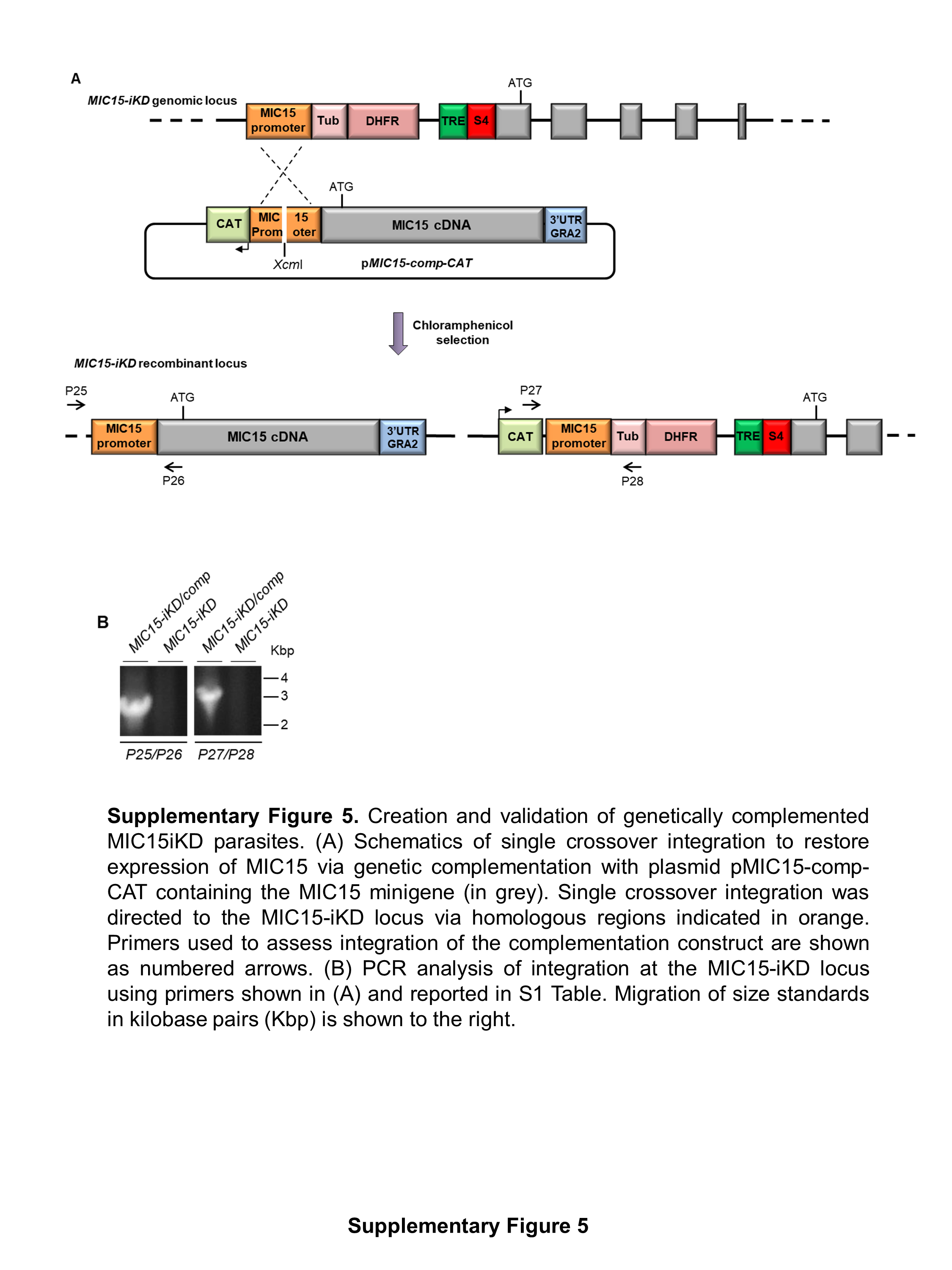

### Supplemental Fig 6

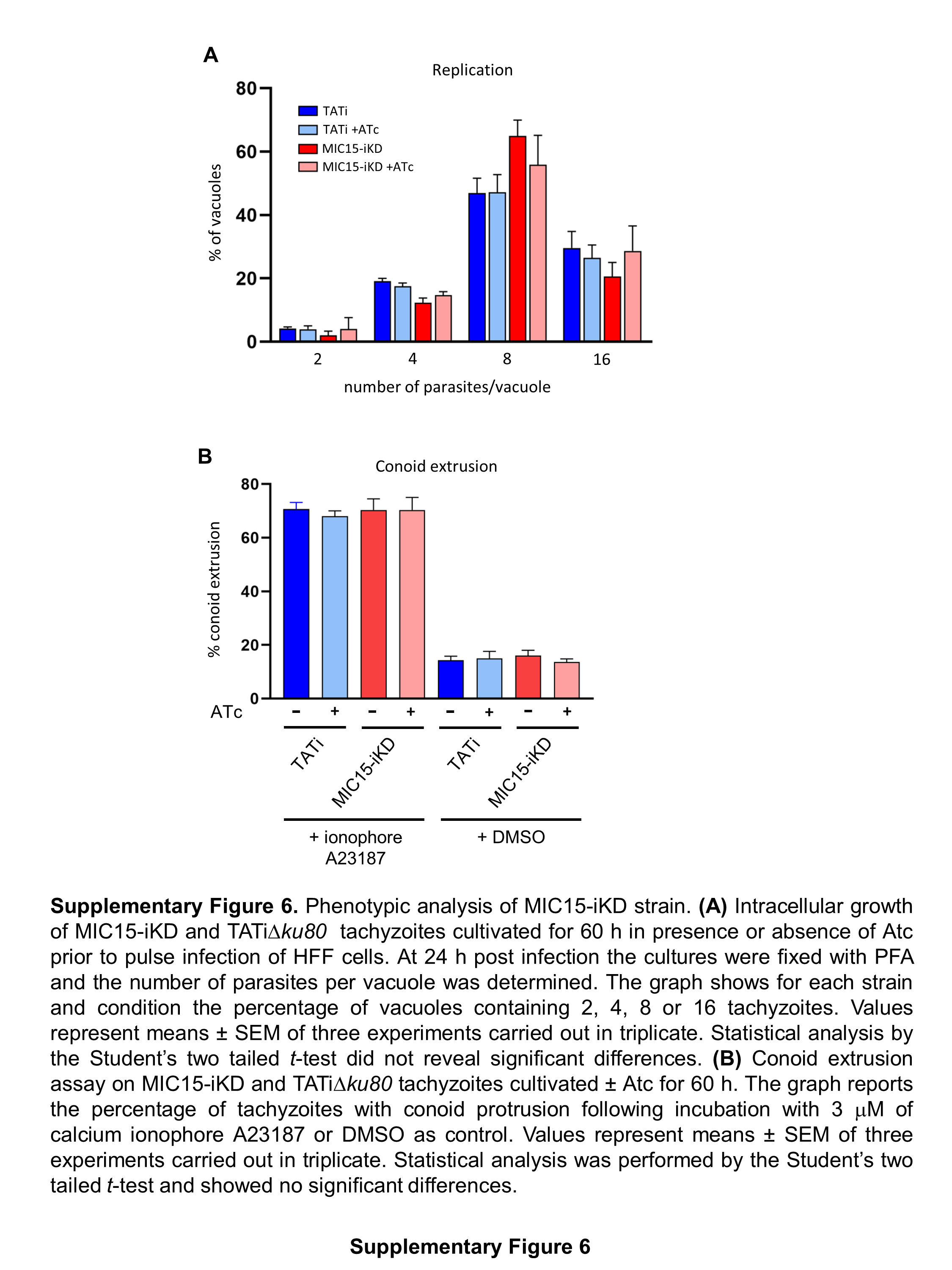

### Supplemental Fig 7

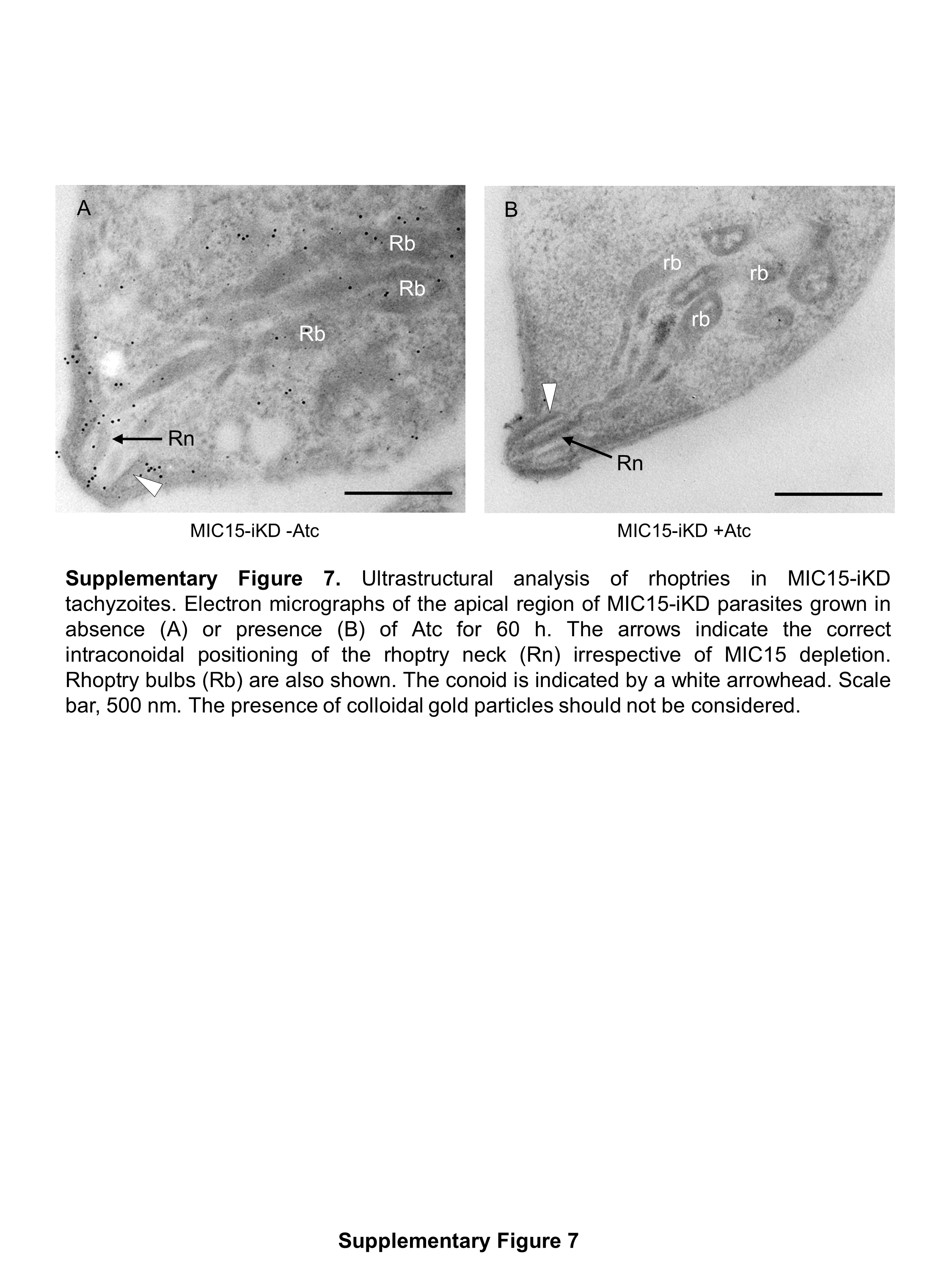
