## Supplementary material for "Functional characterization of the thrombospondin-related paralogous proteins rhoptry discharge factor 1 and 2 unveils phenotypic plasticity in *Toxoplasma gondii* rhoptry exocytosis": Caption to Supplemental Fig. 3

**Caption to Supplementary Figure 3**

**Supplementary Figure 3.** Endogenous tagging and localization of MIC15. **(A)** Schematic representation of three MIC15 variants in which the tags smMYC, Ty or HA were introduced at the N-terminus, upstream of the TMD or at the C-terminus, respectively. **(B)** Western blots showing recognition of the differently tagged MIC15 proteins by the relative anti-tag mAb and by the anti-MIC15 rabbit serum MIC15Nt. The dense granule protein GRA1 or the inner membrane complex protein GAP45 were used as loading controls. **(C)** Immunofluorescence localization of Ty- and HA-tagged MIC15 variants in intracellular tachyzoites stained with the corresponding anti-epitope mAb. For the localization of smMYC-tagged MIC15 see Figure 3D. Nuclei are stained with DAPI. Scale bars, 5 μm.
