## Supplemental Fig 8 for "Functional characterization of the thrombospondin-related paralogous proteins rhoptry discharge factor 1 and 2 unveils phenotypic plasticity in *Toxoplasma gondii* rhoptry exocytosis"

```
#MEGA
!Title MIC14 MIC15 alignment;
!Format
   DataType=Protein
   NSeqs=15 NSites=3764
   Identical=. Missing=? Indel=-;


!Domain=Data;
#BbMIC14 ---------- ---------- ---------- ---------- ---------M AGCGARRSGH GGRRSQLAPL PRLARILTFA
#BbMIC15 ---------- ---------- ---------- ------MGLL AVDDTINFAA PLHRT..LFV ATMALCKLSF SAVFLAFVVM
#CcMIC14 ---------- ---------- ---------- ---------- ---------- ---------- ---------- ----------
#CcMIC15 ---------- ---------- ---------- ---------- ---------. L.AAYCPIMF KKQ.RAKST- ----------
#CsMIC14 ---------- ---------- ---------- ---------- ---------- ---------- ---------- ----------
#CsMIC15 ---------- ---------- ---------- ---------- -----MHACH LFSR.CPLRC ..QVFGAVL. LAVVFGWQLV
#EaMIC14 MAGTYIADYS GYSGSAEHAD ALLSSSPTGS GAAWSGGAVE VSSVHYLAVH LSLSR.KMTR PKHLPSKMLQ LLIVAA..LL
#EaMIC15 ---------- ---------- ---------- ---------- ---------- ---------- ---------- MAVQ...HLV
#HhMIC14 ---------- ---------- ---------- ---------- ---------- ---------- ---------- ----------
#HhMIC15 ---------- ---------- ---------- ------MGDL RVS-----SF SLQQRTMVFR AT.EPFRL.. LAAFIA.FLL
#NcMIC14 ---------- ---------- ---------- ---------- ---------- --ME..AVWS T..DAGRTWF L.A.PACA.L
#NcMIC15 ---------- ---------- ---------- ------MGVF SVQ-----PF RLRTP-MVFT AT.APS.LY. VAAFVAAL.L
#SnMIC15 ---------- ---------- ---------- ---------- ---------. ..FRSAFVTV KSSSCCTSL. CV.VSL.GTV
#TgMIC14 ---------- ---------- ---------- ---------- ---------- --M...AVWS A.SDAGRTW. L.AGPVCALL
#TgMIC15 ---------- ---------- ---------- ---------- ---------- ------MVFR AT.EPFRL.. VAAFIA.FLL

#BbMIC14 --------VL LFACLGSG-- ---------- ---------- ---------- ---------- ---------- ----------
#BbMIC15 --------NG .ARQVQL.TT NLMSNK---- ----WTTVRF NEPMLKPLVF ASPPTLTA-N NFAMVLIGSV NKSSFEARVY
#CcMIC14 ---------- ---------- ---------- ---------- ---------- ---------- ---------- ----------
#CcMIC15 ---------- ---------- ---------- ---------- ---------- ---------- ---------- ----------
#CsMIC14 ---------- ---------- ---------- ---------- ---------- ---------- ---------- ----------
#CsMIC15 --------NG STPR.EV.DA ELATGK---- ----WTAIKF SRSMTEPVVF TSPPNVET-N NFGMVMIGDV KTTGFSARMY
#EaMIC14 LHHQLPSD.G A.PK.LCART DFVTGKIYLK GDGAWERFEF PNPLEKPLVF LGSPDYIG-D YPIEPQVGEV DANGFTARVM
#EaMIC15 --------AG .TY------- ---------- ---------- ---------- ---------- ---------- ----------
#HhMIC14 ---------- ---------- ---------- ---------- ---------- ---------- ---------- ----------
#HhMIC15 --------KG VTCQ.QL.TS TLRKTT---- ----WTTVKF DEPMIDPVVF VSPPEALT-N SFALVLIGGV TTSGFRARVY
#NcMIC14 --------I. ILCSFVC--- ---------- ---------- ---------- ---------- ---------- ----------
#NcMIC15 --------RG VTSQ.QL.VT ELTSTR---- ----WTTIQF EEPMIDPIVF TSPPETAG-N GFALVLLGAV TNSGFRARVY
#SnMIC15 --------SG AELK.DW.HV NARTDQ---- ----WVRVEF KGHIRRPCVF TSPPIIKGKD NYATVLIGDV SSSGFSLRIL
#TgMIC14 --------.. SLCSFLF--- ---------- ---------- ---------- ---------- ---------- ----------
#TgMIC15 --------KG VTCQ.QL.TS TLQKNM---- ----WTTVKF DEPMIDPVVF VSPPEAPT-N SFALVLIGGV TTSGFRARVY

#BbMIC14 ---------- ---------- ---------- ---------- ---------- ---------- ---------- ----------
#BbMIC15 FPRCARL-TT SSSESYPVHW LAVEATETGY YDYSTRQHVE WTAREYVIST AGGFAAGDLL SALFSAWQAH WGIPSVVLAV
#CcMIC14 ---------- ---------- ---------- ---------- ---------- ---------- ---------- ----------
#CcMIC15 ---------- ---------- ---------- ---------- ---------- ---------- ---------- ----------
#CsMIC14 ---------- ---------- ---------- ---------- ---------- ---------- ---------- ----------
#CsMIC15 FPSCASPGGT GSGTKFPVHW LAVESTSEGY FDGSTKQRVD WTAHVATISW SGTLAARTVR SNAAAVWENR -ATPNILIAV
#EaMIC14 VPSCITADLS RTWIGLP--- ---------Y IAWRKTAGRD YVADYVQLNS YSSISPGAVQ TYSYYVHDWD RISIQNMAYI
#EaMIC15 ---------- ---------- ---------- ---------- ---------- ---------- ---------- ----------
#HhMIC14 ---------- ---------- ---------- ---------- ---------- ---------- ---------- ----------
#HhMIC15 FPSCDH--SS SSGTLYAVRW LAVEATSAGY YDSSSRQTID WTAHAVTLGS SGTIPAGYVW SQSFSAWEDR PGTPGVLIGV
#NcMIC14 ---------- ---------- ---------- ---------- ---------- ---------- ---------- ----------
#NcMIC15 FPSCSR--AS NIGASYSVRW LAVDLTSAGY YDRSNKRTID WTAHKLTLAA PGTIPAGHVW STSFAAWQDR PGTPGVLVGV
#SnMIC15 KPHCIPQEQW EAEEPVRVAW LGVASTTEGH SKNTSTNNPH WMASVFTFSM SYDRAASKPW TISDKEWK-L PLEPKILLAV
#TgMIC14 ---------- ---------- ---------- ---------- ---------- ---------- ---------- ----------
#TgMIC15 FPSCEY--SP SGGTLYAVRW LAVEATSAGY YDSSNRQTID WTAHAVTLAS SGTIPAGYVW SQSFSAWEDR PGTPGVLIGV

#BbMIC14 ---------- ---------- ---------- ---------- ---------- ------AAAQ HSLLEGSQRA RQLRSPKASR
#BbMIC15 QNQREMMQEF YSEKEYAN-- ----PVVVHG HNGGILTFSF SRQISYRAYF SVKVG-IFLY VPSPHAAISG VNI.ARRYDW
#CcMIC14 ---------- ---------- ---------- ---------- ---------- ---------- ---------- ----------
#CcMIC15 ---------- ---------- ---------- ---------- ---------- ------.SLW DQSTHAAFAG DPAC.RSGHG
#CsMIC14 ---------- ---------- ---------- ---------- ---------- ---------- ---------- ----------
#CsMIC15 QNQKEMIQGL YSSSEYIN-- ----PLVVNK NEGEEVQFSF SRKISYRRDF KLNIG-ILRY DAVPGAIISG VH.QVRSYEL
#EaMIC14 EEKAAALNM- ---------- ---------- -TVNVYVRGS ISSRRKRYTI TLTTS-TTNN LPTVGVAIMG LGASV.THFG
#EaMIC15 ---------- ---------- ---------- ---------- ---------- ------LLLE LPAEVRAAFP DND.VRS---
#HhMIC14 ---------- ---------- ---------- ---------- ---------- ---------- ---------- ----------
#HhMIC15 QNHREMISAM YSSSEYLN-- ----SLVINS NSGGTLAFSF LRQISHRIGF TLKVG-IFQY DARRGAIING VN.VAQRYDL
#NcMIC14 ---------- ---------- ---------- ---------- ---------- ------LFRE S.AE.LATTP AAEATRATPV
#NcMIC15 QNHREMINAL YSSAEYLN-- ----PVVING KSGGSLMFSF PNQISYRSGF SLKVG-VFQY DAMPGATIGG VN.VAERYEF
#SnMIC15 QNQKEMLESG GFKEEDLTKP LSRIPSVNTE DTKDVVRFKL PARVAEDYSF ELKVAYLRYA DENGSAVVDG TL.K.KLYFL
#TgMIC14 ---------- ---------- ---------- ---------- ---------- ------LVCK S.AEARATTP VSEPREATTV
#TgMIC15 QNHREMISAM YWSSEYLN-- ----PLVINS NSGGTLTFSF LRQISHRIGF TLKVG-IFQY DARRGAVING VN.VAQRYDL

#BbMIC14 QTEAVLDPAL TRSSGTANSA RPGPWR---- ---------- ---------- ---------- ---------- ----------
#BbMIC15 KENEAAPLPE LVMD.SQP.P FVAQ.ATRSE TPLGVLLTQE RLADGTNPVS LVAVHDCRNT ASREAHEFEV TYAF------
#CcMIC14 ---------- ---------- ---------- ---------- ---------- ---------- ---------- ----------
#CcMIC15 DDSDADRTT. HVLG.Y.SVP AARY------ ---------- ---------- ---------- ---------- ----------
#CsMIC14 ---------- ---------- ---------- ---------- ---------- ---------- ---------- ----------
#CsMIC15 KANESTAI.S FD.TRAIPLL FAVQMATAAE APHSLSSRRP LSSSRNDGGS LIALEDCRES ATKQPHVFSV TSAS------
#EaMIC14 GVDIE.R.ST LIT..NGF.H SL.SPVLAFI LPSSVSASHS YAVDSNSNTS CVLRIISDCR TGAKLNDAAS SYLLIAHAQE
#EaMIC15 ---------- ---------- ---------- ---------- ---------- ---------- ---------- ----------
#HhMIC14 ---------- ---------- ---------- ---------- ---------- ---------- ---------- ----------
#HhMIC15 KVNE.WPV.D VVAG..TPAT FAVQ.VTPTE APLALVLRPE LLPDGTYSES LVAVEDCRKG ATHDPHTFDV THAFTGSPAS
#NcMIC14 TS.ESE--TK LL.Q.HNVA. IRARSSIPAS QALSFIDSLR PGPWSGQ--- ---------- ---------- ----------
#NcMIC15 RVNEK.PV.D VVAD..IPAT FAVQ.ATSNE APLALVLTPE SVPDGTDRES LVAVEDCRAA ATYELHRFDI THAF------
#SnMIC15 RSGETSVV.V SGRAFPKYGM FGVHVTVPRS FPAAVVLREV TKETGEVTTM VQGFENCVKI PPGRTSIFEA VLLLQP----
#TgMIC14 RENSETK.-- -L.Q.QNAA. N.ARRSVSTS QALSFFDSMR PGPYKGR--- ---------- ---------- ----------
#TgMIC15 KVNE..PV.D VVAG..TPAT FAVQ.VTPTE APLALVVKPK LLPDGSYSES LVAVEDCRNG ATYEPHAFDV THAFTGNPAS

#BbMIC14 GSKDSNHGPD CYALTKP--P GVVHKDLCTF VCKAMWNAN- QRSYLECDNA AECFNEDFAV ASRISQITS- -LPVVTTLYL
#BbMIC15 TAEHV.G..N .FWPSAK--V PPLLTEP.KL A.H.L.SLY- E..VWA.QDV NA..EHALRS IKTVEAVRE- -VDPSIAGHI
#CcMIC14 ------M..N .QDAKS---T LPNLLHT.QI A.TML.DEK- KED.VT.LDP M...TA..ST P.TLVSVLNN DVEFLLAQN.
#CcMIC15 AADYN.G..Q .KFTGTT--- --.IERQ.S. I.EEI.SKYA AERFMR.ADP MS..-TN.WN TTELAKEPQ- -VDSSILIKV
#CsMIC14 ---------- ---------- ---------- ---------- ---------- ---------- ---------- ----------
#CsMIC15 AALRV.G..N .RKPAAT--V PAMAAEG.KL L.ETL..KR- SSD.WA.EDV ....LK.MRS GDVVHTLAE- -ID.TMRDT.
#EaMIC14 S..HE.M..N .WNAQS---V SSDLLPI.EL T.GT..SQH- KEK.MK.P.P M....T..ST ..A.VEAWGD LFDDRLAGE.
#EaMIC15 AEDHE.G..Q .ARPSFA--- ---------- ---------- ---------- ---------- ---------- ----------
#HhMIC14 ---------- ---------- ---------- ---------- ---------- ---------- ---------- ----------
#HhMIC15 TTHRI.G..N .FRPSAS--V PTSLTEP.KL A.ESL.T..- SDD.WN..DV NA..EN..RS .ETLKALPQ- -IDESIAEGI
#NcMIC14 A.GYL.A... ..N.VTE--. AT.PRG...E T.VLV.EGG- .TRHAT.HDR EA..WRE.MT REVQAARAG- -I.RKFASQI
#NcMIC15 TVQRI.G..N .FRQSAS--V PSALTEP.KL A...L...K- SNEFWG.QDV ND..ENA.RS DET.KALPE- -VDASIAKGI
#SnMIC15 PIIRQ.G..K .REVEAADRM PGHAAVS..D L.LSL...K- SED.W..EDV QQ..LK..-- ---------- -ND.AVAH.F
#TgMIC14 S.VY..A... ..N.AAT--T LT.P.G...E T.VLV.TDG- KKQ.AS.KDR VT..LR.YTS RQVQAAQAG- -I.RKLSAAI
#TgMIC15 TAHRI.G..N .FRPSAS--V PTWLTEP.KL A.ESL.T..- SDD.WN..DV NA..ET..RS .ETLMALPQ- -IDESIADV.

#BbMIC14 STQCSYEPRS PHIAAGQFEH FIPLKLSTLG TRG-WDSTTV CPSNA-IVDV VGAYVGHPDV VEGRVARTYE CTYLNVTRKV
#BbMIC15 A.T.DF.I.D AV.ME-.YQY HVK.SADDVA AGTV.S.SSQ .SGDKSK.GI ...FYAKH.. SLMTGS---Y .EVQD.AHEI
#CcMIC14 ..N.------ ------SY.F ENVITW.DTV SGSI.STEEA .GKHN--IQ. .QLLFSDESI FN.KSTE.EA .KTQD..DLM
#CcMIC15 E.K.------ ------S.DG YQEYSAED.E SSAIFGGE.L .KNQK-ELK. .DFV....SA FR.DDW---N .LLYSQAE..
#CsMIC14 ---------- ---------- ---------- ---------- ---------- ---------- ---------- ----------
#CsMIC15 TSG.AF.T.V .DVES-.YDE .GT.READAE SGSV.S.DDH .ELGRSR.EI .S.LYMKRGA QFPAAL---- .EAID..S..
#EaMIC14 ..S..FAS.P .SSPEE.YTF EQLITW.DTA SGST.NTEVA .GKD.--.QL .QVLFSDEF. LN.KLPE.PS .GIQDI.GLM
#EaMIC15 ---------- ---PQE.YYG YQQFS.DQ.N SGVAFGGEAL .KEKE--LQ. .DFVS...SA FL.GGG---G .MLYSQAEE.
#HhMIC14 ---------- ---------- ---------- ---------- ---------- ---------- ---------- ----------
#HhMIC15 V.S..FAT.T A.LIK-.Y.Y AVN..VNDIV AGTM.STSSH .GGDKMKAEI ...FYEKLG. SILTGS---T .EVQD.S.E.
#NcMIC14 AKE.DF.TK. ..LPLA.... MFL.SATNVQ NNSR..VAA. .SDSQ-RLA. A..L..SK.. .R.VASEVDG .ESVD..SQI
#NcMIC15 V.S.IF.T.. ADLKQ-.Y.. SVN.NVDEIV AGTT.STSSH .GGEKLEAQI ...FYAKEG. SILTAS---P .EVQD.SKDT
#SnMIC15 GISDA----- ------ATRY T.T.R-KNSR E.LA.S.R.D .NGAHNYLI. RE.TY.K.VS .DKPQE---. .DRVD..EE.
#TgMIC14 .RE..FVSK. ..LPLD..Q. MFI.SA.DVE HH.S..AASA .TAGQ-RL.. A..L..PKA. .A.TA.ESVG .ESI...SHL
#TgMIC15 V.S.NFAT.T ADLIK-.Y.Y AVN..VDDIA IGTM.STSSH .GGDKME.EI ...FYEKLG. SILTGS---T .EVQD.S.E.

#BbMIC14 QRHCRNA-YK NPGVACVITR EDFRELQQNY SCISYTP--- PTLALFINCN LGPRVVQLTA KRWMLTKAFE TFRLGVGGAN
#BbMIC15 RQI.QGQ--- -Q.SG.SLGE PQVTA.SVLF TNLCFK.-SE AE.H..FS.- ---------- ---------- ----------
#CcMIC14 VSF.GEQ-VT STSG..KVSM LA.TNASSST L.PAAPN--- .SMLMRF..- ---------- ---------- ----------
#CcMIC15 AEA.QKDRKE .NRLY.ELGI Y.LPNAPELS TICTDV.SNE YRY...YD.- ---------- ---------- ----------
#CsMIC14 ---------- ---------- ---------- ---------- ---------- ---------- ---------- ----------
#CsMIC15 KSK.IAK--- -GTPG.ILGH S.LLDMAKEK .LACFSS-GE AE.HI.F..- ---------- ---------- ----------
#EaMIC14 DNF.QGQ-AS SDSA..NV.E DA.MAAALVT L.GAALN--- .SMLMRF..- ---------- ---------- ----------
#EaMIC15 ASA.QKDKEL .GRIY..LGM DELP.KPDLS DICDSP.SNY YQFV..YD.- ---------- ---------- ----------
#HhMIC14 ---------- ---------- ---------- ---------- ---------- ---------- ---------- ----------
#HhMIC15 EAL.KAQ--- -G.IG.S.ST AEID.ITTK. PGVCFA.-EG AE.QI.F..- ---------- ---------- ----------
#NcMIC14 KQL.AH.-IQ DSQQR...GP DV.S.VTSKF ..ATNET--- .Y...YV..- ---------- ---------- ----------
#NcMIC15 EKL.AEQ--- -S..G.SMSS AQIDQ.T.T. QGLCFA.-ED A..HI.F..- ---------- ---------- ----------
#SnMIC15 RNI.KK.--- --HPP.TFSD AE.DM.TKIA VNKCLV.HDD VE.RMYYD.- ---------- ---------- ----------
#TgMIC14 KDI.AE.-I. ..TER.I.EP SVYSNIISKF ..NVKET--- .Y...YVT.- ---------- ---------- ----------
#TgMIC15 EAL.KAE--- -G.IG.S.SA AQIY..IM.. PGVCFAL-ED AE.QI.F..- ---------- ---------- ----------

#BbMIC14 DAAPYSRSSP GFPRDCCVIS WVWNGTECGV SSVFDHRIRA EGQVDMMRGF LMQWLPNTTC IHFSQCRLAA IL-GGCYCRK
#BbMIC15 ---D.DTMAR F--------- ---------- ---------- ---------- --------E. KLYKSQ.YNP ES-.T.S.PS
#CcMIC14 ---------- ---------- ---------- ---------- ---------- ---------- ---------- ----------
#CcMIC15 ---------- ---------- ---------- ---------- ---------- ---------- ---------- ----------
#CsMIC14 ---------- ---------- ---------- ---------- ---------- ---------- ---------- ----------
#CsMIC15 ---------- ---------- ---------- ---------- ---------- ---------- ---------- ----------
#EaMIC14 ---------- ---------- ---------- ---------- ---------- ---------- ---------- ----------
#EaMIC15 ---------- ---------- ---------- ---------- ---------- ---------- ---------- ----------
#HhMIC14 ---------- ---------- ---------- ---------- ---------- ---------- ---------- ----------
#HhMIC15 ---T.AS... F--------- ---------- ---------- ---------- --------D. ELYESEGVDR EN-.D.M.PN
#NcMIC14 --V.VKVADE I--------- ---------- ---------- ---------- --------G. KLYTPATGLP SP-...E.PA
#NcMIC15 ---T.DATA. F--------- ---------- ---------- ---------- --------D. ELYESGGFDS ES-.A.M.P.
#SnMIC15 ---------- ---------- ---------- ---------- ---------- ---------- ---------- ----------
#TgMIC14 --IDVKVDDD V--------- ---------- ---------- ---------- --------G. KVVTPASSQP DPE...T.PS
#TgMIC15 ---TFAST.. F--------- ---------- ---------- ---------- --------D. ELYESKGFDQ EN-.D.M.PN

#BbMIC14 DMTLCSVEEA ILSPEIVRAS ERGGVIIARG NRGFSGGLAT AWLL------ ADCHKDRVQT ACRRTGTLLA GTEPETKWKN
#BbMIC15 ...A..LD.. KARDSWFHNL SPSHRY.LS. QIRLDPVEKV FQAYNAVDYA SNVAGKDLC. MSNTVSVCKN LRPYTATHL.
#CcMIC14 ---------- ---------- ---------- ---------- ---------- ---------- ---------- ----------
#CcMIC15 ---------- ---------- ---------- ---------- ---------- ---------- ---------- -------DPH
#CsMIC14 ---------- ---------- ---------- ---------- ---------- ---------- ---------- ----------
#CsMIC15 ---------- ---------- ---------- ---------- ---------- ---------- ---------- ----------
#EaMIC14 ---------- ---------- ---------- ---------- ---------- GSASGKPSCL .AHPLRSFYN H..ETRTVND
#EaMIC15 ---------- ---------- ---------- ---------- ---------- ---------- ---------- -------VPY
#HhMIC14 ---------- ---------- ---------- ---------- ---------- ---------- ---------- ----------
#HhMIC15 ALAA..R... TFKSSWLQNL NKSYHYMLH. QIRYVAKSSV FQAFYAAPET SSAGDNTLCA RTNTIVVCKN P.PVSAVVG.
#NcMIC14 GTEQ.VLK.. SMYT.VAIGV AS.FTVH.KN G...KN.DL. .FKS----PS CTGYSV..AC KKAPSS.SK- .DPAK.S.LD
#NcMIC15 AL.P..L..G SFRSSWLQNL N.SYQYTL.D HIRYMPKHSV FQAY----DA MNAGGQSLCA .TNTIAVCKN P.PAAALSG.
#SnMIC15 ---------- ---------- ---------- ---------- ---------- ---------- ---------- ------R.RS
#TgMIC14 GTVQ..LD.S R.YGDVMSGI AY.YTVHTKH ....KN.NV. .FQT----PF CSGATVK.AC KNAPDSASKK .EPAT.A.LD
#TgMIC15 ALAA.TR... TFKSGWLQNL NKSYQYMLH. QIRYVAKSSV FQAFYEAPET SSAGDNTLCA RMNTIVVCKN P.PVSAVVG.

#BbMIC14 RGPDCY---- KPLGSVSKQL CTYACLKLWN T---EGDYIR ATNPYQDFLE MVQEDSLQAL SEL-TYSGRM SIRDCTWEKR
#BbMIC15 M.A...---- --KE.SKS.Y ..MP..EA.S AFMASEMPLS THDYLRM.EV FWHSYTFPS. PKGTSSEFLE Q..N.RFSP.
#CcMIC14 ---------- ---------- ---------- ---------- ---------- ---------- ---------D AK.QT.FA.-
#CcMIC15 V..N..---- ---KGTDQPF .QVP.ITI.R AF---EAKFG VHDT.DQ.MS IWKNYDFTR. .FI-SS.AAE RVQA.IF...
#CsMIC14 ---------- ---------- ---------- ---------- ---------- ---------- ---------- ----------
#CsMIC15 ---------- ---------- ---------- ---------- ---------- ---------- ---------- -----LHGS.
#EaMIC14 G..Q.TNLSD QTVDTLEEAV .RNL..D..D MF--KDQA.A NPS..DE.VL LMRQWHADNP WA.-.STLSE IAIN..YDR.
#EaMIC15 S....F---- ---RGNDQPF .RIP..T..R QY---HRSHQ LRDS.DS.TS LWKTYDFSQ. PSV-DP.SIA RVKA..F...
#HhMIC14 ---------- ---------- ---------- ---------- ---------- ---------- ---------- ----------
#HhMIC15 ...N..---- --VIEHGSEY .HLP..AA.D AFLAAK.WPQ QSDYQAE.ER FWLSFDFPGI PSHTSQ.FLE QLQR.RFAT.
#NcMIC14 ......---- S.TD.NTA.M .KF....... PT--DDSLVN VHT.FD..MT RLSSGD.TVP EL--.DD.NT T..S.K.VDF
#NcMIC15 K..N..---- --IVERGVDY .RLP..EA.G AFVATN.VSE LSDCQSE.ER FWLSFNFP.I PS.TSS.FLD QLDY.RFAT.
#SnMIC15 Q....F---- -RVTKGDANF ..Q..FLI.S SFAA.A.KEN VPD.DAA.E. FWVKNIGSFF DNIL.PEN.Q RLVA.RFS..
#TgMIC14 ...N..---- ..TD.DNR.M .KF....... PA--D.ALVN VNT.FD..KD RIKSGR.TVM EL--.AD.NA A..S.D.A..
#TgMIC15 ...N..---- --VIEHGS.Y .HLP..TA.D AFLAAK.LPQ QSDYQAE.ER FWLSFDFP.I PPHTSQ.LLE QLQR.RFAT.

#BbMIC14 NRFDPPEQY- ------RY-- ---------- --------DT WST------- ---------- ---------- -TGTLEKLCP
#BbMIC15 PTSA.MD..- -EDEISVP-- ---------- --------VP S.G------- ---------- ---------- -SISVDIN.S
#CcMIC14 ---------- ---------- ---------- ---------- EG.------- ---------- ---------- -KFS.YNP..
#CcMIC15 WDYESST.SA SAFSCFVESV CLALAHACRG DPCIRRYG.P .CATRDLCSE FSTFVKK--- ------SSAT N.LR..TT.E
#CsMIC14 ---------- ---------- ---------- ---------- ---------- ---------- ---------- ----------
#CsMIC15 PGTA.R...- -EKEIHQE-- ---------- --------FG SLQ------- ---------- ---------- -SL.VSPG.N
#EaMIC14 AV.GSTDG-- ---------- ---------- ---------- ---------- ---------- ---------- -SFVVRD...
#EaMIC15 WEYEGAA.S- -EFYEIL.-- ---------- --------AG S.N------- ---------- ---------- -LLR.QNK.E
#HhMIC14 ---------- ---------- ---------- ---------- ---------- ---------- ---------- ----------
#HhMIC15 PATA.LQ..- -EDEIDVR-- ---------- --------LP DYG------- ---------- ---------- -AVLIPLG.D
#NcMIC14 IPRCSKKRHR QPYLLFVH-- ---------- --------LP DLGAPPVIVS SRGPVLRACV FQVKDSASIL PSNI.KR..T
#NcMIC15 PATA.LQ..- -EDEIDV.-- ---------- --------LP DFG------- ---------- ---------- -.VLIRLV.D
#SnMIC15 PYNVSI..S- EAGMFIPL-- ---------- --------HE DGS------- ---------- ---------- -PSVF.NF.D
#TgMIC14 I..A.A...- ------E.-- ---------- --------E. .T.------- ---------- ---------- -SEA.SNF..
#TgMIC15 PATA.LQ..- -EDEIDVQ-- ---------- --------LP DYG------- ---------- ---------- -AVIIPLG.D

#BbMIC14 SIGEPQLFQA LIGCSDTIRS GNVSSMCNYA DVTEKVRDTC GANFLNGS-E GCTLDLNALL PREEAC-KSC KGEWGIRLWY
#BbMIC15 NGLRRDVI.. .V.PKTL.TR NVEVLG.R.E ...DT..AA. DEARST.T-T A.VI.TSL.T .SAIL.---Y SS.VVLAVRS
#CcMIC14 DTYTTEIVA. VY..ESVLL. QGKKAA.EVK ...AVLNSV. SSLNTEDA-- S..IAEDL.P SKT.L.--PS CDTFTMMAY.
#CcMIC15 TYEQFS.VNV VS.S.S.L.E .KRVTG.AE. AL.DA.ERLL P.E.VK..LR TYEAS.EGIE .TFDT.---S ED.REFFVSW
#CsMIC14 -----MV.DE IA.THC.V-- -----G.RLN ..-------- ---------- ---------- ---------- ----------
#CsMIC15 PNEV.E.V.. .A.DRKATVL .YRTAG.R.E ...AS.ASL. ..A.AS..-V R.SVPVSS.P TKA.L.--PR FTSV.LTVRT
#EaMIC14 DSRFADIVG. SY..NSV.VD .ELK.A.QVH .A.SV.QQA. ----TSRTLT E..VFG.QFP .TL.L.--PS CDAFSLVVF.
#EaMIC15 NYERFS.V.V VA.PRELLEN .RSITG.AAM .I.DS..SSL PEA.GT.DVR NYAVS.SG.K TTLDT.---T EAGASYFVSW
#HhMIC14 ---------- ---------- ---------- ---------- ---------- ---------- ---------- ----------
#HhMIC15 KLHRREVI.. .V.SASS.VL .VHEPG.R.E .A..T.QRE. E.AVSARK-T T.EI.SSLFT SKDIL.--NT -STIKLIIRG
#NcMIC14 DSRK..IVR. FV....SL.Y .KE.NN...L .I.NELTMR. .SA.T..T-A ....HPEE.. SEDD..Q... TDQ...Q.Y.
#NcMIC15 .LYV.EVV.. ...PAS.LL. SVHEPG.R.E ....T..KA. QNDWSR.A-T T.AI.VSLFT .K.IL.--.T -STVKLVVRG
#SnMIC15 NS.ALEIVDV VV.PKVWVMN .EAVEG.STL NM..F.KRF. HQPIKD.E-R K.VIPRSL.P TK.QL.---A TEMC..F.R.
#TgMIC14 NS.TL.IVR. FL...YSLVD ..E.QN...V .I.KE.TKK. .SAWRSSK-K ....GADE.V AEAD..ET.. E.....H.Y.
#TgMIC15 KLHHREVV.. .V.SASS.ML .VHEPG.R.E .A..T.QRE. K.AVSARK-T T.EI.SSLFT SKDIL.--NT -STIKLIIRG

#BbMIC14 NCSPGI---- --DPSKVRCR VQKAIPSSNQ EDS--LCRCA SMGDPCT-MD QAEADETWKS TVEMQTSVLL ADDVSYTYYT
#BbMIC15 ..V.DP---- -SLKPDFE.E PYLVTGRI.T NGT--A.T.P NNA....ITE SRLS.SW..- .LPTGSI.S. .NA.IWAAPP
#CcMIC14 ..RISE---- -NTLAGYDIF THSSSG.LTY SGDIPS.G.V DGST.A.-EE EVV.KSS..A YLTPPAT.R. GGAR.LFVDK
#CcMIC15 ..AR.AHDSA SE.KLPFQ.T LYR.QGKQDA THH--S.A.P NSAT..D-.. A.LVSRS.RE H.SEGAT.S. SNYLVFKPSK
#CsMIC14 ---------- ---------- ---------- ---------- ---------- ---------- ---------- ----------
#CsMIC15 R.I.EP---- ---------- .FQGAGAADS VPRE------ ---------- ---------- ---------- ----------
#EaMIC14 ..KLRL---- -ESLGGYGII PHALSGNIVY SGNIPS.S.. EGSV.A.-QE EIT.LTS.QA LITPPE.FV. SGARGVSVDE
#EaMIC15 S.V.NV--TM ASESVPFS.N IYPVMTK-A. DTNFHS.A.P NSAKT..-AN A.AVSRS.IA NTPQSAT.S. ..YLV.KTPE
#HhMIC14 ---------- ---------- ---------- ---------- ---------- ---------- ---------- ----------
#HhMIC15 T.I..P---- -NL.TDYT.A LYHTTGVVSR DGT--S.T.P NSA....YDE GQLTAS...N E.NTGAT.S. .NS.VW.APP
#NcMIC14 S.TLET---- ----.TSL.S LNQTMTGLDE SENEYA.G.P YLA.M.D-LE E...NDS..D ..RGF.VAI. KGNLT.GLSG
#NcMIC15 ..I..P---- -SL.TDYT.V PYRTTGTVAR NGK--S.S.P NSAN...HEE GRLTAS..Q. EFKSGVT.S. ..R.IW.APP
#SnMIC15 ..VG.S---- --Q.DNYS.E ALPTTGEFTS .----S.T.P NSA....YFE ARNSEKY.TQ D.PEYAT.AF -GSHGVRLPR
#TgMIC14 ..T.ET---- ----PSAS.S LHQ.TTGLDE TENEYA.G.P YLAEM.D-LE ...TSTV..D ..RGY.VAI. KGNLT.GLNG
#TgMIC15 T.I..P---- -YL.TNYT.A LYPTTGVVSR DGR--S.T.P NSAY...YEE GQLTASR..N E.NTGAT.S. .NN.VW.APP

#BbMIC14 RGR--FYRQS AIRGNTCQSS -WLRVLCNEP PSQISLLPPL KVGPTSASAS AGGRSSTRCT SKAAGGFFRA AHRMGWLANC
#BbMIC15 GTSLYGFTNT DAHRDI.APD PFHY...KD- ---------- ---.------ ---------- ---------- --------..
#CcMIC14 N.KLE.SIWA PSSDTD.LDD -DVQ.F.KR. SQTA.DEGMM NPT.------ ---------- ---------- --------L.
#CcMIC15 A.LLSVSLME EL-STP.AEE -SSY...EV. ---------- ---------- ---------- ---------- --------S.
#CsMIC14 ---------- ---------- ---------- ---------- ---------- ---------- ---------- ----------
#CsMIC15 ---------- ---------- ---------- ---------- ---.------ ---------- ---------- --------D.
#EaMIC14 D.ELT.TIWA PSSEYD.L.E -FVKI..KH. VE.DPDEGMM NPT.------ ---------- ---------- --------L.
#EaMIC15 S.TGSLVLMA DL-.TV.GDT GSSY...EG- ---------- ---.------ ---------- ---------- --------S.
#HhMIC14 ---------- ---------- ---------- ---------- ---------- ---------- ---------- ----------
#HhMIC15 GTSEYT.THT DAHDYV.STE EHHF...KDL SPSALP.TGR E--.------ ---------- ---------- --------H.
#NcMIC14 GSKRYHS--T S.NSY....P -TT....KP. .AT.EIV..S Q.-.------ ---------- ---------- --------D.
#NcMIC15 GT.DYA.THT DAQDYV.SAN EHHF...KDR MP.EVTTSSR N--.------ ---------- ---------- --------H.
#SnMIC15 P.SQVV.TL. DRHELY..A. SNNF...KAA TVVVNTPVVR RE-.------ ---------- ---------- --------E.
#TgMIC14 GYKRYHT--T TVSSD..E.T -GS....KP. .AS.E.S..N QT-.------ ---------- ---------- --------D.
#TgMIC15 GTSEYT.THT DAYDYV.STD EHHF...KDL TPSALP.TER E--.------ ---------- ---------- --------H.

#BbMIC14 KKAFSVESDA TSVDDAVCIE ECQKMLQTEC ADSANSWLCA ARSL-THCYI PNVAKTTCLV TKDV-GPFVE AFKSCSC--A
#BbMIC15 RE..AIDMH. S.A..RE.Q. A.AALVVSG. .TAQSK...V ..A.-SK..T ..TESM..QI DMSP-YT.SA .DGT.R.--T
#CcMIC14 EE.Y.TKVSS GTFQHEL.R. Y.AHL.KNT. .SAE.K.A.V FENS-.N... ..STR...AI RAVS-AEYNP S.N..G.END
#CcMIC15 SS...TVEG. SAA..EE.KT Q.SQKAGGL. .A...I...I ..Q.HAE... ..RNSL..YI ESP--..EDA SHGT.A.-ST
#CsMIC14 ---------- ---------- ---------- ---------- ---------- ---------- ---------- ----------
#CsMIC15 AN.Y.TV.GV S.AA.TA.Q. D.AAV.AST. KEDT.K.I.V .TA.-PT.AV ..ADRI..NI ETS.-.DYD. QSHT.G.--P
#EaMIC14 EES..TKPGK R.SH.EL.R. T.SHV.KNK. SS...K.A.I NDNI-.D..V ..SLRK..GI RGIS-PQYND KYRT.G.-DD
#EaMIC15 GR...TLEG. .EE..LE.RK A.TAKSSG.. GSTR.I...V .L.I-QG..V ..TSSLA.HI ELP--..EDA YNGT...-PS
#HhMIC14 ---------- ---------- ---------- ---------- ---------- ---------- ---------- ----------
#HhMIC15 LNS....P.. ST...RD.QD Q.ASLFAGR. GQAFYK...V .KG.-PK... ..TDSF..QI DTNP-SAYNS SYG..G.--S
#NcMIC14 NSS..IKKG. ..S..SL.VN ..SIVM.RT. .SA..V...V GKT.-PN.FV ..R.....VI ESS.-.AYQA S.Q....--P
#NcMIC15 LRG..SQPA. .T...RD.QD A.ASLFA.S. .QAK.R...V .QA.-PK... ..TDSF..Q. DTNP-SVYST SYGA...--S
#SnMIC15 SR.HL.QKEQ ATGS.VQ.QF A.ISAAKDQ. .SAE.P.T.V .EA.-PA.M. ...DQV..HI .SP.A.LYDA DAQT.G.---
#TgMIC14 TTS..TKTG. S.S..GL.VN ..IIAT.G.. SKAT.A...V VKKV-SN.F. ..K..S..FI ETE.-.NYQA E.Q....--P
#TgMIC15 LNS...KP.. ST...RD.QD Q.ASLFA.R. RQ.FYK...V .QG.-PE... ..TDSF..QI DTNP-SAYNY SYG..G.--S

#BbMIC14 DGSMPCTTEE VAATQYEWEK IFTA-----S DAYAVLSGQR VV--GPSKAI QADYGMGTSH GCG-NPRYKR VFCRGTSTVT
#BbMIC15 GMGS...VD. AEV.RPD.IE S.Q.LGVGDR EGVV.AKNF. ALWI.DPDKW KYEA.VNKGA ..VA..WIAG ...GVKLAIK
#CcMIC14 PTME...ES. .KMLSVD.TS D.MEHYQG-K EGLLL.RNM. .M--DGTGS. YT.F.VSQNL A.TRASK..G I....NEI--
#CcMIC15 SSLGA..SL. AEV.HFV.QE SLSQFWDG-. EGVI.AKNN. ALWPSAKASW HYEA.-SQAG L.EQKRWI.G ...L.SLA--
#CsMIC14 ---------- ---------- ---------- ---------- ---------- ---------- ---------- ----------
#CsMIC15 ESHP..SV.. .E..R.D..S S.KS--AGST GGVV.AK.R. ALWLSD.SRW RYEN.FTQDG ...R..WIVG ...KAVG...
#EaMIC14 PTME...ES. .KMLAHD.MG Y.KDHYQG-K GGLIL.RNMG .M--ED.GT. YT.F.VA.-- ..TRSSE..G I....-----
#EaMIC15 SS.G...FL. AD..A.A.KS SLAQFWRG-A NGVIMAKYN. ALWPSTPASW RYEA.-HRYD Q.EKRTWIRG ...M.-----
#HhMIC14 ---------- ---------- ---------- ---------- ---------- ---------- ---------- ----------
#HhMIC15 GAYP..SRH. AI..KLD.IT A.S.L-SD-K KGVV.AR.KQ ALWTENPDRW RYEN..LQGS ..LN.SWFLG ...GAHLS..
#NcMIC14 NA.....ED. .EV.R....Q T..P-----E E.FV.VAP.K .L--..G.T. .P....AGEP ...-SS.... .....A.GK.
#NcMIC15 GTYS..SA.. AEV.KHD.LT A...L-SG-R KGVV.AR.KQ ALWI.NPDKW RYEN..NQGS ..LN..WFVG ...GARLS..
#SnMIC15 PD.K..DVAS .R..THA.MP D.ETY-SG-R PGVI.AASRQ ALWTNNPGRF YYEN..HK.G ...E..WIRG ...ESKADAG
#TgMIC14 EA.....E.. .E..R....P T.AP-----D N..I.VAPNK .L--..T.T. .P....AAEP ...-SS.... .......GN.
#TgMIC15 GAYP..SRN. AN..RLD.IT A.R.L-SH-K KGVV.AR.KQ ALWTENPDRW RYEN..LQGG ..LN.SWFVG ...GAHLS..

#BbMIC14 GVVRYDETAD CS-------- ---------- KVESLYTELT SNFECRFGCG AVTKKCK--T I---MKQNPN LYAS-EF--L
#BbMIC15 PPA.G.AAPL .A-------- ---------- EAVTDDPAKA TSSA..VH.A S.F.E..--Q .---VS.GSP I.SV-AL--Q
#CcMIC14 ----GRQ?PN .D-------- ---------- GAV.TDLSNV .DYQ.AYS.Y KIYNQ.R--- -----E..DT SI..------
#CcMIC15 --------P. ..-------- ---------- TAVTTNPQKA .TLY.QYY.. H.EWE.RVLM .-----.K.T ..KD-IM--E
#CsMIC14 ---------- ---------- ---------- ---------- ---------- ---------- ---------- ----------
#CsMIC15 LP..GG.CCF S.QLKLWQVL GMQMQKKPVG TIGAPVDPCL VSLSVQAFVS .LPRTASSIL SSRGVSSKSP CNRR-G.CMP
#EaMIC14 ------LQPS .D-------- ---------- GTI..NKNDV .DYQ.AYW.H E.HST..--- ------ETQP SN..-----A
#EaMIC15 -----SLAP. ..-------- ---------- TAAT.NHWKA TTSY.KYL.. NIEWR.R--F T---.VD..T ..KD-LY--Q
#HhMIC14 ---------- ---------- ---------- ---------- ---------- ---------- ---------- ----------
#HhMIC15 PPA.R.VLPL ..-------- ---------- SAAKVEDPS. PYPA..VL.A QLL.E..KIF S---TTEGAS A...-V.--D
#NcMIC14 ..T....... .E-------- ---------- NAT..DSAVI .DS..HYA.S I.I....AT- ----I.K..D A.E.-.Y--A
#NcMIC15 PPA.G.ALPV .A-------- ---------- GAAMTADPGS PHPL..VT.S KLLEE..KLL S---SDGGES S.P.-V.--G
#SnMIC15 ASNEG.GIPL .--------- ---------- NTAVSQNP.F T.A...IR.H TTLEE.RNEY E---ELGKST SVEEA.YVGA
#TgMIC14 S.T...A... .E-------- ---------- NAR..DASVI .D....Y..S TRI....--A .---.T...D VHE.-.Y--A
#TgMIC15 PPA.G.VLPL ..-------- ---------- SAAKVENPD. PYPA..VL.A Q.LNE..KVF S---TTEGAA A...-V.--D

#BbMIC14 CYQEQVST-- ---------- ----VPGMEN C-----SIHR K-----LVDP DIGIGNMEGG YVVAEQTWTQ VTFKNHI-DN
#BbMIC15 .FEAAAAA-- ---------- ----H.TLMF .-----.YQI A-----PE.. .A.KATLDA. WT..TTE.VR ...GADFSEP
#CcMIC14 ..STAQKD-- ---------- ----D.TLRQ .SVPLGKYEH GQSAGVVR.G F.AES..LT. V.AITTE.IK .A.PSLVFP.
#CcMIC15 .FKQLQTE-- ---------- -----SALKY .-----ELDT T--------- ----A..AVS WM..SG.ES. .L.AEP.YPP
#CsMIC14 -------G-- ---------- ----LQTINK S--------- ---------- ---------- ---VAGE..E AKTETLVLS.
#CsMIC15 .FSREQLSKT VFMRVQVRGV QEKFSKPFSF .FAFGFRVCT V-----PKET .P.E.HL.A. WLA.SGS.HK .A.STAFSTL
#EaMIC14 .LY.TQK.-- ---------- ----HSILKQ .-----.VPS E-----.E.K EQQDS..VA. TAIVTTE.KA .D.ESPVSP.
#EaMIC15 .FERLKVG-- ---------- -----TVL.F .-----HVKT T--------- ----A..AVS WML.SG.E.T .R.AQPVYPT
#HhMIC14 ---------- ---------- ---------- ---------- ---------- ---------- ---------- ----------
#HhMIC15 .FKARGKG-- ---------- ----T-TLD. .-----TYEL G-----PENR ET.VAELHT. WT..SSS.SR .L.GVEFA.P
#NcMIC14 ..TDRLEN-- ---------- ----I.KLSS .-----L.ES .-----.... ET.V.SL.A. .L.VT.Q..S .S..KE.-E.
#NcMIC15 .FTDRAKG-- ---------- ----T-VLDR .-----TYDL G-----TE.R EP.AAELHT. WT..TAS.SR .L.GVEFTEP
#SnMIC15 .FKRRR.A-- ---------- ----DFRLSQ .--------- ---------- ---------- ---------- -L.GTITVHD
#TgMIC14 ..TA.LN.-- ---------- ----I.KLRQ .-----LVES .-----.... ET.V.S..A. FI.VT.E..S .A..KE.-E.
#TgMIC15 .FKARGKG-- ---------- ----T-TLD. .-----TYEL G-----PENR AT.VAELHT. WT..SAS.SR .L.DVEFA.P

#BbMIC14 PVVITSIPAS TNAAGFVQIR NVKPTGFEVR LSNDICSVGF AYPYTIV--- -GWLASSEGN FVVSGSESYV RVGVAVAYGN
#BbMIC15 ....VGV.RT IDPYYKTV.. ..ST.S...K .HR.N.TLAK TQSRSAAV-- -S.M.LP..R YLT.NVRNP. KAVKLQLSTF
#CcMIC14 ...L.GLMEP LDTF.Q...T D.SA...SI. .AL.Y.RI.Y .AA.AKA--- -S...V..ER YA....GTPF ...TMDITID
#CcMIC15 ...FSGA.QA LAGVPKLA.S A.TSYS.K.Q AFGS..GSSP QAASAELKVV TSY..LPP.Q Y.AKH.PIRL L..SLSLTQL
#CsMIC14 .SYVFASVIG .DSFPL.ALS RQTK------ ---------- ---------- -S.FPGGPPA NLPA.----- ----------
#CsMIC15 ....IG..DT LTPFVRPV.. ..SASN..LK ..RQG.TTSD TQQVDTATV- -S.M.MA..R YST.NVRNPI ..MTVEVR.D
#EaMIC14 ...FIGLLDP LGTF.Q...T A.SE...YI. .AE.H.RI.Y .PA.AQ.--- -S.M.I..DS .TMPT.GTLI .AATVD.TVG
#EaMIC15 .I.FIGS.TH IAGIPRLG.S E.TSDS.R.Q AFGVV.G.RP TAASSDLKIV ASY..IPP.Q Y.TKHASIRL V..SLEITAA
#HhMIC14 ---------- ---------- ---------- ---------- ---------- ---------- ---------- ----------
#HhMIC15 ...FLG..RD .QPFYTPVV. L.TK.S...K .YRNN.GLDD TRS-SSSPV- -S.M.IP..A YLTEFV.NP. ..MKLPMTTR
#NcMIC14 A..V..L.ST ...PS.I..K D.TS...KI. MK..F....L .S...T.--- -.....GQ.V .....AKR.. ...TTI.SL.
#NcMIC15 ...FLGVARN .QPFYNPVV. S.TK.S...K .YR.N.GSAD TRS-.SAPV- -S.M.IP..V Y.TEYV.NPI ..VSLPLTTR
#SnMIC15 AALV------ ---------- ---------- ---------- ---------- -P...LT..R HMPRNA.NGI .ALKVELNPP
#TgMIC14 ...V..L..T ...VPVI.VK G.NS...KI. MK..F..... .S...S.--- -......Q.T .L...AK... ...TTL.SL.
#TgMIC15 ...FLG..KD .QPFYTPSV. L.TK.S...K .YRNN.GLDD TRS-SSAPV- -S.M.IP..A YLTDFV.NP. ..MKLPMTTR

#BbMIC14 HEGVIRYFVP --MQSDSPVV LLQAQSSSAD DEERYIAAPI VTSATNTEAN FRVASVGQIP ATDAIHVGYM IFD--EIPKS
#BbMIC15 APTSMSLTFR NLKSPEDI.A VA.I..VDPA TTATEGLGV. .SVVESDHVE ISLVANKSVD -ISSVT..LL VMG-KQDTEL
#CcMIC14 KDMD.LPLIQ --VH.SK.LL .T.I.NIAS. LPPTD.PFVT ISRLDA.L.T IS.HTTSTAA QQKKFT...L YM.--Q.EST
#CcMIC15 GAFQ..IPFA -IQHPSKA.I ..EP.AATGR FPAVIL..SA FNLLELNAQG LQARLSCTED -FAS.T...L .A.LS.AGSK
#CsMIC14 ---------- --------.A F.EKD.MRTG SAAAEQV.-- ---------- ---------- ---------- ----------
#CsMIC15 .PALVSTGLD NLKDP.EV.A ...V.KVEPP TTTPDLVDVV IS.LSSSRIQ LLPVMKSKVD GAHTLT..LI VMG-KQH.TL
#EaMIC14 E.IDL---L. GLKLRT.STK .MDPK----- ---------- ---------- ---------- ---------- ----------
#EaMIC15 GDFLL.VPFT -LANPSEA.. ..EP..AIGK SPNAIA..AV FRTIE.NTDG IKIRVSCVE. -FVS.S...L .A.MSYDSS.
#HhMIC14 ---------- ---------- ---------- ---------- ---------- ---------- ---------- ----------
#HhMIC15 TPT.LTFSLS GLKEPEEM.A .A.V.DVTPT SVSTDRV.VV ISNLKTNALE LSLVVDESVY -FSTVT..IL .SG-KQN.DL
#NcMIC14 QTTT...LPR --.N.AT.L. ...H.D-.GR TSGD..V.AN LMASSSS..T .QLTT..PGE PE.PLK.... V..--..DQD
#NcMIC15 SPT.LTVSLS GVKAPEEV.A .A.L.DV.PN TVSTGRI.VA ISNLKSNSLE LSLLMDDSVS -FSSVT..IL VSG-KQH.DL
#SnMIC15 KNLPV.VDFE NLRF.YEA.. ...V.QINPP NTSVEAVSVG ...LHSWG.E IALLEETPSR -FSSVV..LV FIG-KQS.EL
#TgMIC14 QPTT..HLPR --.N.AA.L. F..HED.EGG S-GN..V.ST ..ASSAS..T .KLTL..SGQ PSEP.VI... ...--..RQE
#TgMIC15 TPT.LTFSLS GLKEPEEM.A .A.V.DVTPT SVSTDRV.VV ISNLKTNSLE LSLVVDESVN -FSTVA..IL .SG-KQN.DL

#BbMIC14 KCSVGCKLNG ISL------- ---------- ---------- ---------- ---------- ---------- -QTIVGQRPQ
#BbMIC15 AAGLPPL..R FR.------- ---------- ---------- ---------- ---------- ---------- -H.FRVRK-G
#CcMIC14 E..T...T.N LA.------- ---------- ---------- ---------- ---------- ---------- -E.KSATLNK
#CcMIC15 SSPLAAS.G. R..AVSYLGE SGNGTKIKRG GKGAGSDCGQ EAIALTLRGN NANCNILCWA FCSFLLPRQA S.SLAPS.E.
#CsMIC14 ---------- ---------- ---------- ---------- ---------- ---------- ---------- ----------
#CsMIC15 AAGLPAV.SH RRF------- ---------- ---------- ---------- ---------- ---------- -..FSLPE-G
#EaMIC14 ---------- ---------- ---------- ---------- ---------- ---------- ---------- ---------S
#EaMIC15 SSPLAAS.G. R..AAN---- ---------- ---------- ---------- ---------- ---------- SSEFFV---.
#HhMIC14 ---------- ---------- ---------- ---------- ---------- ---------- ---------- ----------
#HhMIC15 AAGLSPL..R YH.------- ---------- ---------- ---------- ---------- ---------- -..FRIPA-A
#NcMIC14 Y..LS.Q... .A.------- ---------- ---------- ---------- ---------- ---------- -..VYENISG
#NcMIC15 AAGLYPV..R SY.------- ---------- ---------- ---------- ---------- ---------- -E.FRIPG-D
#SnMIC15 SFGIPPT.SR RRI------- ---------- ---------- ---------- ---------- ---------- -E.FFVDS-L
#TgMIC14 Y..L..Q... ..M------- ---------- ---------- ---------- ---------- ---------- -..DYADSNE
#TgMIC15 AAGLSPL..R YH.------- ---------- ---------- ---------- ---------- ---------- -..FRIPA-A

#BbMIC14 TWD--EERTD GVMIGGN--T TVFGTFIAPI TTSPTVVSRQ AVAWSAGERL TPSGGVVFDW DFPVGGSERT QASTGAHEGV
#BbMIC15 AYSASHIFGK D.PFA.PNMP YI.S.T.ITR S.DQVFQVSD Q.VITRRPTT D.FTWSALL. KAICEPTRHF K.IPDSADLE
#CcMIC14 KKKSPVSYSA ENFSQA---P LM..SIVPTK .D-------- ---------- ---------- -----.KK.M L.Q..SQD.S
#CcMIC15 LF.STQVDLA AIPFPE.-LP .F.TPHFF.Q VARVGFDGNE KYSIQWKASA DSATWTPSA. MALC.TD.LL SLMPHSMLTF
#CsMIC14 -----ARSG. AMSQN.E--- --------ST .A..KTR.S. QNTTGN.GTE NSRTSS..-- ---------- ----------
#CsMIC15 LF.PAAILG. DMLFS.TTAP HL.ASAVVQR .SDSVPDVGD E.TLAKRPTM ETMTFSLLA. ---------- ----------
#EaMIC14 G.SNTITYST E.F--..RPP LI..SVVVRV GGKTKMLVKD QLEIASPT-- ---------- ---------- ----------
#EaMIC15 IFNSR.VDL. DIHFPA.-LP SF.PPHFF.Q LAGERPD.EE VFSVQWQHKE DGR.WIPSV. ASSCSVT.DV LFKVPNSTIR
#HhMIC14 ---------- ---------- ---------- ---------- ---------- ---------- ---------- ----------
#HhMIC15 EYTATSIFGE .LYLS.GIMP H..ASA.QTR SRTEELEESD R.VI.RRATT ..FTWSALL. KKTCQAEHVF E.VENPSDLT
#NcMIC14 G.T--F.S.. S.VLA.H--. V.Y.AVVD.S HSK.VLI..S E.S.FP.SDI RSE....L.. GYV.P.GKD. .TPISSRAAQ
#NcMIC15 KYAATSILGE .LFFA.KYMP H..ASV.HTR SRDQQLEESD Q.VI.RRATT .AFTWSALL. NKTCQTKRLF E.VQD.TGLL
#SnMIC15 H.TLNSSQR. PIGERV.KHS H..A.PVL-R FDPKGMRPTD .IVLQQRPDM NGINWKLLP. KRICSSRGSY GVADARDFTK
#TgMIC14 R.T--A.S.. N.ILA.Q--. V.Y.SVVD.D G...VF..KH D....S.NEI RSE...IL.. RYA.P.GKDE H.RSHPSGAQ
#TgMIC15 AYSATSILGE .LYLS.GIMP H..AAA.QTR SMTKELNESD R.VI.RRATT ..FTWSALL. KKTCEAKHMF E.VENPSDLI

#BbMIC14 RSVPQAQNEG GDDFRGRTAA IQLRMNTKTR Q--VETPEAE DEDAAVFRVS DADGSAASRW FPVIDTVFDS NCEQRSFRRS
#BbMIC15 VV-------- .FYIEANS.E PTPSTPELGD I--CSLFS.Q SFVTTLKQNC VNLCN--DGK PSSVVEHCEQ K.RP-VVDPE
#CcMIC14 TK-------- ---------- ---------- ---------- ---------- -------AQ. T.KAFV.TND I.SEFV.QEV
#CcMIC15 T--------- .LYIDATLSS .CNVKAIEDK A--LTDA.CT EMCSG..QL- -------YCS NSSNPAYCIE LHSPKD.LD.
#CsMIC14 ---------- ---------- ---------- ---------- ---------- -------GN. Y....LIQ.G D.AEG.LQ.G
#CsMIC15 ---------- ---------- ---------- ---------- ---------- ---------- ---------- ----------
#EaMIC14 ---------- ---------- ---------- ---------- ---------- -------A.. S.KLYVISQE S..DFV.EEV
#EaMIC15 FT-------- .LYI------ ---------- ---------- ---------- -------.AI SSS.------ ----------
#HhMIC14 ---------- ---------- ---------- ---------- ---------- ---------- ---------- ----------
#HhMIC15 I.-------- .IYVEA.EDE PSAILGNRHD L--CLSFIGQ SFESG.VAAC MG.CR--.AL PASMQLHCQH E.ASNI.AP.
#NcMIC14 NA-------- .SQSSRSAEN SGGD.GGSAL GSLMQMSQK. SPAL.L.SNT R------.G. Y.A..IINN. R.TTG.VS.T
#NcMIC15 IA-------- .LYLES..PE PSAILANRQD I--CLAFIGQ NYENGLV.AC TDACR--NAL PSSL.QHCVH E.RSSI.APG
#SnMIC15 YV-------- .LLVEALES. PLVLTGKDDL ETPC.DVV.H QFVR.RSASC EEQC.EEDGD MESTMNQCVD T.VEG--DSY
#TgMIC14 T.-------- ---------- --DSKTS.ND IGESLMQVS. NLSVPGA.-- -------.G. Y.A..IIYN. K.TVG.VS.T
#TgMIC15 IA-------- .IYVEA.KDE PSAILGNRHD L--CLSFIGQ SFESG.VATC MGACR--.AL PASVQLHCQD E.ALNI.AP.

#BbMIC14 NELPAKQSS- -------VRL LYVRTGRSRL FVVGRRAPSK RPARRLQRAD AQDGLDLRDL VSVHGGGQSS PLGADSLRRQ
#BbMIC15 GLEAC..G-- -------CAE NIEDEFQK-- ---------- ---------- ---------- ---------- ----------
#CcMIC14 PSGSSTLR-- -------MVA .H.QSHITPN ---------- ---------- ---------- ---------- ----------
#CcMIC15 CT..RRLQQT -------TSP RQPQRHS.AF LCFAWPL--- ---------- ---------- ---------- ----------
#CsMIC14 TTF.SSPAD- -------... ..AKSVSATP PSQEN.PVYR PD-------- ---------- ---------- ----------
#CsMIC15 ---------- ---------- ---------- ---------- ---------- ---------- ---------- ----------
#EaMIC14 SV.F.TVL-- -------LMV .HIKDHFTPN ---------- ---------- ---------- ---------- ----------
#EaMIC15 ---------- -------CNV KA.QDL.N-- ---------- ---------- ---------- ---------- ----------
#HhMIC14 ---------- ---------- ---------- ---------- ---------- ---------- ---------- ----------
#HhMIC15 VHATCMSD-- -------CES .LEAQYEE-- ---------- ---------- ---------- ---------- ----------
#NcMIC14 .RF.NGSAQN PSKK---.A. ...SS.T.E. SSSNLPSYGV D--------- ---------- ---------- ----------
#NcMIC15 VNEACMTD-- -------CAR KVEAQFED-- ---------- ---------- ---------- ---------- ----------
#SnMIC15 ..GACRGV-- -------CEA KLP..S.E-- ---------- ---------- ---------- ---------- ----------
#TgMIC14 .IF..AGGRK NSVSRRL.A. ...SS.T.EI SSSSFPPYGA D--------- ---------- ---------- ----------
#TgMIC15 VHTKCMTD-- -------CES .L.AQYEE-- ---------- ---------- ---------- ---------- ----------

#BbMIC14 ATRGIRESHA GLPLRARTRR AQRRRAWLRR RVCLCAQERA GQLPRTCPAS SHTDAKCSEE CTAVTEACSV T--SENMWVC
#BbMIC15 ---------- ---------- ---------- --.ASQCQD. D--------- ---SLA.ITS .ED.MANQCR SAS.KDAMN.
#CcMIC14 ---------- ---------- ---------- --.TE.KPKN T--------- AAH.VD.VK. .SSLSAL.PP P----.S.D.
#CcMIC15 ---------- ---------- ---------- --SEEDMPSD EAQGE.SGKR EERLLVFFF. AGRLLLLRGG P--RASSSMR
#CsMIC14 ---------- ---------- ---------- --.QA.RQKE .--------- --SSRD.VT. .EELAR...A K--GRDI.E.
#CsMIC15 ---------- ---------- ---------- ---------- ---------- ---------- ---------- ----------
#EaMIC14 ---------- ---------- ---------- --.AK.KSKG .--------- .A...D.VA. ...IS.K..P P----LA.E.
#EaMIC15 ---------- ---------- ---------- ---------- ---------- --S.GIMIDH .R..PCDAAE N---------
#HhMIC14 ---------- ---------- ---------- ---------- ---------- ---------- ---------- ----------
#HhMIC15 ---------- ---------- ---------- --.AGQCTAT .--------- ---T.H.HN. .KQFLGNKCD NTA.SDVAA.
#NcMIC14 ---------- ---------- ---------- --.AS.SGK. .--------- -SATTA.IKA .QTTE...TE A--D.KA.P.
#NcMIC15 ---------- ---------- ---------- --.AGQCSIE S--------- ---TES.HRK .KEFLGNKCE GVSLADVEA.
#SnMIC15 ---------- ---------- ---------- --..RQCRA. E--------- ---.KR.K.I .EDLVASLCE R--HPDKRL.
#TgMIC14 ---------- ---------- ---------- --.AS.AGKK N--------- -YAA.E.AAQ .E.TV.G.TD S--D.DA.P.
#TgMIC15 ---------- ---------- ---------- --.AGQCTAT .--------- ---T.S.QRK .KQFLGNKCD STA.SDVAA.

#BbMIC14 FLRRFPHALK DAC------- ---------- ------RVLP ETAACILVNM ESKEYSAMRG WDPVTMTCKC PYDVPACSKE
#BbMIC15 V.YGL.VTEL E..VFNEAYT SDQTPIPELP EAEEDMP-NQ SY.D.FVLD. R-QG.DRFI. ..SR.A.... .N.FA..TSA
#CcMIC14 .VAAASEK.. AD.-----YI ESYIDTGQTV TTSTTVPPIG D.VQ.RV... .DD..F.DA. ..VNA.S.L. .QGFSP.TQQ
#CcMIC15 ..SF.AWHRA H..---MRLF TASVRHSGAA AFAAAMEDAE TS.Q.RV.D. R--K.PSDLA .NSAAF..G. .SHAVP.TSA
#CsMIC14 .VEST.KEW. QK.------T VMPTSATEAL TTSTTAPPID PA...TI.D. QAL..QN... ..SK.DS.N. .HEI....PQ
#CsMIC15 ---------- ---------- ---------- ---------- ---------- ---------- ---------- ----------
#EaMIC14 .IASASQE.Q AK.------- ---------- ------YIE. ASVQ.RV.D. S.R---LDI. ..VEA.S.S. .QGFSP.THH
#EaMIC15 ---.P.DDIS EEA------- ---------- DVGEAVEEGL PLVK.RVFD. R--THPSNLA ..SASF..G. .GLAVP.T..
#HhMIC14 ---------- ---------- ---------- ---------- ---------. SEAA.A.E.. ...T...... .NEA....A.
#HhMIC15 LEFVT.ISVF EQ.TLLVEYT VGEGGASEGF EVEEDTTPSQ TYEN..I.D. RDESFA.FA. ..SRVAS... .NGFR..TS.
#NcMIC14 .F.Q..LEQ. RS.------Q ITAVASDAVA TSTSTPPPVD HAER.V..D. REEA...D.. ...N...... .NGA....A.
#NcMIC15 LEYST.V.FF EE.ILVITYS SKEGTVSEGF ESETETVPDQ TYDD.VV.D. QDPRFATSV. ..SRVAS... .NGYR..TS.
#SnMIC15 LQ..------ ---------- -----DQDSF DTAQENEDDS SASQ.TI.D. S.P.LKSFA. ..VD.R.... .QGTH...SS
#TgMIC14 .F.Q..IDQ. KK.------Y ITAVASDLAA TSTTTAPPVD KDER.V..D. SEAA...E.. ...E...... .N.A....A.
#TgMIC15 LEFVT.PSVF EQ.TLLVEYS VSEGGPSDGF EIEEETTPSQ TYEN..I.D. RDERFT.FT. ..SRVAS... .N.FR..TS.

#BbMIC14 QATFDLFNWS SSVGHRGSLC TGQ-TKVV-K GLARGFKLTS VDLCG--SL- --PDFSSSTW QNLPLPPYMA DDVTVGDTVL
#BbMIC15 AVDAGHHWRT ELLRP-DG.. AA.GDAAF-M A..QR.RS.T GN..E--LA- --KIG.RI.. EDNT..A.PD Q..NTE.S.V
#CcMIC14 .VEQ.RSY.I ED.TPWSDI. RL--SYT.-Q PV.W..LQVA N...------ --MNS.NLQ. GQRQ..L.SL N..YLDRGY-
#CcMIC15 E.GR..AWLR DLRSK-.G.. EKAPAE..-Q PVVQEYWQL. ..A.SLE.-- --SGNVYLS. IKKTY.A.PD LPGPGTGQES
#CsMIC14 ..L..SLT.. KLL.PE.... KSA-GGTL-. VV.H.....Q QN..E--KG- --FST.QTS. EQI.F...AG E.LIL..NTI
#CsMIC15 ---------- ---------- ---------Q KVCQS.R--- ---------- ---------- ---.YS.ADT K.IA.VGLY.
#EaMIC14 .VEQ.RAH.I QDISPW.Q.- ---------- ---------- ---------- ---------- ---------- ----------
#EaMIC15 D.SQ..TWVQ EIHAE-.G.. HSA-EEA.-Q PVTKM.LQLD Q.A.SANM.- ---QSHTL.. SQRTY.A.PD LPGAGSGPQS
#HhMIC14 .T.YN.I.H. .VALGQ.AI. AAA-.GT.-S .I..H..RST .....--D.- --SNL.P.S. EQ........ ...VLD.E.I
#HhMIC15 TVNTQNHWRT ALLAS-AG.. KE.PNGMF-N AVTQQLWS.T G...ALAAD- --EQQPPIS. ASNM..A.AD Q.IVLDNSTI
#NcMIC14 ...YN.V.H. .V.LNQ.TV. ASAAEA.F-- .I..Q..R.T A....--D.- --TNV.T.S. EQ..F..... ...N.D.E..
#NcMIC15 TVNAQRHWRT NLRLP-.G.. EA.PNGAF-N PVTQHLWS.T D...ALAAD- --ATEPAI.. SSNR..A.AD Q.INL.TSIA
#SnMIC15 AVEQQIL.RA ALREAVQTF. EERQSPTFAT PVT...-A.P EN..EAVQ.G NRKQSGPL.. KQI.P.V.SD L.LKLD.VAV
#TgMIC14 ...YN.I.H. .VALGQ.AI. AAA-.GT.-S .I..H..RST L....--D.- --SKL.PLS. EQ........ ...ILD.E.I
#TgMIC15 TVNTQNHWRT ELLAS-AG.. KE.PDGRF-N AVTQQLWS.T G...ALAAD- --EQQPPIS. ASNM..A.VD Q.IVLDNSTI

#BbMIC14 RSGHHLCTQE YDYVFCP--- ---------- ---------- ---------- ---------- ----SKLITT TT-TASTSKW
#BbMIC15 ...EYE.RNS WF.....--- ---------- ---------- ---------- ---------- ----ADATS. ..-V.P.VTK
#CcMIC14 ...YNQ..TD VL.....--- ---------- ---------- ---------- ---------- ----AASL.. ..-STAAPV.
#CcMIC15 L..S...QYS TPF....GER MPWGRRGGSL LLIDSRGVEK EASAAFFLQQ SHCPFAFRTA AQATGEELDE EKKEEASGSD
#CsMIC14 .G.YYR..E. FR.....--- ---------- ---------- ---------- ---------- ----G..LY. ..-.TAEPP.
#CsMIC15 TA.EAEFAAV SEF.------ ---------- ---------- ---------- ---------- ---------- ----------
#EaMIC14 ---------- ---------- ---------- ---------- ---------- ---------- ---------- ----------
#EaMIC15 V..TY..QHT TP..L..--- ---------- ---------- ---------- ---------- ----ADAAGV ADEEEE..RI
#HhMIC14 ...YY...KS .HH....--- ---------- ---------- ---------- ---------- ----.S.... S.-.TPAP..
#HhMIC15 ...DYQ.SEI WH.....--- ---------- ---------- ---------- ---------- ----AAATS. ..-QEPPAPP
#NcMIC14 ...YY..SKT .N.....--- ---------- ---------- ---------- ---------- ----.S.... ..-.TPAP..
#NcMIC15 .G.EYQ.SKK WH.I...--- ---------- ---------- ---------- ---------- ----AQATS. ..-Q.PPPAD
#SnMIC15 P..DS..RSR WTH...---- ---------- ---------- ---------- ---------- ----TIQTSK E.-A.APPSP
#TgMIC14 ....Y...KS .HH....--- ---------- ---------- ---------- ---------- ----.S.... S.-.TPEP..
#TgMIC15 ...DYQ.REI WH..V..--- ---------- ---------- ---------- ---------- ----AAATS. ..-QEPPVPP

#BbMIC14 DWPANQDCVA GEWTSWSSCS EHCYSNT-GP RAIKRRTREV FLPRRGSGED C-TLLQHRLC EVDEEVPLCS KYCWRSEWGD
#BbMIC15 EEITLKTAIV ...GE..P.. GT.I.S--SW TPKRM.K.Q. LAQLPT.ATP D--.WEMED. ---LGL.I.G TA..D.D.TE
#CcMIC14 ...E.A..LP .Y.AD..E-- ---------- ---------- -----.N.TP .-..EETVS. AS.TLL.-.D SL.LH...TE
#CcMIC15 AALDPPS.YE S..GD..A.. AT.VAGD-.S TPYRH...IP LVADGLNTAV SCI.EESMQ. L--GLLQQ.E GS..TAD.SQ
#CsMIC14 S...DS...P TD..E..A.. AT.F.GSK.S KPA.T.K.FI LV.KK.R.Q. .-V..EQEP. .RSSDF.A.E EF..QGN.SE
#CsMIC15 ---------- ---------- ---------- ---------- ---------- ---------- --GLVL.--- ----------
#EaMIC14 ---------- ---------- ---------- ---------- ---------- ---------- ---------- ----------
#EaMIC15 IKHLPSS.FV ...GE..A.D AT.RGP--FH SPQRY.K.SP IVVEAVAEDP DCI.EEKQS. --GHSLEP.V G...TA..TE
#HhMIC14 ....R....P .......K.. MD..AGS-.A LPA.S.S.S. LID...T.NM .-V.ED.T.. K.GS...Y.E DL..ITD...
#HhMIC15 FTATLNTAIV ...GE..A.T GT.F.Q--WW TPKRT...V. LAELSN.QIP S--VSDTAT. ---LDL.A.G TV..ER..TE
#NcMIC14 ....DR...P .......K.. MD..PGS-.T LPT.S...P. L.A...T.AP .-A.EEQDI. NIGS...Y.E DL..IT....
#NcMIC15 FTTTLETAIV ...GE..A.T GT.R.Q--.W TPKRT...V. LTELPE.KIP S--.SETGA. ---FDL.E.N TV..DR..TE
#SnMIC15 SS.VLDVPIL S..LE.GG.. AT.RTD--QA VPVRT.R.FQ YV---.GSN. FPP..ETKA. ---DDL.V.G EV..M...S.
#TgMIC14 ....R....P .......K.. MD..VGS-.A LPV.S.S.S. LVD...T.NV .-V.ED.T.. K.GS...Y.E DL..ITD...
#TgMIC15 FTATLNTAIV ...GE..A.T GT.F.Q--WW TPKRT...L. LAELSH.QIP S--VSETAT. ---LDL.P.G TV..ER..TE

#BbMIC14 WSDCEEKVLK FGKPPVLTRT KQRHVFMDHN GVCSAEELVA YDQSTC--RG SRSGPAAPSP TPVPSLAEVG HSVARGFGEG
#BbMIC15 .G..A.YTVD LKAGAE.Y.R QVK.I.SYSA AA.GNDKF.R .EKCEE--EM AEVYEDVSAM DTTT.RGAGH .RGVYDGPD.
#CcMIC14 ..S.S..L.E ..VA..K.SL REKL.SAEAG EA.A.QH.KE ..A.R.TGK? G.RSNHQL.L PQPSA.ELGS SEIVDEG---
#CcMIC15 ..A.AL--VL KDREVIKA.R RYKP..AG-A KY.KID.IQQ FLPCESNEIQ EKASE.GSPQ FAEVFSLSES ----------
#CsMIC14 .GA.KLST.. ..ED.LR... .I.YI...GG D..PT.S.IQ H.TA..ANT. ..TV.SVEAT PSA..EIDEP QPLPSPKP.H
#CsMIC15 .QQPAAFL.V L.EQ.----- ---------- ---------- ---------- ---------- ---------- ----------
#EaMIC14 ---------- ---------- ---------- ---------- ---------- ---------- ---------- ----------
#EaMIC15 ..S.SA--VL S.GEISFS.R RYKP.LAGV- DY.DM.AIQE LQSCLRPVTT .SDASRL--- ---------- ----------
#HhMIC14 ..G..Q...Q I.QAA.RA.. .L..I..GTA ....SD-HIQ ..Y.G.--KS GMDSTGGRED GSAH.MV.QR SR.TDAV---
#HhMIC15 ..E.KLFTIV M.QGLEYF.Q QIKPI.DFVE EA.GQD.HER .ERCGEEQDN -EET..TSTL QTDSL.ETRS LT.SH.TAAH
#NcMIC14 ..G..Q...Q I.HS..RA.. .L..I..GTA ...GS.TH.K ..Y.G.--TS GMDSSGGR.D NAA..MV.QQ SRATDVV---
#NcMIC15 ....KLFTTV M.QGLEYY.Q QI.PI.DFVE DA.G...R.R FAKCGDANDS VEQRDSRITR RTDSPW.ASY LA.LSQSEGR
#SnMIC15 .GE.A.TFV- -.NAMAFVQA RVKY.YDY-. EA.ADDNFLE LRDCVDEYEI ---------- ---------- ----------
#TgMIC14 ..G..Q...Q I.QAA.RA.. .L..I..GTG ....SD-HIQ ..Y.G.--KS GMDSSGGRED DSAH.MV.QK SR.TDAG---
#TgMIC15 ..E.KLFTIV M.QGLEYF.Q QIKP..DFVE EA.GLD.HER .ERCGEEQDN -GET..TSTL QTDSL.ETRS LT.SH.TAAR

#BbMIC14 GEAQRSTAEE ETPGFSSGEE GERDPKSGGA APEHEDVEKE E--------- ---------- ---------- ----------
#BbMIC15 RLP-FPASPT RASSSLIPSA EQLSFFVDS. D.YAKLTQAR MGFMDRRRSR SRSVSVEDGS GAAVAQPEGV KHREPGANRR
#CcMIC14 ---------- ---------- ---------- ---------- ---------- ---------- ---------- ----------
#CcMIC15 ---------- ---------- ---------- ---------- ---------- ---------- ---------- ----------
#CsMIC14 R.TEEKGLAQ PNEKSAAPGN ..GK------ ---------- ---------- ---------- ---------- ----------
#CsMIC15 ---------- ---------- ---------- ---------- ---------- ---------- ---------- ----------
#EaMIC14 ---------- ---------- ---------- ---------- ---------- ---------- ---------- ----------
#EaMIC15 ---------- ---------- ---------- ---------- ---------- ---------- ---------- ----------
#HhMIC14 ---------- ---------- ---------- ---------- ---------- ---------- ---------- ----------
#HhMIC15 ALPRV.MVPA RERSSFASLH .T.S-HVPAS .D-AKII.RR KGFISRRRSV PPSGYLEETS E-QVS----- ----------
#NcMIC14 ---------- ---------- ---------- ---------- ---------- ---------- ---------- ----------
#NcMIC15 SASR..R.LT QASSPFYAAY ET.S.S.PVS TQ-V.RTQ.R RGFMQWQRSV TRSGGLEEDS DAQRG----- ----------
#SnMIC15 ---------- ---------- ---------- ---------- ---------- ---------- ---------- ----------
#TgMIC14 ---------- ---------- ---------- ---------- ---------- ---------- ---------- ----------
#TgMIC15 ALPRV.MDPA LERPSFASLH .T.S-HVPAS TD-AKII.RR KGIMSRRRSV PPSGYLEETA E-QVA----- ----------

#BbMIC14 ---------- ---------- ---------- ---------- ---------- ---------- ---------- ----------
#BbMIC15 HSEEPPDGHN EPGRRRGETE TNVARPAVGR GGRSQEAPGR VAYQVDSGNN EGLLAAIETV ASPDKLHGGG LGAGAASESE
#CcMIC14 ---------- ---------- ---------- ---------- ---------- ---------- ---------- ----------
#CcMIC15 ---------- ---------- ---------- ---------- ---------- ---------- ---------- ----------
#CsMIC14 ---------- ---------- ---------- ---------- ---------- ---------- ---------- ----------
#CsMIC15 ---------- ---------- ---------- ---------- ---------- ---------- ---------- ----------
#EaMIC14 ---------- ---------- ---------- ---------- ---------- ---------- ---------- ----------
#EaMIC15 ---------- ---------- ---------- ---------- ---------- ---------- ---------- ----------
#HhMIC14 ---------- ---------- ---------- ---------- ---------- ---------- ---------- ----------
#HhMIC15 ---------- ----HGGESE Q--SGKASNN GSRRHRTSRK QKRGLESIHS D---A----- ----SVHGSG ----------
#NcMIC14 ---------- ---------- ---------- ---------- ---------- ---------- ---------- ----------
#NcMIC15 ---------- ----RRDVDE Q--GGAAIQR ETRRPQATGK QRSGVDIAYT D---ADASTS ARAGDIYGSA ----------
#SnMIC15 ---------- ---------- ---------- ---------- ---------- ---------- ---------- ----------
#TgMIC14 ---------- ---------- ---------- ---------- ---------- ---------- ---------- ----------
#TgMIC15 ---------- ----HGGESE Q--SGKASQN GSRRHRASRK QKRDLESIYS D---A----- ----SVRGSG ----------

#BbMIC14 ---------- ---------- ---------- ---------- ---------- ---------- ---------- --------SS
#BbMIC15 KEVDTARSHE ELERAAPLVS DERQEDKGES SRNAETEIAS HAAAAPWDAG HGNVAMGTEL PKGMTEPAPA GEPVEPKKAQ
#CcMIC14 ---------- ---------- ---------- ---------- ---------- ---------- ---------- ----------
#CcMIC15 ---------- ---------- ---------- ---------- ---------- ---------- ---------- ----------
#CsMIC14 ---------- ---------- ---------- ---------- ---------- ---------- ---------- ----------
#CsMIC15 ---------- ---------- ---------- ---------- ---------- ---------- ---------- ----------
#EaMIC14 ---------- ---------- ---------- ---------- ---------- ---------- ---------- ----------
#EaMIC15 ---------- ---------- ---------- ---------- ---------- ---------- ---------- ----------
#HhMIC14 ---------- ---------- ---------- ---------- ---------- ---------- ---------- ----------
#HhMIC15 ---------- ---------- ---------- ------ESTL HGTGAN---A HREQKEWT-- ---------- --------.K
#NcMIC14 ---------- ---------- ---------- ---------- ---------- ---------- ---------- ----------
#NcMIC15 ---------- ---------- ---------- SNRSNVEATF NGPETP---A HGWAQIGS-- ---------- --------A.
#SnMIC15 ---------- ---------- ---------- ---------- ---------- ---------- ---------- ----------
#TgMIC14 ---------- ---------- ---------- ---------- ---------- ---------- ---------- ----------
#TgMIC15 ---------- ---------- ---------- ------ESTL HGTGTN---A YRDQIEWT-- ---------- --------.K

#BbMIC14 RSPPSEGAEV ADDGATFSLR SANAGTVNA- ---------- ---------- --------DD EDQTHASPRS HVHNRERALR
#BbMIC15 A.GGPLT.PA ...TVATGSE T.AFVGRASR GRKRARTDNS STRTRRGTRA ARAGVVPVCS AVSAWSGCD. PCAIH-QGTA
#CcMIC14 ---------- ---------- ---------- ---------- ---------- --------GE SWSEWS.CDA PCLMN--GRK
#CcMIC15 ---------- ---------- ---------- ---------- ---------- --------TV .TSDWSGCDV PAAYK-----
#CsMIC14 ---EN.SSDR P.Q.G.A.GS LLQHSSETVE S--------- ---------- --------C. RLSAWSGCTA PCQPPAGD.G
#CsMIC15 ---------- ---------- ---------- ---------- ---------- ---------T ATVA.SGS-- ----------
#EaMIC14 ---------- ---------- ---------- ---------- ---------- ---------- ---------- ----------
#EaMIC15 ---------- ---------- ---------- ---------- ---------- --------CS TADGMDECKN LFIDT-----
#HhMIC14 ---------- ---------- ---------- ---------- ---------- --------CK KSSGWSACSL PCQQPGAKAV
#HhMIC15 S.SG.ATRDS GAR..EADTS LLQKARRARR GAGRFRNTRS RARNQTP--A --------CS VVSEWSGCD. PCLPH-KGST
#NcMIC14 ---------- ---------- ---------- ---------- ---------- --------CK KSSRWSACSL PCQQPGAKTV
#NcMIC15 ..SEAAAGTS Q.RA.GAGAD LL----RTQL STGVTVDAGN LSGVQDHDRA --------CS TVSEWSGCD. PCLPH-KGAT
#SnMIC15 ---------- ---------- ---------- ---------- ---------- --------YE TIGAYDEGDL ----------
#TgMIC14 ---------- ---------- ---------- ---------- ---------- --------CK KSSGWSACSL PCQQPGANAV
#TgMIC15 S.SR.AIRDS GAR..EAGTS LLQKARRARR GAGRFRKSRS QARNQTPDKS --------CS VVSEWSGCD. PCLPH-KGST

#BbMIC14 EDSFNSGSQK LYLRTKTTAL IMYAAAKDRC RLASSTSNRV -NCNLVGPRY SSPEDLLYCR QQCEYIVKEC KKTAA---LR
#BbMIC15 ARRYRLAL-P GLNNARYCT. PALDDKHLCP D.QACEYPKI -D.SM.TAGR QTE..ATE.Q IV.KATFAS. .TMMTGTVA.
#CcMIC14 PRRHRLSIKP GANEEGGRVY Y--------- ---------- -S.DDIF.QH DI.GTRE..Q EK.RE.L.S. SDESS---.K
#CcMIC15 --GYVLSL-- ---------- ---------- ----CA.--- -D.SQAT.VH VDSQSGDA.V KK..AVKQA. T.LLEKN-AL
#CsMIC14 VKQYQL.L-P RNVD.PIS.Y CATNTRDCSE K.QACKEDPT PD.S..SG.. AAADETKQ.. EL..IA.DA. A.S..---.I
#CsMIC15 ---------- ---------- ---------- ---------- -P..A.RAHS LI.------- ---------- ----------
#EaMIC14 ---------- ---------- ---------- ---------- ---------- ---------- ---------- ----------
#EaMIC15 ---------- ---------- ---------- ---------- -Q.SD.VAT. RE.AS.RN.V DK.KTVEAK. RTLSSQD-AL
#HhMIC14 AEQ.QL.L-P ETVAAS..SY CASRSRACME KKPPCVLDVA PD.S..K... D..QEA.H.Q EL.ANALTR. .EQ..---.Q
#HhMIC15 PRRYRLAV-P GQNEANYCTV PLSGKEHLCT D.PQCAHPNF -D.AK.TASR LTD..IAA.K TV.AEV..A. EAIMSAVFSL
#NcMIC14 AQQ.QL.L-P DTVTDS..SY CSAR.RDCTD SKPACVLDIE PD.S..K... D..GEAAE.Q EL.ASAKAR. RDQ..---.Q
#NcMIC15 PRRYQLAL-P GQNQAHYCTV QHVGHEHLCT N.PVCAHPTF -D.SK.TASR MTD..MAA.K TV..EVLAV. DSMMNVPFAL
#SnMIC15 ---------- ---------- -------LSD DVDPLVAEDA GQSEK.R.SR GAHSEAAASL YADFTQQSQH TSLGTASL.F
#TgMIC14 TQQ.QL.L-P ETVTAS..SY CASRSRACVE KKPPCVLDVA PD.S..K... D...EA...H EL.NNALTR. REQ..---.Q
#TgMIC15 PRRYRLAV-P GQNEANYCTV PLSGTEHLCT D.PQCAHPNF -D.AK.TASR LTD..IAA.K AV.VEV..T. ETMMSAVFSL

#BbMIC14 NMTQWECV-- --IEGLGGIG RCRYKKDGAA AQAMRPPV-C FLSRTRQV-D KSNPLSFLAT GKEYGRG--- ----------
#BbMIC15 YTSIEH..LV RFA.HQSYP. Q.QLTTEVQG GASHPQDTK. .P.S..RR-V A.GEFP.IFQ LPVSK.SAVS SDEPQNNSAD
#CcMIC14 EQ.PLQ.FHQ NIVQR.TLT. Q.H.NP.L-- DKE.LA..-. .P.L..MND. D.SEFR..DP NSPSA----- ----------
#CcMIC15 ESPFKS..LS ELASSEFYL- T.KFSGALQT GKVA-----. RPTF..II-. P.SRYA..KN TAPDEETSAD TGSVEDSRLP
#CsMIC14 QQS.....AA TLNDAGD--. S.L.EREAVE DRVV....-. ....AQRA-. ENTS.P..H. DEAT.AK--- ----------
#CsMIC15 ---------- ---------- ---------- --STS.SLYL T.KK..LIRA SRTA..LTWD PALSS.R--- ----------
#EaMIC14 ---------- ---------- ---------- ---------- ---------- ---------- ---------- ----------
#EaMIC15 KTPFKR.FLN GLSDAGFFL- T.KFAEK--- -.KISR.AT. R.TF..V.-N PDAEF...TD SAAE.APAES PGGQSFIAP-
#HhMIC14 FQ.....I-M AF.NSS.VT. ..QIQ.GAVS KTTL....-. ........-. A.KS....T. D.TS...--- ----------
#HhMIC15 YTSLEQ.ALV QFQ.YEQFP. Q.HIPEEY-- -TSTQRATK. .P....RR-V A.GE.P.IFQ V--STASGTS P---SDASSS
#NcMIC14 HQS....I-M AA.NSA.LT. ..HFP..TIS QNTL....-. ........-. ATKR..Y... D.DS...--- ----------
#NcMIC15 YTSLEQ.ALV HFD.YEQYP. Q.HIPSEF-- -TST.Q.ST. .P....RR-V A.GE.P.IFQ LSASTSSKVS PSETLDSSPS
#SnMIC15 TGSNLSVLDL TAKGDGDSA. NISNPDT.-- ---------- --.DVQKQ-. IGS.VPTSNG .ESPVA.--- ----------
#TgMIC14 FQ.....I-M AF.NSS.VT. ..QIQ.GAVS KTTL....-. ........-. A.KS....T. D.TS...--- ----------
#TgMIC15 YTSLEQ.ALV QFQ.YEQFP. Q.HIPEEY-- -TST.R.TK. .P....RR-V A.GE.P.IFQ V--STASGTS PSATSDAASS

#BbMIC14 ---------- ---------- ---------- ------SCMC ASPDNVPCSA EEAAAS-ADE WTKLLNTSVC PFGTLG--AF
#BbMIC15 ATPSLVSMGT FAAPTSGVVN PLSPSWSASS TTAFID..E. FD..HE..T. P..RDAAYNF LYPA.TYPL. AAT.S.EK..
#CcMIC14 ---------- ---------- ---------- ------Q.L. LEK.A...T. A.VFE.-REY AFPSSDI.S. .LQDSA-G.M
#CcMIC15 V--------- ---------- ---------- ------..A. LEDGLE.... DDIVQDWLNI RNA..GA.GL CTTA.AQEPL
#CsMIC14 ---------- ---------- ---------- ------..K. .T.AT..... ..VWR.-S.T .G..ISST.. .LND..--..
#CsMIC15 ---------- ---------- ---------- ---------- --.AGT.V.G PAC.QMDFEP VRPI.L.VK. T---------
#EaMIC14 ---------- ---------- ---------- ---------- ---------- ---------- ---------- ----------
#EaMIC15 ---------- ---------- ---------- ------..A. LDEGLE..T. .DILEDWKGI KDSF.AADGL CSAEDALKPL
#HhMIC14 ---------- ---------- ---------- ------.... .VTNS....P ..V.L.-F.D .A.Q.GST.. ...Q..--..
#HhMIC15 VAASFVSAGT VEAPRSRVVN AL--AFEASA SQTSID..E. LD.ADE..T. Q..RD.LF.S LYLF.THPL. AESPAAED.L
#NcMIC14 ---------- ---------- ---------- ------.... .ATNS....P ..V.L.-F.D .S.Q.TSD.. ......--..
#NcMIC15 FSASFVSMGT LAAPSSRIRN AL--DFEAAA SGRYID..E. L..EDE..T. Q..RD.LF.S LYLF.THPL. ADSPGAEE..
#SnMIC15 ---------- ---------- ---------- ---------S .EEHQETTL. T.R.DG---- -SSNVRHDGG VHESD.----
#TgMIC14 ---------- ---------- ---------- ------.... .VTNS....P ..V.L.-F.D .A.Q.GST.. ...Q..--..
#TgMIC15 VAESFVSTGT ASAPSSRVMN AL--AFEASA SQTSID..E. YD.ADE..T. Q..RD.LF.S LYLF.THPL. AESPAAEG.L

#BbMIC14 FSSPDSKGS- -ADSAVFFGA TDLGRVHCPI --SQGTEPGG DQP-----TY TDFASPEAMN EFCEHGLPFW ----------
#BbMIC15 .TRDELT.R- -..ASTY.AL KGMQ.I.... PMYRALIE.K .SKVVNVV.F SK.E.K.EL. N...K..EV. ----------
#CcMIC14 .G-.WGS..- -...KLY.A. A.SSKIF..L SN.KNQ.QKS .NA-----.. SK..T.AE.. DY.AN..D-- ----------
#CcMIC15 WQRSEESATK AP..SI.LAF AG...F...L ---HTSIKDT TADSQERF.F .Q...T.SL. D..Q...NY. EGVENPLTCK
#CsMIC14 .DKANLS..- -...S...A. SGGV.....V ---MTSPS.A SL.-----.. .S.KNDAELD T...K..SS. ----------
#CsMIC15 ---------- ---------- ---------- ---------- ---------- ---------- ---------- ----------
#EaMIC14 ---------- ---------- ---------- ---------- ---------- ---------- ---------- ----------
#EaMIC15 .--------- ---------L LKSNAL.HR. TTAKS.D--- ---------- ---------- ---------- ----------
#HhMIC14 .N..N.E..- -...F.Y... S......... --.KSSDL.S RF.-----.F .E.E...S.. N...N..... ----------
#HhMIC15 .TMSEPNQT- -..A.SY.AL RG...M...V PMYKSLTEVK SVLTPKRV.F SE.KTK.EL. D..HK..SN. ----------
#NcMIC14 .N....TA.- -...F.Y... A......... --.KSSDL.. HL.-----.F AQ.E...S.. S...N..... ----------
#NcMIC15 .N.THPNQT- -..ATSY.AV RG...M...V PMYKSSVQVK SVLTPKRV.F SE.KTKQEL. ...HK..ST. ----------
#SnMIC15 -KL..A.PE- --------RQ D.TSAAER.V DS.NVEQE.V HGSDG---KL .EARAA.ERQ G--------- ----------
#TgMIC14 .N..N....- -...F.Y... S......... --.KSSDL.S RF.-----.F .E.E...S.. N...N..... ----------
#TgMIC15 .TMSEPNQT- -..A.SY.AL RG...M...V PMYKSSTEVK GVLTPKRV.F SE.KTK.DL. ...HK..SN. ----------

#BbMIC14 ---------- ---------K NTALSVGSLD CRQAKSKA-- GKQK------ DCRAACQKLL NACIASDE-- -AWENCVAKS
#BbMIC15 ---------- ----DRRASN SIRDGPIIPR .EA.S.VT-- .DDNLNG--- V.QRR.IQSR TY.S.RS.TV TELAS..KDQ
#CcMIC14 ---------- ----AQYQYS SSTPYDID.. .DAVVPRDAT ASLE------ ..QSK.RQIQ ED.EI.TV-- -NFLS...TK
#CcMIC15 PAAFRDGCLH AAAPDASAAA E.RADTRMP. .SG.MA.NSE .PEPTEGTV- F.QDL.R... R....EAQEF -SY.A.LIRP
#CsMIC14 ---------- ---------- -SGTTESIV. .SH.ML.DTA .TK------- ......E.V. AK.DK.AP-- -DLKA.YKEA
#CsMIC15 ---------- ---------- ---------- ---------- .NSR------ ---------- ---------- -S.RRT.YAE
#EaMIC14 ---------- ---------- ---------- ---------- ---------- ---------- ---------- ----------
#EaMIC15 ---------- ---------- -----ARIP. .SK.IA.TPH .VEPAEADTA A.KKL.LE.K DT.KDLLKKE -SYDV.LIEP
#HhMIC14 ---------- ---------- -QETPEPLV. ..EVVP.N-- ASTT------ ..PGR.H.AV VN.M.NSD-- -SLHS.INEA
#HhMIC15 ---------- ----KRNVPP ELIG.ASIPN .MLVTPRE-- .AEP-Q---- ..TLL.SETI SS.S.ASLSF VELSQ.IEDK
#NcMIC14 ---------- ---------- -KETLQPVV. ...VVE.Q-- STTM------ ..H.H..QAV VG.TTNH.-- -.LDK.I.SA
#NcMIC15 ---------- ----RANTPD ELLG..NIPN .ELVTQRR-- DVES-HA--- ..KLL.S.IM SS.S.ASLSF VEMSQ.IKDK
#SnMIC15 ---------- ----DTSAAE H.DGRSNVKG GNER.PQ.RP S.PP------ YLARVVRQRR PDGL.RN--- -NGDLPP.GR
#TgMIC14 ---------- ---------- -QEKREPLV. ...VIP.N-- ASTT------ ..PGR.H.AV VN.MV.ND-- -SLD..INEA
#TgMIC15 ---------- ----RQNVPP EIIG.ASFPN .MLVTPRE-- .AEP-Q---- ..ALL.SETM SS.S.ASLSF VELSQ..QDK

#BbMIC14 MTAPDYADNC EMNIEAKLGH GMMLCKQKAV DCQYSEWYEW SECSLTCKSA -GDGNDSYRI RERRLLSSAE NGG-----VC
#BbMIC15 LISLGFHET. TTPEVLAP.R .IIM..R.LH N.V.T..T.. ....PS.FDW -AE.LVPT.V .S.N.VADDD V--------N
#CcMIC14 RRITKFDVG. ATKGKRLA.R .FVF..A.QK ..I....GA. ....A..R.G P.GMAG.V.L .T.QIVAPST G..EQCRFLT
#CcMIC15 FET.EF..K. T--------- ---.SPE.EP K.EFT..S.. .A...S.IPS A.NVSS.V.. .T.EA.EYGK ---------R
#CsMIC14 LKD.EF.AE. .VGLDRN..G .LVF..YAR. N.EF.P.SD. NT.THS.RNG E..S.....V .T.A.RG..L ...-----.-
#CsMIC15 VASNRFS--- ---------- ---------- ---------- ---------- ---------- ---------- ----------
#EaMIC14 ---------- ---------- ---------- ---------- ---------- ---------- ---------- ----------
#EaMIC15 FQSSEFT.K. A--------- ---FTQEVKP K.EFT..S.. .A.TAS.I.S IAGSGSAM.. .T.DI.EDGK ---------K
#HhMIC14 LS.G.F.E.. .LQSA....K .L.F..RIRT ..E....S.. .D..R..R.G -.GDEE.V.V .S.T..VA.. H..-----S.
#HhMIC15 L.QS.FYSK. SAPEVLAP.E .II...K.VS T.D.T..S.. .T.TA..FDW -DE.VIPL.V .S.DFV.NSP E--------S
#NcMIC14 LSD.KFTA.. .LRSSV...K .L.F..HIRT ..K....S.. .G.....RTG -RNDEE.I.. .D.S..VA.. H..-----H.
#NcMIC15 LNQSNFYS.. STPEVLAP.E .LI...K.VS T.D.T..S.. .T.TA..FDW -DA.SVPL.V .A.E.E..SP E--------T
#SnMIC15 V.V.E.F.QL DG-------- ---MASRYET A.--PP.LS. .G.DAP.IFH P.RPPRR.KA TFSGF.P.EA ---------.
#TgMIC14 LS.G.F.E.. .LQSSV...K .L.F..RIRT ..E....S.. ....R..R.G -..DEE.V.V .G.K..VA.. H..-----S.
#TgMIC15 L.ES.FYSK. SAPEVLAP.E .II...K.VS T.D.T..S.. .T.TA..FNW -DE.VIPL.V .S.DFA..SA D--------S

#BbMIC14 DAERDSSNGQ SVVAIGLCDW LPDCPGVSAQ GDIHI---LM PKEEPKLSPW TPHR--TTTS TRDPDTPLPG QEDTVCTIVN
#BbMIC15 PEQCRNGSR. GT.QTEE.TK V.V.H.SEDV DIEG.PV-VE ..P..VVPA. SDT.ADAGEE VEER.GDHAA TTAVK.FLA.
#CcMIC14 HKTSS.TTDT GT.ELEI.TF QKL.----DD S.DWSTSSIT ..P..S.ED. SADILS.S.T .TE.R----D PGEMI.D..D
#CcMIC15 SFKCRDFSLA GKQLVTT--- ---------- ---------- ---------- ------.KPP SV.RNSIPAD ITPQR..F..
#CsMIC14 -C.S.DKS.S .L.D.Q.... ..L.S.EALP EPDYK---IF .RK..R.AE. .TE.--...T .V...A.P.. S.A.....ID
#CsMIC15 ---------- ---------- ---------- --------.L .RTGAD.PR- ---------- ---------- ----I.AVLR
#EaMIC14 ---------- ---------- ---------- ---------- ---------- ---------- ---------- ----------
#EaMIC15 S.HCSEFSFF GT.ET...PD I.L.NTTEGA TVEL.L--FT T.AP..VE.I KFQTSS..RR PQ.L.A.PAN MTLEQ.S.ET
#HhMIC14 ..DIED.H.G .LSDVQ.... ..P.-DAL.D AEAYV---I. ..P....AE. SAT.--...T .VI.G--.QN ENKIL....D
#HhMIC15 RMSCRLESQN DA.QTEK.N. M.V...AEGE EEDDATGGVE .RG..IVP.. S.E.-P.DEN NQATGSEDIV PGTVE.YVT.
#NcMIC14 ..DIED.Y.W .LNDVQV... .....DAPVD ..SYV---V. ..S..QVGE. S.T.--...T .E..SS-HED EDGIL....D
#NcMIC15 RTSCRVDSEN DL.ETEK.T. M.T..DAEGE DMPDV---.E .RK..TVG.. S.E.-HLDDN NQAMGSDELV .GVVK.YLTD
#SnMIC15 ARISK----- ---HVDD.LD ..L.------ ---------- ---------- ---------- ---------R .I.VS.DQFE
#TgMIC14 ..DIED.H.G .LSDVQ.... ..P.-DAL.D TETYV---I. ..P....AD. SAT.--...T .VI..--.QN ENKIL....D
#TgMIC15 RVLCRLESQN DAIQTEK... M.V..EAEGE EEDDATGGVE .RG..IVP.. S.E.-P.DEN NQAMGSEDIV SGTVE.YVT.

#BbMIC14 MAD-FRTSER GYDTAAESCK CPGKTTVCSR GEASNSREKW ESIMESICGR NGAGQIFAQG MEEFNCGSRS FQSF--SGIF
#BbMIC15 .S.LLYEAV. ...E.YRA.. ..QGRKP.T. A..MATLDN. TKDV.AA.E. GRS.A.VLMD GDR.F.ATG. .-----GKEQ
#CcMIC14 ..T--DSRKM ...K.YRT.. ..SY.RP.LA D..EA..SL. NQT.MVL.KD .KTAR.PLAS FIA.D.ET.T .FDA-HADLN
#CcMIC15 .DA--EGATK E..ISTD..S ..NGYRP.F. ...LLAGQN. QTDAYKL.AS LSN.T.G.RN F.QYS.VF.G .EKDLTVDAA
#CsMIC14 .R.-.KKA.. ...AKHK..R ..SR.RL.A. ...M....N. QTHV.T..A. DSQAD.Y..E L.K.S.A..E .VAV--V.S.
#CsMIC15 V.P------- ---------R L.ES.----- ---------- ---------- GKKLLFLPTR TKT.SWET.- ----------
#EaMIC14 ---------- ---------- ---------- ---------- ---------- ---------- ---------- ----------
#EaMIC15 VDV--SRTLS A..LS.D..S ..HGYRP.Y. K..FL.VDN. INEAYAL.AA TSSAI.GTRH FY..S.VH.G .VDSSAFSSS
#HhMIC14 ...-.SK.H. ....ET.... ..YNAR.... T..V...DN. DEL.QTV.ES ..Q.E.L... ..T.S.SE.V .R..--N.TL
#HhMIC15 .GTIMTSYY. ..NQEYHG.N ...GRRP.T. A..VA.LDL. AKDSGAL.DQ GM.TM.S.AA G.A.F.VTG. .-----GK.D
#NcMIC14 ...-.SD.H. ...PET...T ..YN.KM... T..A...DN. .DL.DT..NK .NQ...L.K. ....T.TN.V .R.S--KSTL
#NcMIC15 .STLINSGS. ..NAQYRT.N ..SGRRP... A..LA.LDY. TKET.AL.Q. GMTTV.AVAH SQA.S.VT.. .-----EELD
#SnMIC15 AVT-YTVA.K DFCRE----S .EEVFRQ..- ---------- --------DL LRE.KYM.TS IQK--.FL.I .PETLPHPSK
#TgMIC14 ...-.SK.H. ....ET.... ..YNAR.... T..V...DN. DEL.QTV.ES ..Q...L... ..T.S.SD.V .R..--R.TL
#TgMIC15 .GTIMTSYY. ..NQEYHG.N ...GRRP.T. A..VA.LDL. TKDSGGL.DQ DM.TM.S.AE G.A.F.ATG. .-----GK.D

#BbMIC14 SESPEEDCKS GDLAYIFCKG EGPIQNDTVM THM--TISLI LGIVFGFIFI YWAVQYSLDI QKVLGITGRY VELSNMLEEV
#BbMIC15 APLS.AA.GT SEYD.VL.E. DR.WE-EHTV .RW--IVC.L ..LTL.IAVV LCCL...G.L ..LV.LA.S. PR.VQE.VAL
#CcMIC14 ESTA.TA.LT EQYSSV..VV AEDLESERRQ AAITKILI.L V..ML.VAVV L.FI...V.V .....LR... ...V..TK.-
#CcMIC15 .SVE.KEIFC AKAK.ML.AV SD.GIAAENA .VFL-ITC.C ..AFT.IVVA .ISL...I.L ..LV.LR... ATI.LE.QQL
#CsMIC14 .AN......T DSM..V...D ...L...RTV LQF--IVA.V I..LL.LT.V .C.L...V.. .QA..LA... ....H..Q..
#CsMIC15 ---------- ---------- ---------- ---------- ---------- ---------- ---------- ----------
#EaMIC14 ---------- ---------- ---------- ---------- ---------- ---------- ---------- ----------
#EaMIC15 TSVK.KETYC SK.R....AK NESSEDAEHS LTFV-IA..C ..V.A.LAVA .LSL...I.L ..LV.LR... .AATLE.QQL
#HhMIC14 GQT..AH..G E.AT....EA -.A.D..LIF .EV--MM.M. T.V.M.IAVV ...I...G.V .....LA... ...T.L.H..
#HhMIC15 TSLS.SS.T. SEYQ.VL.E. -H.YEGIANL .TW--V.C.L ...GG.IC.V LSC....S.. ..L..LA.S. PV.VQNVT.L
#NcMIC14 .QT...H... S.AT.L...D -.A.DSGLIF .QL--LMAFF T.V.LA.AVV ...I...G.V .....F.... .....L.N..
#NcMIC15 ISWTDSA.T. SQYE.VL.E. -V.YEGI.TL .TW--L.CVL ..MGG.VC.V LFCL...S.L ..LV.LA.S. PV.VQNVT.L
#SnMIC15 CPLSPAHRRG K-----L.QT .KECD----- ---------- ---------- --CIFAAG-- ---------- ----------
#TgMIC14 GQT..AH... E.AT....ED -.A.D..LIF .EV--MM.I. T.V.M.IAVV ...I...G.V .....LA... ...T.L.H..
#TgMIC15 TSLS.SS.A. SEYQ.VL.E. -H.YEGIANL .TW--V.C.L ..VGG.IC.V LSC....S.. ..L..LA.S. PV.VQNVT.L

#BbMIC14 DKQDEEKGAE GEQQEGDDEF DKLEGEEDGG ENQ-----YE EGADDAEGTY GEGCEQEEAD EQYNAADD-- ----------
#BbMIC15 ETREK.NAKR .PEALDTWSL SSRSASLI.D LGRERAAAFS ARV.SQATEL LL.E.EGRGT ASETQ.ERRD SAVSFMG---
#CcMIC14 ---------- ---------- ---------- ---------- ---------- ---------- ---------- ----------
#CcMIC15 REKRD.EES. E--------- ---ALLLNSD DKHTEPDVTP DSVPAY..DE NDSAPVRRNS LRNLQ----- ----------
#CsMIC14 EEKEN.AQ-- ---------- ---A.SAGDD .KDGEGLPAG R.RGKLL.ER .N.EPE.GGN SELTKEGG-- ----------
#CsMIC15 ---------- ---------- ---------- ---------- ---------- ---------- ---------- ----------
#EaMIC14 ---------- ---------- ---------- ---------- ---------- ---------- ---------- ----------
#EaMIC15 QEMRAGNEES PLLRPFSARQ QRSFSYDLRE HDSADAADAA DA..A.DAAD AADAADALGA ADAVD.AGAA DAADAAGAT-
#HhMIC14 SLEPQAHEE. ---------- ---KA...E. PDE-----DA .PVAAV..NI ELD.NRR.EE .NGEESEED- ----------
#HhMIC15 QERESHSLRR QG-------- ---NISANSE HSNAHAPSLS DSGW.VD.NH SA.SAVP.E. PWQFGDR.NP PLLGTRKYA-
#NcMIC14 SEAAAA.EEQ ---------- ---..RA..E DDEALCEDDA .PVEKVA.EE EMAGNA..EE .MEEDSEG-- ----------
#NcMIC15 QARENHILRR QV-------- ---AIASHED LSEPDSPWHS GT.WTGDVAL PGRSAFL-D. PWRSEE-GAE PLLGIGNSAV
#SnMIC15 ---------- ---------- ---------- ---------- ---------- ---------- ---------- ----------
#TgMIC14 SQEPQAHEE. ---------- ---KA..GE. PDE-----DA .PEAAV..NS EFD.NPRAKE .NGEESEQ-- ----------
#TgMIC15 QERESH.LRR QG-------- ---NISATSE RSDAHALSLS DSGW.VD.NQ SA.SAFP.EE PWQFEDR.EE PLLSTRKYS-

#BbMIC14 ---GSLDDRA PDDAGELDND AEQDGQEEPE GMYGSDEEGY TEEE------ ---------- ---------- ----------
#BbMIC15 ---.RPSS.G FSRPRRRSTQ LGGASRTPRA SLVA.RGL.K LTGLGSGAQS IGSAERSPST SRVFLEDLDT VGSLYSRAVL
#CcMIC14 ---------- ---------- ---------- ---------- ---------- ---------- ---------- ----------
#CcMIC15 ---------- ---------- ---------- ---------- ---------- ---------- ---------- ----------
#CsMIC14 ---------- --NSEV.EES GNSELP.DKR EWRA.T.S.S VN.------- ---------- ---------- ----------
#CsMIC15 ---------- ---------- ---------- ---------- ---------- ---------- ---------- ----------
#EaMIC14 ---------- ---------- ---------- ---------- ---------- ---------- ---------- ----------
#EaMIC15 ---.AG.IAG AA.VANVSDA .AAANASDA. AASSP.IDAA AKK.L----- ---------- ---------- ----------
#HhMIC14 ---------- -------ASY S.S.SDDDED DEVNE.DDDI VS..------ ---------- ---------- ----------
#HhMIC15 ---RNSSSVG IPEPSVIGQT SPT------- ---------- ---------- ---------- ---------- ----------
#NcMIC14 ---------- ---------- ---.ALDDE. DS.EDGDYDD S...------ ---------- ---------- ----------
#NcMIC15 LSPRHAECSG ASQ.PAMNRI -----RTPRA SIVARRSTER RLA.LGSERE ---------- ---------- ----------
#SnMIC15 ---------- ---------- ---------- ---------- ---------- ---------- ---------- ----------
#TgMIC14 ---------- ---------- -----DASDS ESGSD.DDDI VS..------ ---------- ---------- ----------
#TgMIC15 ---RN.GSAG IPEPS.IGQT SPTQQRVPRA SLAAQRSTCQ LSSRLGQANS PEQSLERRSL KLRDSHADVE LQRTLRKEMK

#BbMIC14 ---------- ---------- ---------- ---------- ---------- ---------- ---------- ----------
#BbMIC15 NDESETLTML PHPRRRSEGS RSESIYPSGQ ANIEGRRRRT TLGARASGFL SGASSRKGST GGREGEIKPS QGDARSKDSD
#CcMIC14 ---------- ---------- ---------- ---------- ---------- ---------- ---------- ----------
#CcMIC15 ---------- ---------- ---------- ---------- ---------- ---------- ---------- ----------
#CsMIC14 ---------- ---------- ---------- ---------- ---------- ---------- ---------- ----------
#CsMIC15 ---------- ---------- ---------- ---------- ---------- ---------- ---------- ----------
#EaMIC14 ---------- ---------- ---------- ---------- ---------- ---------- ---------- ----------
#EaMIC15 ---------- ---------- ---------- ---------- ---------- ---------- ---------- ----------
#HhMIC14 ---------- ---------- ---------- ---------- ---------- ---------- ---------- ----------
#HhMIC15 ---------- ---------- ---------- ---------- ---------- ---------- ---------- ----------
#NcMIC14 ---------- ---------- ---------- ---------- ---------- ---------- ---------- ----------
#NcMIC15 ---------- ---------- ---------- ---------- ---------- ---------- ---------- ----------
#SnMIC15 ---------- ---------- ---------- ---------- ---------- ---------- ---------- ----------
#TgMIC14 ---------- ---------- ---------- ---------- ---------- ---------- ---------- ----------
#TgMIC15 GNQVWDKTV- ---------- ---------- ---------- ---------- ---------- ---------- ----------

#BbMIC14 ----
#BbMIC15 DDAK
#CcMIC14 ----
#CcMIC15 ----
#CsMIC14 ----
#CsMIC15 ----
#EaMIC14 ----
#EaMIC15 ----
#HhMIC14 ----
#HhMIC15 ----
#NcMIC14 ----
#NcMIC15 ----
#SnMIC15 ----
#TgMIC14 ----
#TgMIC15 ----
```

**Supplementary Figure 8.** Alignment of MIC14 and MIC15 amino acid sequences used to generate the phylogenetic tree. The amino acid sequences of MIC14 and MIC15 orthologs are identified by the following accession numbers: *Toxoplasma gondii*, MIC14 (AY089992), MIC15 (DQ459408); *Besnoita besnoiti*, MIC14 (XP_029221473), MIC15 (XP_029222264); *Cyclospora cayetanensis*, MIC14 (XP_026191791), MIC15 (XP_026191793); *Cystoisospora suis*, MIC14 (PHJ15367), MIC15, (PHJ22147); *Eimeria acervulina*, MIC14 (XP_013251268), MIC15 (XP_013251270); *Hammondia hammondi*, MIC14, (HHA_218310), MIC15, (HHA_247195); *Neospora caninum*, MIC14 (CEL70510), MIC15 (CEL70693); *Sarcocystis neurona*, MIC15 (SN3_00200915).

**Supplementary Figure 8**
